# Tightly regulated, yet flexible, directional switching mechanism of a rotary motor

**DOI:** 10.1101/642140

**Authors:** Oshri Afanzar, Diana Di Paolo, Miriam Eisenstein, Kohava Levi, Anne Plochowietz, Achillefs N. Kapanidis, Richard Michael Berry, Michael Eisenbach

## Abstract

Biological switches are wide spread in many biological systems. Among them, the switch of the bacterial flagellar motor has generated much interest because it affects a mechanical process rather than a chemical reaction, it controls the direction of rotation of a rotary motor rather than being an on/off switch, and it is exceptionally ultrasensitive. Yet, the molecular mechanism underlying its function has remained unknown. Here we resolved unique features of this mechanism: On the one hand, it is tightly regulated by multiple means, involving three binding sites and two different covalent modifications, with the binding specificity being dictated by the type of covalent modification and by a strict binding sequence. On the other hand, it endows the motor with flexibility as it involves an intermediate stage of brief switches that provides a “go/no go” situation, in which the motor can either proceed to a stable rotation in the new direction or shift back to the original direction. This intermediate stage appears to be a means of the cell to produce angular deflection of swimming while maintaining directional persistence. Furthermore, we show by mathematical modeling that such a switching mechanism can provide ultrasensitivity. This unique combination of tight regulation, flexibility, and ultrasensitivity makes this switching mechanism of special interest.

## Introduction

Switches are vastly known throughout the field of biology, from transcription and expression of genes to controlling processes of signal transduction, cell fate and cell cycle, to mention a few^1-3^. Most of these switches turn processes on and off. An exception is the switch of the bacterial flagellar motor, which controls the motor’s direction of rotation rather than an on/off process^4^. This dissimilarity combined with this switch’s unique properties — controlling a mechanical rather than a chemical process and being exceptionally ultrasensitive^5^, made it a challenging system of investigation. Indeed, in spite of decades of studies, the molecular mechanism underlying switching of the bacterial flagellar motor has remained obscure.

The switch of the flagellar motor is a large complex at the motor’s base, consisting of multiple copies of the proteins FliM, FliN and FliG (Figure 1A). Since chemotaxis of bacteria is achieved by modulating the direction of flagellar rotation, the main control target in chemotaxis is the switch. In bacteria like *Escherichia coli*, the switch shifts the direction of rotation from the default, counterclockwise, to clockwise in response to binding the signaling protein CheY, which shuttles back and forth between the chemotaxis receptor complex and the flagellar motor. The dependence of clockwise generation on CheY concentration is highly cooperative, meaning that the motor is ultrasensitive^5^. The mechanism underlying this ultrasensitivity is unknown and is quite intriguing in view of the observation that CheY binding to the motor is non-cooperative^6,7^.

**Figure 1.**
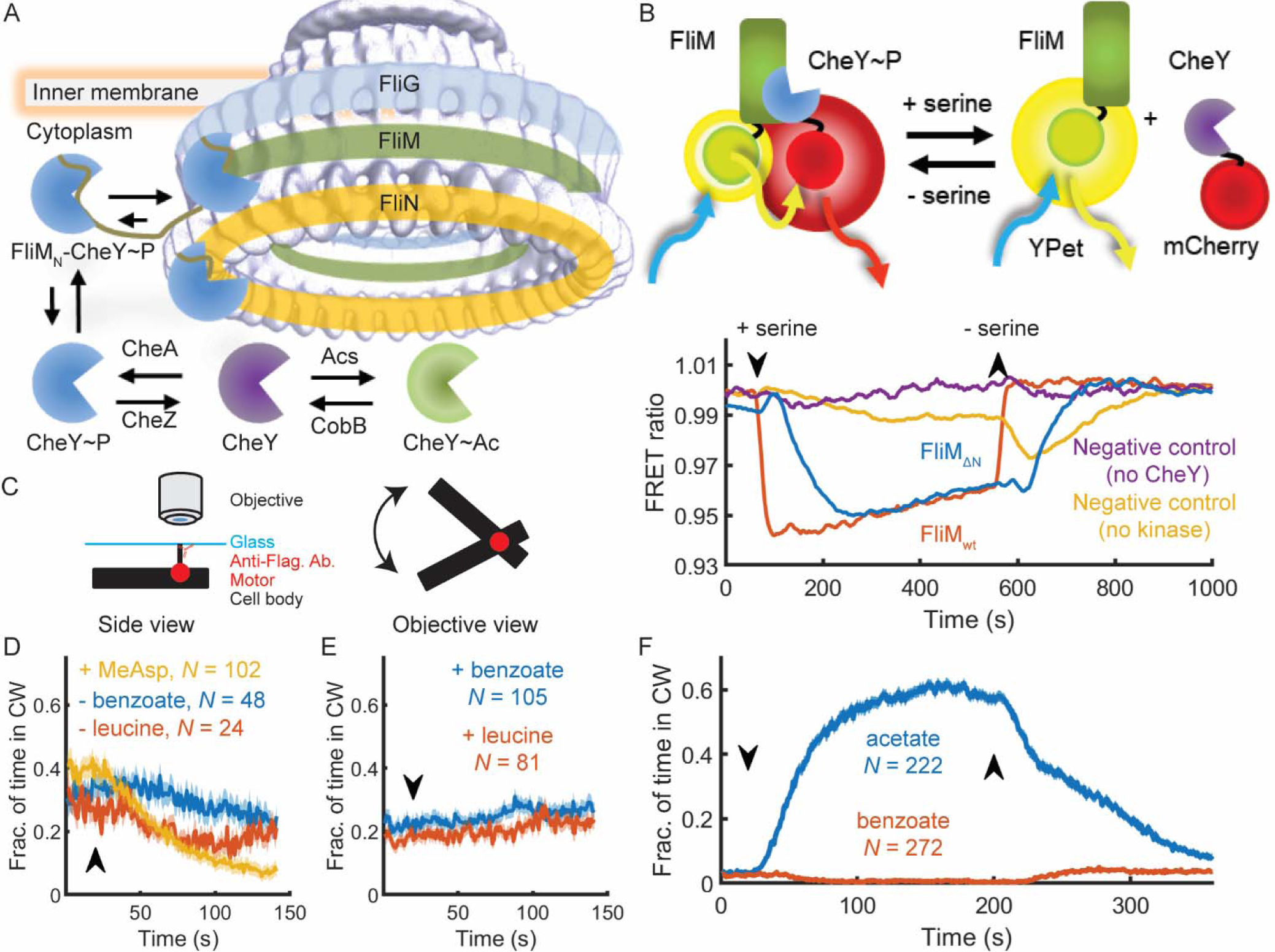
CheY binds to low-affinity sites at the switch to produce clockwise rotation. **A,** CheY interactions in *E. coli.* The electron density map shown here was produced by Thomas *et αl.*^44^ and downloaded from http://dx.doi.org/10.1093/nar/gkv1126. **B**, CheY binds to FliM_ΔN_ motors. **Top**, Experimental scheme of the FRET experiment. The attractant serine lowers the phosphorylation level of CheY. As a result, CheY dissociates from FliM and the energy transfer from YPet (conjugated to FliM) to mCherry (conjugated to CheY) is reduced. **Bottom**, Representative FRET measurements of cells in response to an attractant stimulus (0.1 mM serine). (See Figure S1 for the detailed analysis and Figure S2 for other measurements.) FRET ratio is the ratio of mCherry to YPet fluorescence. CheY-mCherry and mCherry concentrations were ^~^170 μM in the case of FliM_ΔN_ and ^~^15 μM in the case of FliM_wt_ (Figure S3A for calibration of CheY concentration). Data were smoothed with a mean window at the size of 10 data points. Strains used: EW677 (FliM_wt_, red), EW659 (negative control of FliM_ΔN_ without CheY, purple), EW637 (FliM_ΔN_, blue), and EW636 (FliM_ΔN_ Δ*cheA*, yellow). **C**, Experimental scheme for flagellar motor tethering. The stub of sheared flagellum is tethered by antibody to glass^43^ in a flow chamber^45^. The direction of cell body rotation indicates the direction of motor rotation. D, The response of tethered *fliMM*_ΔN_ Δ*cheZ* cells (strain EW635), induced for CheY expression from a plasmid (200 μM IPTG), to positive stimuli. Lines and shaded regions are the mean time spent in clockwise rotation ± SEM. The arrow marks the introduction or removal of stimuli, as indicated. N is the number of cells. MeAsp and leucine were used at 1 mM, benzoate (pH 7.0) at 50 mM. E, As in D for negative stimuli. F, Response of tethered *fliM*_ΔN_ cells (strain sPW416), induced for CheY expression from a plasmid (800 μM IPTG), to acetate and benzoate (both at 50 mM, pH 7.0). Down- and up-arrows mark the introduction and removal of stimuli, respectively. Other details as in D.

Each cell contains multiple flagellar motors. When they rotate counterclockwise the cell swims in a rather straight line. When a considerable fraction of flagella rotate clockwise, the cell preforms a vigor turning motion termed tumbling^8,9^, as a result of which the subsequent linear swimming is in a randomly new direction. It was proposed^10^ and demonstrated^11^ that anisotropic tumbling (in which only some of the motors rotate clockwise for a very short time), does not randomize the swimming direction and maintains directional persistence of swimming. This behavior was proposed to markedly improve the performance of collective migration^11^, implying an evolutionary advantage. Maintenance of directional persistence requires extremely short intervals of clockwise rotation. However, it is unclear how an ultrasensitive motor can produce both relatively long clockwise intervals for tumbling and short intervals for directional persistence.

CheY can be activated by phosphorylation to generate clockwise rotation. This activation is regulated by CheA and CheZ as specific kinase and phosphatase, respectively (Figure 1A). A receptor-mediated attractant response (or removal of a repellent) inhibits CheA activity; stimulation by repellents (or attractant removal) enhances its activity (for reviews —^12-15^). Another covalent modification that activates CheY to generate clockwise rotation is lysine-acetylation^16-18^. The regulation of acetylation is known to involve Acs and CobB as acetyl-transferase and deacetylase, respectively (Figure 1A)^19,20^. It has been shown that CheY acetylation is involved in bacterial chemotaxis^21^ and that it is inversely affected by CheA and CheZ^22^. Yet, the role that it plays in chemotaxis is still obscure.

Earlier studies of the corresponding author’s group revealed that phosphorylated CheY (CheY^~^P) mainly binds to the switch at the *N*-terminus of FliM (FliM_N_)^23,24^. Subsequently, Blair’s group reported on two additional sites with weaker binding of CheY (termed hereafter ‘low-affinity sites’), one at FliN, to which the binding is FliM_N_-dependent and requires that CheY would be phosphorylated^25^, and one at FliM at other location than FliM_N_^26^. On the basis of this information combined with mutational analysis and a structural model, this group further suggested that the interaction of CheY^~^P with FliM_N_ serves to capture CheY^~^P and that switching to clockwise rotation involves the subsequent interaction of CheY^~^P with FliN^25^. NMR analysis in *Thermotoga maritima* by Dahlquist’s group identified the middle domain of FliM (FliM_M_) as a low-affinity binding site for CheY^27^ suggesting that similar binding may also occur in *E. coli.* It is not yet known how CheY binding to these sites affects the process of clockwise generation. Here we addressed the question of the mechanism underlying the switch function. We show that the motor employs three sequential steps of CheY binding to distinct sites at the switch, each with a different outcome. Binding to the first site activates CheY at the switch. Binding to the second site generates transient motor switching and enables binding to the third site, which stabilizes the switched state. We demonstrate by mathematical modeling that such a gating mechanism can provide ultrasensitivity and we show how short clockwise intervals (leading to directional persistence) and long intervals (leading to tumbling) are produced. We also reveal the role of CheY acetylation.

## Results

### CheY binding to the switch’s low-affinity sites generates clockwise rotation

Suspecting that one of the reasons for the insufficient progress in understanding the switch mechanism is the masking effect of the high-affinity binding of CheY to FliM_N_, making it difficult to distinguish between the functions of each of the individual CheY-binding sites at the switch, we investigated the binding and functional interaction of CheY with motors in which FliM had been truncated to remove FliM_N_ (termed hereafter FliM_ΔN_). To study binding, we employed *in vivo* Förster resonance energy transfer (FRET). We measured the interaction between overexpressed CheY-mCherry and FliM_ΔN_-YPet-labeled motors, using a Δ*cheZ* background to ensure that CheY-mCherry was mostly phosphorylated (upper part of Figure 1B for an explanatory scheme; Figure S1 for a detailed representative analysis). Addition and subsequent removal of the attractant serine caused reduction or enhancement, respectively, of the FRET signal, implying an interaction of CheY-mCherry with the low-affinity sites (lower part of Figure 1B, blue). No response was observed in a negative control with overexpressed mCherry (Figure 1B, purple). The amplitude of the response (Figure 1B, blue) was similar to that of non-truncated FliM (termed hereafter FliM_wt_)-YPet motors (Figure 1B, red). However, in the case of FliM_ΔN_ cells, the CheY-mCherry concentration had to be an order of magnitude higher (^~^170 μM vs. ^~^15 μM in FliM_ΔN_ and FliM_wt_ cells, respectively) and the response was slower. This slower response was expected because phosphorylation and dephosphorylation of overproduced CheY requires more time and because the absence of FliM_N_ impairs the on-rate of CheY^~^P binding to the switch. The response of CheY-mCherry under conditions that do not allow its phosphorylation (Δ*cheA* background, i.e., the kinase is missing and the phosphatase is present) was complex and smaller in amplitude (Figure 1B, yellow).

To determine whether CheY^~^P binding to the low-affinity sites is functional, we examined whether it can generate clockwise rotation of the motor. We tethered cells containing FliM_ΔN_-YPet motors and ^~^100 μM CheY in a Δ*cheZ* background (to ensure phosphorylation of CheY) to glass via their flagella (Figure 1C for experimental scheme), and analyzed their direction of rotation with an automated home-made software. The cells responded to positive stimuli (attractant addition or repellent removal) with reduced clockwise rotation (Figure 1D). Repellent stimulation had hardly any effect in this background (Figure 1E). CheY acetylation, stimulated by the acetyl donor acetate, also generated clockwise rotation in FliM_ΔN_ cells (Figure 1F, blue). Although acetate also functions as a repellent, the clockwise generation was the outcome of acetylation because the repellent benzoate, which acts as a repellent by the same mechanism as does acetate^28,29^ but does not serve as an acetyl donor^17,19^, did not generate clockwise rotation (Figure 1F, red). These results suggest that CheY binding to the low-affinity sites is functional in the sense of generating clockwise rotation, and that acetylation is apparently more potent than phosphorylation for this.

### FliM_N_ filters CheY from the cytoplasm and activates it

The finding that FliM_N_ is not essential for clockwise generation raised the question of what its role is. FliM_N_ promotes the interaction of CheY with the switch because, in its absence, CheY must be overexpressed for clockwise generation. Therefore, a reasonable possibility, suggested earlier^25,27^, is that FliM_N_ tethers CheY to the switch, thereby elevating the local concentration of CheY at the low-affinity sites. According to this possibility, FliM_N_ acts as a ‘fishing line’, increasing the rate of CheY association with the clockwise-generating low-affinity sites at the switch. However, this might not be the only function of FliM_N_. It has been found that FliM_N_-bound CheY adopts an intermediate conformation between the active and inactive states^30^ and enhances the rate of CheY phosphorylation by small phosphodonors^31^. This begs the question of whether these observed features serve to increase CheY’s potency to generate clockwise rotation when it is tethered to FliM_N_. To examine this possibility, we studied the motor’s behavior in a FliM_ΔN_ strain that expresses CheY fused to FliM_N_ (FliM_N_-CheY) from a plasmid prepared and employed earlier by Blair’s group^25^. We compared it with the motor’s behavior of the same strain that expresses wild-type CheY instead of FliM_N_-CheY. The expression of both proteins was induced in parallel and under identical conditions to different levels by an IPTG-inducible promoter. We made this comparison in ΔcheZ and Δ*cheA* backgrounds, elevating and lowering CheY phosphorylation, respectively. In a Δ*cheZ* background, cells containing FliM_N_-CheY^~^P spent much more time in clockwise rotation than cells containing CheY^~^P (Figure 2A; red and cyan, respectively). In a Δ*cheA* background, cells expressing FliM_N_-CheY could generate low levels of clockwise rotation whereas cells expressing wild-type CheY could not produce clockwise rotation (Figure 2A; orange and purple, respectively). This implies that CheY can acquire an active clockwise-generating conformation as a result of binding to FliM_N_.

**Figure 2.**
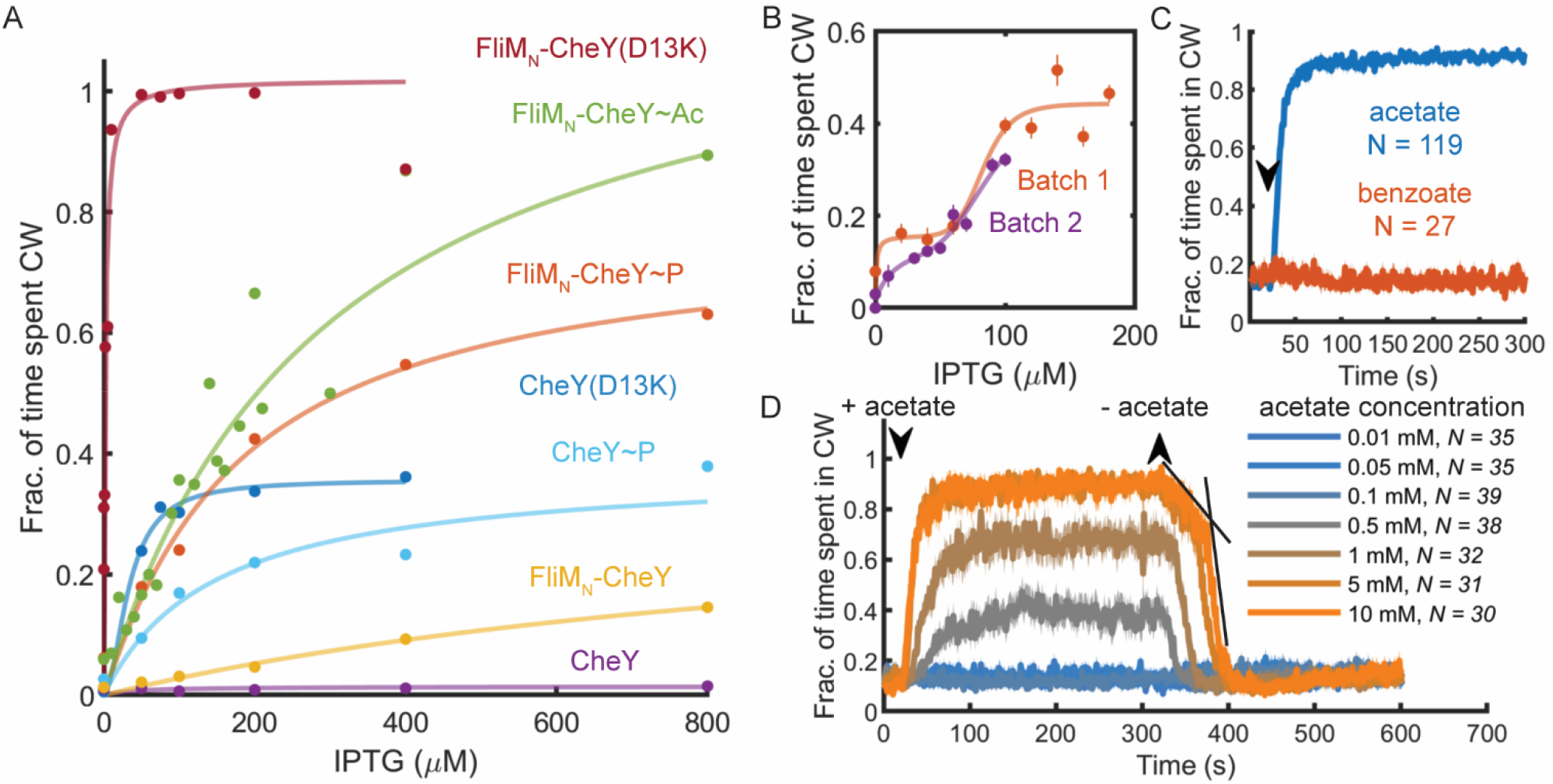
FliM_N_ activates CheY and triggers a biphasic motor response. **A,** Dependence of clockwise rotation on the levels of CheY variants and FliM_N_-CheY variants. The figure shows the mean of clockwise rotation in tethered *fliM_ΔN_ ΔcheZ* cells expressing CheY (cyan) or FliM_N_-CheY (red; strains EW694 and EW697, respectively), *fliM_ΔN_ ΔcheA* cells expressing CheY (purple) or FliM_N_-CheY (yellow; strains EW693 and EW696, respectively), and *fliM_ΔN_ ΔcheA* cells expressing CheY(D13K) (blue) or FliM_N_-CheY(D13K) (dark red; strains EW737 and EW739, respectively). IPTG was used as an inducer. For FliM_N_-CheY^~^Ac (green), acetate (10 mM, pH 7.0) was employed. Each data point is the average of all measurements at a given IPTG concentration, weighted by the sample number of each experiment. For data, see Table S1. B, Dependence of clockwise rotation on the level of FliM_N_-CheY-Ac. The curves, taken from the data of the green curve in A (strain EW696), show two batches (red and purple) in which the clockwise level was determined at several IPTG concentrations. Each data point is a mean ± SEM of the data shown in Table S1 for the second and third batches of this strain (in the presence of acetate)combined. **C,** Comparison of the mean response of *fliM_ΔN_ ΔcheA* cells expressing FliM_N_-CheY (strain EW696) to acetate and benzoate. FliM_N_-CheY expression was induced with 800 μM IPTG. Acetate and benzoate were at 10 mM each (pH 7.0). D, Mean response of *fliM_ΔN_ ΔcheA* cells expressing FliM_N_-CheY (strain EW696) to introduction or removal of varying acetate concentrations (pH 7.0). FliM_N_-CheY expression was induced with 800 μM IPTG. Black lines mark the linear part of the biphasic response to acetate removal. N is the number of cells.

To substantiate the conclusion that FliM_N_ activates CheY independently of phosphorylation, we took advantage of CheY(D13K) being a constitutively active non-phosphorylatable variant of CheY^32^ and compared its dose-dependent clockwise-generating activity with that of FliM_N_-CheY(D13K) under non-phosphorylating conditions, i.e., in a Δ*cheA* background. While the response of CheY(D13K) was similar in magnitude to that of CheY^~^P (Figure 2A; blue), the FliM_N_ fusion extremely increased the clockwise generation potency of CheY(D13K) (Figure 2A; dark red), so much so that clockwise rotation could be generated just by the leaky expression of the protein from the plasmid (levels that were roughly equivalent to endogenous CheY expression levels). Thus, FliM_N_ fusion to CheY(D13K) substantially compensated for the FliM_N_ deletion from the motor. This strongly supports the notion that FliM_N_ functions to promote switching in a phosphorylation-independent manner.

### Motor response to CheY is biphasic

Acetylation, mediated by saturating concentrations of acetate, greatly enhanced clockwise rotation of FliM_N_-CheY in FliM_ΔN_ motors (Figure 2A; green). In batches in which we measured the dependence of clockwise rotation on FliM_N_–CheY-Ac concentration at high resolution, we noticed a biphasic dependence (Figure 2B). The clockwise-generating effect of acetate was acetylation specific as benzoate had no effect on the clockwise rotation (Figure 2C). The finding of biphasic dependence of clockwise rotation on the level of FliM_N_-CheY-Ac (Figure 2B) led us to investigate whether the relaxation of the motor from clockwise to counterclockwise rotation follows a biphasic trajectory. This is because the relaxation only depends on the dissociation constant of CheY from the motor, making it much easier to determine. (At equilibrium, the motor’s occupancy with CheY would be rather constant due to Rate_on_ being equal to Rate_off_, with Rate_on_ = k_on_[CheY], Rate_off_ = k_off_, and k_on_ and k_off_ being the rate constants for the binding and dissociation, respectively.) To this end, we measured the motor’s response to acetate removal in *fliM*_ΔN_ Δ*cheA* cells expressing FliM_N_–CheY. The response to acetate removal was monophasic at acetate concentrations of 0.5 and 1 mM (Figure 2D, gray and brown), with fitted exponential decay rates (k_off_) of −0.085 and −0.099 s^−1^, respectively. However, at higher acetate concentrations, the response became biphasic with a slower response preceding the fast one (Figure 2D, dark- and bright-orange). At 5 mM acetate, the decay rates of the fast and slow phases were −0.095 and −0.006 s^−1^, respectively (Figure 2D, black straight lines). Similar values were measured for 10 mM acetate (−0.091 and −0.005 s^−1^, respectively). The similarity of decay rates for the fast reaction in high- and low-acetate concentrations may indicate that they originate from the same process. Noteworthy, the response to acetate removal of cells expressing CheY and FliM_ΔN_ motors in otherwise wild-type background was also biphasic, though the sequence of phases was inversed and the apparent decay rates were seemingly different (Figure 1F). (Similar measurements with phosphorylation-mediated stimuli could not be performed because all repellents examined, excluding acetate, stimulated an attractant-like response — Figure S4.) This difference between the responses of FliM_N_-CheY- and CheY-expressing strains to acetate removal is consistent with the function of FliM_N_ as CheY activator, affecting the binding stability of the latter to the motor. Overall, these observations further argue for a biphasic reaction of the motor to CheY binding.

### CheY binding to the motor is biphasic

Because of the biphasic response of FliM_ΔN_ motors (Figures 1F, 2B, 2D), we examined whether CheY binding to these motors, i.e., to the low-affinity sites, is biphasic as well. To this end, we measured the amount of CheY bound to FliM_ΔN_ motors as a function of the intracellular concentration of CheY. We determined the amount of bound CheY by FRET photobleaching, measuring the increase in fluorescence intensity of the donor, FliM_ΔN_-YPet, due to photobleaching of the acceptor, CheY^~^P-mCherry (Figure 3A, B). This technique was used previously for FRET measurements of chemotactic signaling at the population level^33^. We preformed the measurements *in vivo,* in single cells, using a Δ*cheZ* background to promote CheY phosphorylation (Figure 3C, experimental points, showing one experiment out of three). We could fit a binding curve to each of the measurements (Figure 3C, colored curve) describing the sum of two binding processes, one a saturation process and the other — a cooperative-like process (Figure 3D for fits of individual experiments, Figure 3E-F for breakdown of the fitted curves). It appears that the saturation process was predominant at low CheY concentrations, making about 20–30% of the total binding, and the cooperative-like process was dominant at high concentrations, making about 70–80% of the total binding. It should be noted that it is unlikely that the measured FRET phases were produced by the interaction of free FliM_ΔN_-YPet with CheY-mCherry molecules in the cytoplasm. This is for two reasons. First, no CheY^~^P-mCherry binding to FliM_ΔN_-YPet was detected *in vitro* at these CheY concentrations. And second, we only measured spots of high intensity FliM_ΔN_-YPet signal. Also, the two binding phases were not the outcome of CheY^~^P-mCherry binding to both free FliM_ΔN_-YPet in the cytoplasm and to FliM_ΔN_-YPet within the motor, yielding the non-cooperative and cooperative phases, respectively. This is because clockwise rotation could be observed at CheY concentrations lower than those of the cooperative phase. Had the non-cooperative phase been the outcome of binding to free cytoplasmic FliM_ΔN_-YPet, clockwise rotation could not have been observed. Two CheY-binding phases, as found here, may well give rise to the observed biphasic motor response to CheY (Figures 1F, 2B, 2D). Hypothetically, each of these phases could be associated with a different CheY-binding site at the motor, i.e., FliN or FliM_M_.

**Figure 3.**
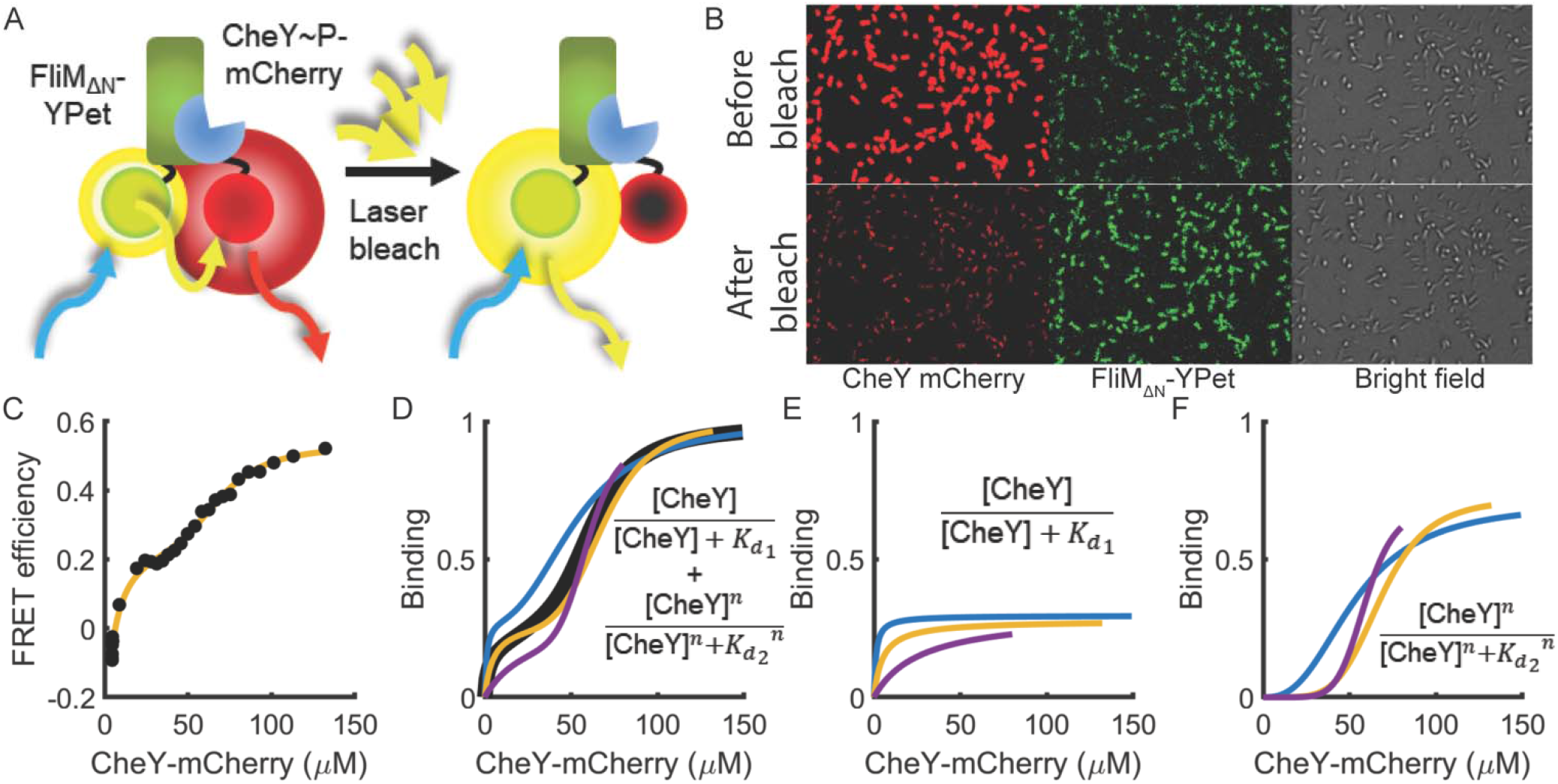
CheY binding to the switch is biphasic. **A,** Experimental scheme: the ratio of YPet fluorescence before and after mCherry photobleaching indicates the extent of CheY bound to FliM_ΔN_. **B,** Demonstration of FRET acquisition, showing the enhancement of FliM_ΔN_-YPet fluorescence due to mCherry photobleaching. **C,** A FRET response curve of *fliM*_ΔN_ Δ*cheZ* cells expressing CheY-mCherry from a plasmid (strain EW637) as a function of the CheY-mCherry concentration. One repetition of the experiment out of three. FRET efficiency was calculated as the ratio of yPET fluorescence before and after photobleaching of mCherry. The data points are the mean ± SEM of 100 measured spots each. Colored curve is a fit to the sum of a saturation- and Hill-binding curves (each multiplied by a proportion factor to account for a different point of saturation for each process). The conversion of mCherry fluorescence intensity to CheY concentration was estimated according to Figure S3B-C. **D,** Fitted curves of three independent measurements, including the one shown in C. Black is the average of the three fits. The average fitted values obtained from the black fitted curve are: *K_d 1_* = 11 ± 8 μM, *K_d 2_* = 60 ± 4 μM and the Hill coefficient *n* = 4.8 ± 1.2 (± SEM). **E,** The saturation-like phase of the fitted curves shown in D. **F,** The sigmoidal (Hill-binding) phase of the fitted curves from D.

### The biphasic response to CheY is reflected in motor kinetics

To determine whether the biphasic motor response to CheY is reflected in the motor’s switching kinetics, we produced survival distributions of clockwise interval length at different levels of clockwise rotation using FliM_N_-CheY-Ac in FliM_ΔN_ motors in a Δ*cheA* background (Figure 4A; intervals processed from the same measurements that were used to construct the green curve of Figure 2A; see Methods and Supplementary Information Methods for analysis details; see Supplementary Movie S1 for a demonstration of a single-cell rotation analysis). The survival distribution describes the probability of a clockwise event to end as time passes. At low clockwise levels, the survival distribution of clockwise intervals appeared as a fast, monophasic exponential decay (Figure 4A, blue; note the logarithmic scale of the ordinate, where exponential decays are seen as straight lines). A monophasic decay suggests the occurrence of a single motor switching process. However, at intermediate clockwise levels and beyond, the distribution became biphasic with the second phase becoming more prominent at increasing clockwise levels (Figure 4A, gray to orange), suggestive of the occurrence of a second process. The distribution of the counterclockwise intervals appeared to mirror the clockwise distributions, being monophasic at high clockwise levels and shifting to biphasic at lower levels (Figure S5A). To better represent the two phases of clockwise rotation, we fitted each of the distributions in Figure 4A with a biexponential expression, and extracted the average length of a clockwise interval of each phase in each clockwise level (see Supplementary Information Methods for details; in cases where the biexponential fitting failed, each part of the distribution was fitted with a monophasic expression). The first phase of clockwise generation linearly increased in length up to saturation, generating only very short intervals of clockwise rotation (Figure 4B, blue). Intervals of the second clockwise-generation phase increased in duration in a nonlinear way, producing relatively long intervals of clockwise rotation (Figure 4B, red). The distribution of rotation rates of these motors was Gaussian-like and centered around −5 Hz and 5 Hz for counterclockwise and clockwise rotation, respectively (Figure 4C). Notably, the average reversal frequency of these motors was roughly 8-fold higher than that of wild-type motors (Figure 4D compared with peak reversal frequency of ^~^0.8 s^−1^ in wild-type cells containing constitutively active CheY^34^), meaning overall shorter clockwise intervals. These results suggest that the two phases of CheY binding to the switch of FliM_ΔN_ motors bring about two phases of clockwise generation. We think that this biphasic response was not detected in earlier measurements^34^ because of the presence of FliM_N_, which convoluted the responses.

**Figure 4.**
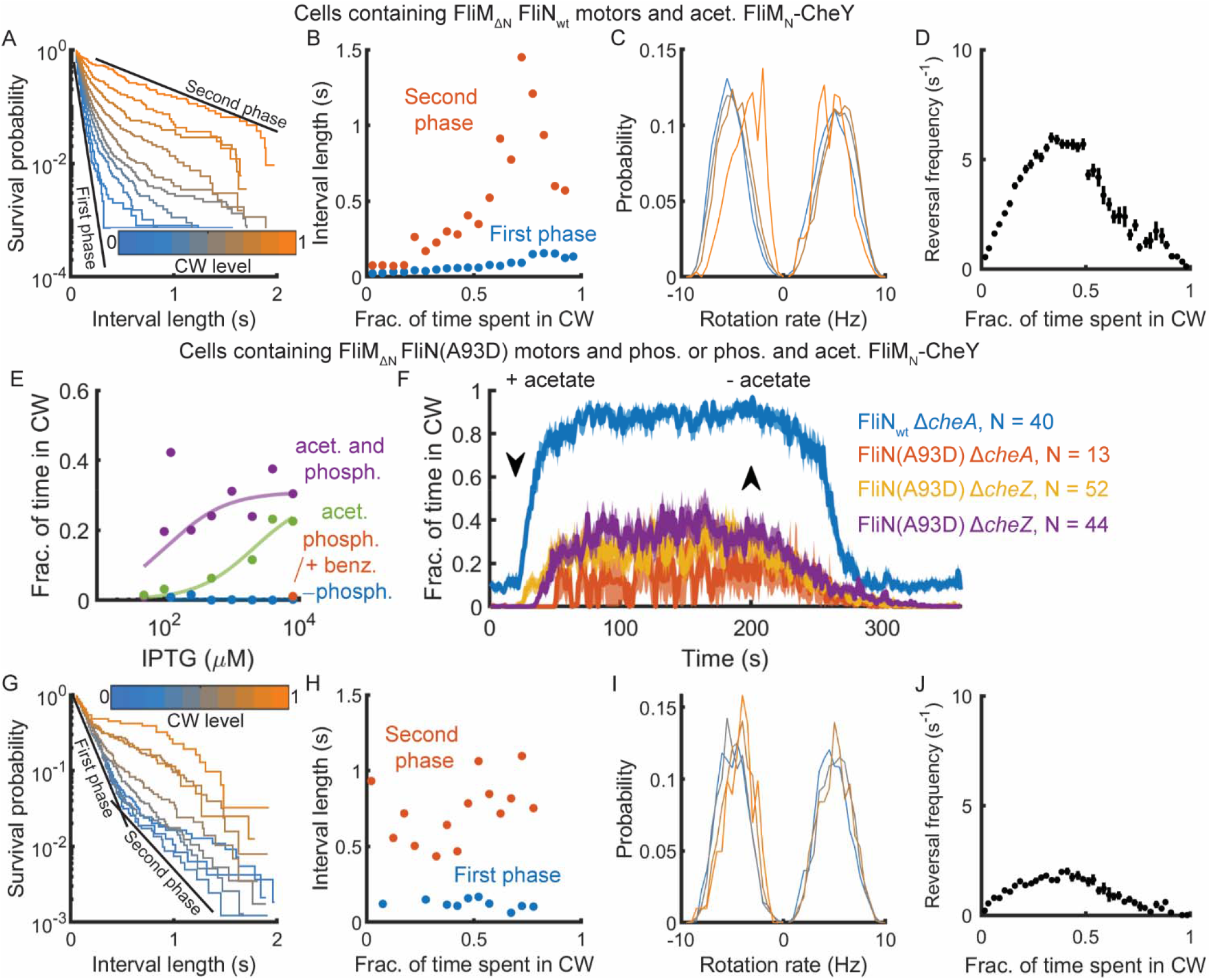
Clockwise generation is a biphasic process with one phase associated with FliN. **A,** Distribution of clockwise interval lengths at different average clockwise levels in Δ*cheA* cells containing FliM_ΔN_ motors and expressing FliM_N_-CheY (strain EW696), in the presence of acetate (10 mM, pH 7.0). Color gradient changes in 0.1 steps from 0 to 1 clockwise rotation. The black lines show the distributions associated with the first and second phases. B, Average interval length of each phase, calculated from the distributions in A. C, Rotation rate for clockwise or counterclockwise intervals (positive or negative, respectively). See A for the color scheme. **D,** Average reversal frequency as a function of the clockwise level of the cells shown in A. See Figure S5 for other kinetics and side-by-side comparisons with similar measurements performed in this study. E, Effects of FliM_N_-CheY acetylation and phosphorylation on clockwise rotation of tethered cells containing FliM_ΔN_ FliN(A93D) motors. Acetate and benzoate concentrations were 10 mM each (pH 7.0). Non-phosphorylated and phosphorylated CheY was produced by using Δ*cheA* and Δ*cheZ* backgrounds (strains EW713 and EW714, respectively). Each data point is the average of all measurements at a given IPTG concentration, weighted by the sample number of each experiment. For data see Table S1. **F,** Response of tethered cells having FliM_ΔN_ FliN(A93D) motors and containing FliM_N_-CheY-Ac and FliM_N_-CheY-P-Ac (red or yellow and purple, respectively) to acetate removal. Tethered cells having FliM_ΔN_ FliN_wt_ motors and containing FliM_N_-CheY (strain EW696) are shown for reference (blue; this is the orange curve in Figure 2D after cutting some time segments before and after acetate removal to synchronize with the other responses). Acetate concentration was 10 mM (pH 7.0). In all strains, IPTG concentration was > 100 μM. The data shown are the mean ± SEM. **G-J,** As in A-D, except that the cells used were Δ*cheZ* containing FliM_ΔN_ FliN(A93D) motors and expressing FliM_N_-CheY (strain EW714).

The measurements above (Figure 4A-D) were based on motors of cells in which acetylation of CheY was promoted. We also obtained similar biphasic behavior when CheY was activated by phosphorylation or by D13K substitution, with or without FliM_N_ fusion (Figure S5B-F), meaning that the biphasic motor response was not the outcome of acetylation or phosphorylation dynamics.

### FliN is associated with the first phase of clockwise generation

The simplest explanation of biphasic binding that results in biphasic clockwise generation is CheY binding to two different sites on FliM_ΔN_ motors. Since, as mentioned above, the known CheY-binding sites in FliM_ΔN_ motors are the low-affinity sites at FliN and FliM_M_, we examined the involvement of each of them. First we determined the involvement of FliN by studying the generation of clockwise rotation in a *fliN*(A93D) mutant, in which the interaction of FliM_N_-CheY^~^P with FliN is greatly impaired^25^. We found that clockwise generation by FliM_N_-CheY^~^P was highly impaired in the *fliN*(A93D) mutant (Figure 4E, blue), supporting the conclusion of Sarkare *et al.*^25^ that FliN is involved in FliM_N_-CheY^~^P binding. Pushing phosphorylation further by the repellent benzoate did not increase clockwise rotation (Figure 4E, red point). Unlike phosphorylation, FliM_N_-CheY acetylation and, the more so, acetylation and phosphorylation together, led to significant clockwise rotation (Figure 4E, green and purple, respectively). Unlike the biphasic response of FliM_ΔN_ FliN_wt_ motors in cells containing FliM_N_-CheY-Ac to acetate removal (Figure 4F, blue), the response of FliM_ΔN_ FliN(A93D) motors in cells containing FliM_N_-CheY-Ac (Figure 4F, red) or both phosphorylated and acetylated FliM_N_-CheY (Figure 4F, purple and orange) to acetate removal was monophasic. The average fitted decay rate of these responses of FliM_ΔN_ FliN(A93D) motors (−0.0135 ± 0.0027 s^−1^, mean ± SD of the 3 measurements in Figure 4F; calculated as an exponential fit to the curve) was closer to that of the slow phase of FliM_ΔN_ FliN_wt_ motors (Figure 4F blue for comparison; this curve is a slight modification of Figure 2D orange, where the fast and slow rates are −0.097 and −0.006 s^−1^, respectively). This suggests that the process, which underlies the slow phase of response to acetate removal, is associated with a CheY binding site other than FliN at the motor. This also suggests that the *fliN*(A93D) mutation impaired the process that enabled the fast decay phase.

To determine whether this impairment is reflected in motor kinetics, we calculated the distribution of the clockwise interval lengths of motors in this mutant under phosphorylating and acetylating conditions (Δ*cheZ* background in the presence of acetate). As in the case of non-mutated FliN, two phases of clockwise generation were observed (Figure 4G). However, the first clockwise-generation phase was saturated already at the lowest clockwise level (Figure 4H, blue) along with the emergence of the second phase (Figure 4H, red). Similar observations were made when the cells lacked CheA (in which case only acetylation occurred; compare Figure S5G with Figure S5H), suggesting that the generation of long intervals were the result of CheY acetylation. The distribution of rotation rates of these motors (Figure 4I) was similar to those of FliN_wt_ (Figure 4C). The occurrence of long intervals could also be demonstrated by the peak reversal frequency of these motors (Figure 4J), which was roughly 3-fold lower than that of wild-type FliM_ΔN_ motors (Figure 4D). This implies that short intervals of clockwise rotation, manifested as frequent reversals, are associated with CheY binding to FliN, and long intervals, manifested as low reversal frequency, is mostly dependent on CheY binding to another site.

### FliM_M_ is associated with the second phase of clockwise generation

The suppression of short intervals of clockwise rotation when CheY interaction with FliN was impaired strongly suggested that long intervals are produced by CheY interaction with a switch site other than FliN. An obvious candidate site for this binding is the other low-affinity binding site, FliM_M_, shown to bind CheY in *T. maritima*^27^. To examine the plausibility of CheY binding to FliM_M_ in *E. coli,* we employed *in vivo* crosslinking of cells expressing FliM_ΔN_-YPet, using a non-specific crosslinker, glutaraldehyde. (We wish to point out that FRET experiments *in vivo* cannot distinguish between CheY binding to FliM and FliN due to being in close physical proximity to each other.) The advantage of using crosslinking is that, beyond being carried out *in vivo*, it can detect weak interactions. We tracked the crosslinking products by SDS-PAGE and scanning the gel for FliM_ΔN_-YPet fluorescence. This enabled us to quantify the extent of complex formation. Among many other crosslinking products, we obtained a product at the size of a complex between FliM_ΔN_-YPet and CheY (Figure 5A, lanes 1–3). The formation of this complex was dependent on CheY overexpression (Figure S6), implying that the complex indeed contained CheY. These observations are consistent with the possibility of CheY binding to FliM_M_ *in vivo*, but they do not rule out a possibility of CheY binding to another FliM domain (e.g., to the C terminus domain of FliM). Various direct *in vitro* binding assays between purified CheY and FliM_ΔN_ were not conclusive, probably due to the low affinity of FliM_ΔN_ for CheY.

**Figure 5.**
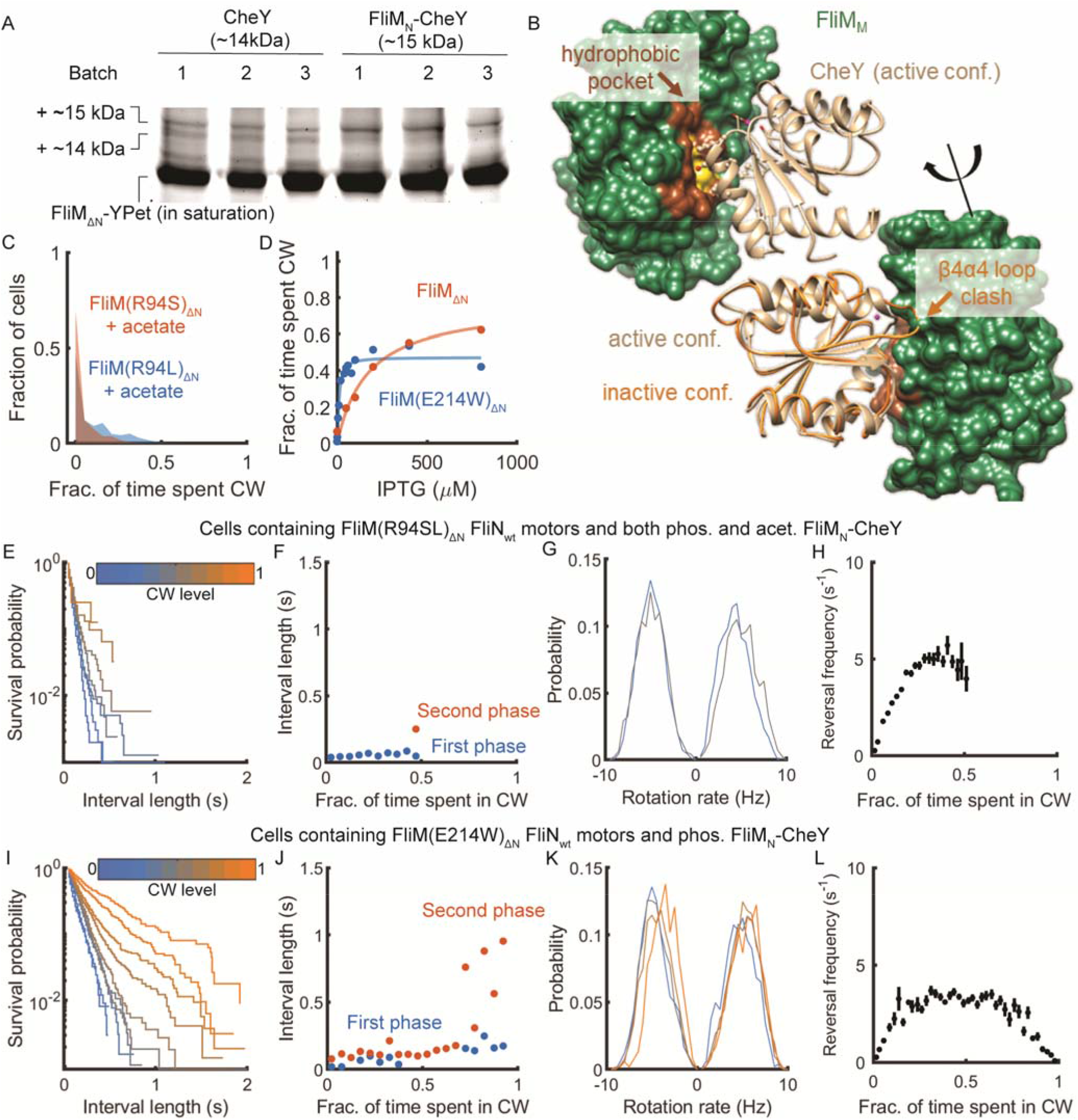
The second switching phase is associated with CheY binding to FliM_M_. **A,** CheY crosslinks with FliM_ΔN_-YPet. The cytoplasm of *flim*_ΔN_-YPet Δ*cheZ* cells overexpressing CheY (strain EW694) or FliM_N_-CheY (strain EW697) from a plasmid was crosslinked by glutaraldehyde and resolved by SDS-PAGE. 1, 2, 3 stand for three different experiments that underwent this procedure. Note that that gel running suffered from parabolic distortion in band positions. The annotations relate to the lowest positions of the bands. The gel was imaged for FliM_ΔN_-YPet fluorescence. To see the crosslinking products, the intense fluorescence of monomeric FliM_ΔN_-YPet is shown at saturation. See Figure S6 for additional details. B, Predicted binding interfaces of CheY (ribbon) and FliM_M_ (green; space filling) in *E. coli*. A binding depression area in FliM_M_ and its surrounding ridge are in yellow and brown, respectively. **Up-left,** The predicted interaction of active CheY (beige) with FliM_M_. **Down-right,** The complex rotated by approximately 180° and aligned with the inactive conformation of CheY (orange), demonstrating that the inactive conformation of the β4α4 loop of CheY clashes with FliM_M_ whereas the active conformation docks well onto FliM_M_. **C,** Clockwise rotation of tethered *fliM(R94S)_ΔN_* Δ*cheZ* cells or *fliM(R94L)_ΔN_* Δ*cheZ* cells (strain EW731 or EW733, respectively) expressing FliM_N_-CheY in the presence of acetate (10 mM, pH 7.0). FliM_N_-CheY expression was induced by 8 mM IPTG. For data see Table S1. **D,** Clockwise rotation of tethered *fliM*(E214W)_ΔN_ Δ*cheZ* cells (strain EW718) at various levels of IPTG for the induction of FliM_N_-CheY expression. Each data point is the average of all measurements at similar IPTG concentrations, weighted by the sample number of each experiment. The red points and curve are taken, as a reference, from Figure 2A. For data see Table S1. **E,** The distribution of clockwise intervals for *fliM(R94SL)_ΔN_* Δ*cheZ* cells expressing FliM_N_-CheY in the presence of acetate (10 mM, pH 7). F, Average interval length in each of the phases as a function of clockwise level for the distributions in E. G, Rotation rate for clockwise or counterclockwise intervals (positive or negative, respectively). See E for the color scheme. H, Average reversal frequency as a function of clockwise level for the cells shown in E. I-L, Same as E-H, except that cells of *fliM*(E214W)_ΔN_ Δ*cheZ* expressing FliM_N_-CheY were used in the absence of acetate. See Figure S5 for other kinetics and side-by-side comparisons with similar measurements performed in this study.

To examine the likelihood that CheY binding to FliM_M_ is functional, we employed docking analysis between CheY and FliM_M_ based on their binding interface in *T. maritima* (Supplementary Methods for detailed description). We especially paid attention to the β4α4 loop of CheY, known to be at the CheY-FliM_M_ binding interface in *T. maritima*^27^ and predicted in our earlier all-atom molecular dynamics simulation to be activated by CheY acetylation at its clockwise-generating site Lys-91^35^. In the docking analysis, we produced a number of models of CheY binding to FliM_M_ in *T. maritima,* and translated those fitted well with the NMR results of Dyer *et al.*^27^ to *E. coli* proteins by superposing the active and inactive forms of CheY upon the model of FliM_M_. We substantiated the superposition results by docking the *E. coli* proteins independently of the *T. maritima* model. This docking demonstrated that while the active conformation of CheY well docked onto FliM_M_ (Figure 5B, upper-left docking model), the inactive conformation of the β4α4 loop of CheY clashed with FliM_M_ (Figure 5B, lower-right). This is consistent with the proposition that acetylation generates clockwise rotation by enhancing the binding of CheY to FliM_M_, and that this enhancement is done by modulation of the β4α4 conformation.

The analysis predicted two prominent latching interfaces of CheY with FliM_M_: An electrostatic interface mostly contributed by arginine at position 94 of FliM (Figure S7A), and a hydrophobic patch in FliM (Figure S7B). To determine the functional relevance of the first interface, we studied motors with substituted arginine at position 94. We examined the rotation of motors of cells expressing FliM_ΔN_(R94S) or FliM_ΔN_(R94L) in a Δ*cheZ* background and overexpressing FliM_N_-CheY. These motors hardly rotated clockwise even when benzoate was used to further stimulate CheY phosphorylation. Very low levels of clockwise rotation were observed following the promotion of CheY acetylation by acetate (Figure 5C). These results are in line with the docking model’s prediction that FliM(R94) is involved in CheY binding. To determine the functional relevance of the second interface, we examined whether replacement of a charged residue with a hydrophobic one in the hydrophobic CheY-FliM_M_ binding interface (E214W) would increase clockwise generation. Indeed, clockwise rotation in the *fliM*_ΔN_(E214W) mutant in a Δ*cheZ* background was observed at much lower FliM_N_-CheY expression levels (Figure 5D). It appears that the mutation affected the motor’s sensitivity to CheY rather than affecting its intrinsic clockwise level because the clockwise rotation levels of *fliM*_ΔN_(E214W) in a Δ*cheA* background, where FliM_N_-CheY is mostly inactive and does not contribute much to clockwise generation, were low and comparable to cells containing non-mutated FliM_ΔN_ (Figure S8; blue). Yet, when FliM_N_-CheY was activated by acetate, the response of cells with FliM_ΔN_(E214W) motors exceeded that of FliM_ΔN_ (Figure S8; red).

To determine whether the proposed interaction of CheY with FliM_M_ affects the second phase of clockwise generation, we examined the distributions of clockwise interval lengths and reversal frequency in *fliM*_δN_(R94SL) and *fliM*_ΔN_(E214W) mutants. In *fliM*_ΔN_(R94SL) mutants, as anticipated, except for the highest observed level of clockwise rotation, only the first phase of clockwise generation was apparent (Figure 5E, F). The rotation rate seemed to be almost independent of the clockwise level (Figure 5G). The peak reversal frequency of the mutants was roughly similar to that of its wild-type FliM_ΔN_ parent but the mutants did not reach high clockwise levels (Figure 5H). In the *fliM*_ΔN_(E214W) mutant, the distributions of clockwise intervals were mostly monophasic and, in some cases, seemingly biphasic (Figure 5I). Fitting of the distributions with monophasic- or biphasic expressions (where applicable) showed that the incline of the second phase was pushed to high clockwise levels (Figure 5J, cf. Figure 4B). The rotation rate of these motors did not vary much with the level of clockwise rotation (Figure 5K). The peak reversal frequency of the FliM_ΔN_(E214W) motors was between the values of wild-type FliM_ΔN_ motors and FliN(A93D) FliM_ΔN_ motors (Figure 5L *versus* Figure 4D, J). Since the apparent monophasic distributions shown in Figure 5I could represent unresolvable mixtures of both phases, we tried segregating between them by fitting the distributions with biphasic exponential expressions. This attempt was unsuccessful. If, indeed, the two phases cannot be resolved in this mutant, it may be that the E214W mutation increased the affinity of FliM_M_ for CheY to a level comparable to that of FliN for CheY. Consequently, intervals initiated by CheY binding to FliN could immediately be extended by subsequent binding to FliM_M_. Furthermore, the inability of the *fliM*_ΔN_(R94SL) mutant to exhibit long clockwise intervals suggests that CheY binding to FliM_M_ is required for stable clockwise rotation. Taken together, these results suggest that the second phase of switching is favored by CheY^~^Ac interaction with FliM_M_, and that the outcome of this interaction is stabilization of the rotation in the clockwise direction.

### CheY^~^P and CheY^~^Ac interactions with the motor result in short- and long-lived interactions, respectively

The observation that the long clockwise intervals produced by FliN(A93D) motors could only be obtained in the presence of acetate (Figure 4E) raised the possibility that CheY^~^Ac is associated with a motor interaction other than FliN, probably FliM_M_ according to the results described above, with a resultant prolonged dwelling of the motor in the clockwise state. To put to the test this possibility and to determine whether prolonged clockwise state can be associated with prolonged CheY^~^Ac dwell at the motor, we measured *in vivo* the dwell time of single CheY molecules at FliM_wt_-YPet motors. We compared between phosphorylating conditions, acetylating conditions, and conditions under which CheY is both phosphorylated and acetylated. Note that these experiments could not be performed with FliM_ΔN_ motors due to the very small encounter rate of CheY with such motors. We electroporated CheY(I95V) molecules labeled with a maleimide modification of the photostable organic dye Atto647 into FliM-YPet expressing cells^36^. The experiments were carried out both in a Δ*cheZ* background to make CheY fully phosphorylated and in a Δ*cheA* background to make CheY non-phosphorylated. The cheY(I95V) mutation was designed to increase CheY affinity for FliM_N_^37^, thus enhancing sampling of otherwise rare binding events. Custom-written software tracked CheY(I95V)-Atto647 molecules and estimated switch locations in each cell using FliM-YPet fluorescence images. CheY(I95V)-Atto647 molecules dwelling within 75 nm of a switch for >30 ms were interpreted as binding to the switch (Figure 6A; see Methods). Both in the absence and presence of acetate, CheY(I95V)-Atto647 molecules were observed to dwell at the switch. Presence of acetate, however, markedly increased the dwell time (Figure 6B *versus* 6C and Figure 6D *versus* 6E for the survival probability, i.e., the probably to remain bound to the switch; Movie S2). Notably, the very long dwell events in Movie S2 were only detected in the presence of acetate. The extension of CheY’s dwell time at the switch by acetylation (Figure 6B–E) taken together with the generation of long clockwise intervals by acetate even when CheY binding to FliN is impaired (Figure 4G) suggest that FliM_M_ has a preference for CheY^~^Ac.

**Figure 6.**
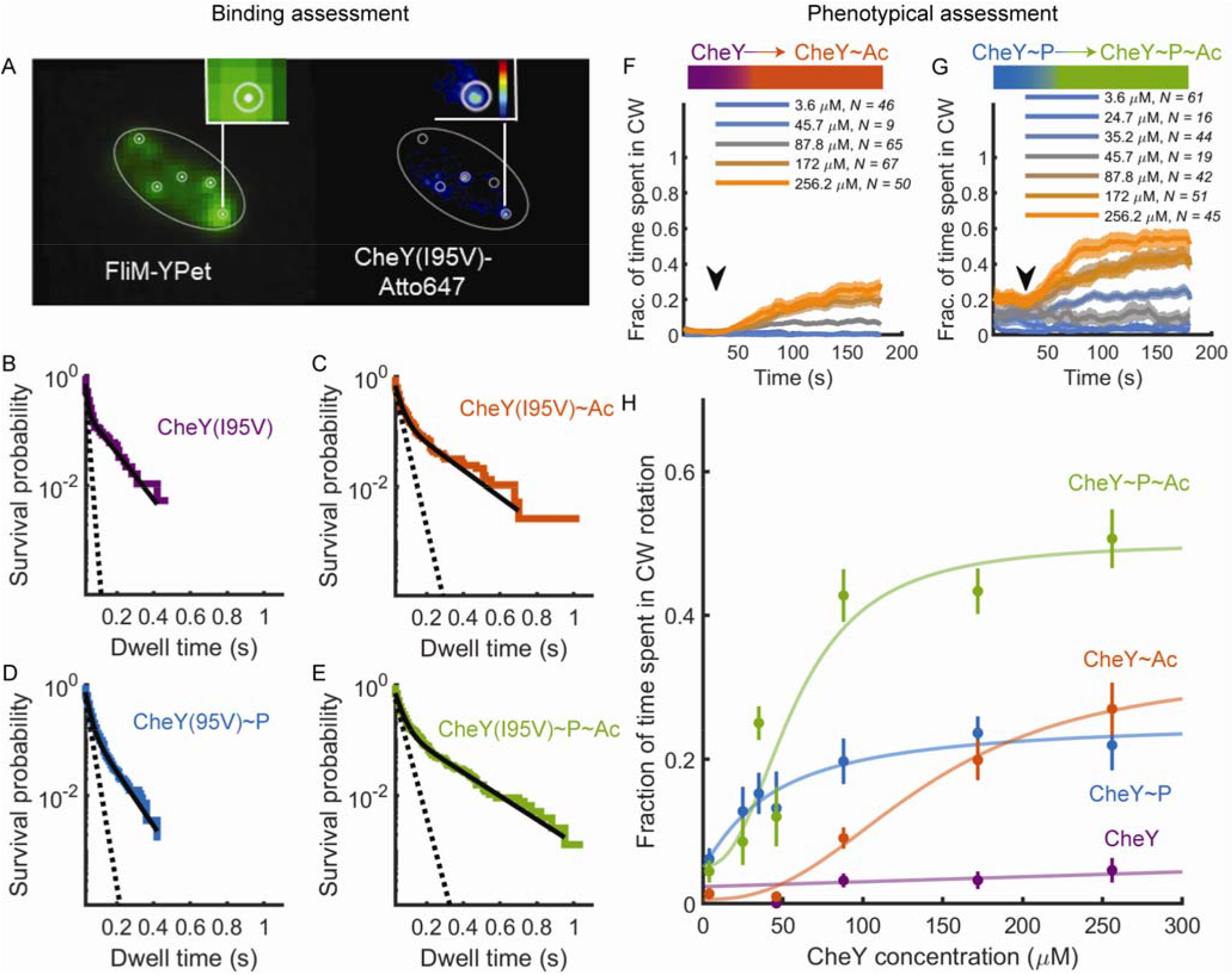
CheY^~^Ac and CheY^~^P differ in their switch-binding kinetics and effective concentration. **A,** Demonstration of single-molecule tracking in a single cell. Cell outline is shown as an ellipse. Left, Imaged FliM-YPet fluorescence. White dots indicate the estimated switch locations. White circles illustrate a 75 nm radius around these locations. Right, Contour map of the normalized sum of probabilities of the localization events of CheY(I95V)-Atto647 (from blue to red is low to high; black is zero probability). White circles show switch locations from the left panel. **B,** Survival probabilities of unmodified CheY(I95V)-Atto647 at the switch (Δ*cheA* background; strain EW668). 63 cells were recorded with a total of 1316 trajectories in which CheY was found to interact with FliM. Note the logarithmic scale of the ordinate. Black line, a bi-exponential fit; dashed line, a fit of the fast decay process (see text for details). **C,** as in B for acetylated CheY(I95V)-Atto647 (presence of acetate, 50 mM, pH 7.0; strain EW668). 56 cells were recorded with a total of 1904 trajectories. **D,** As in B for phosphorylated CheY(I95V)-Atto647 (Δ*cheZ* background; strain EW669). 82 cells were recorded with a total of 2414 trajectories. **E,** as in B for phosphorylated and acetylated CheY(I95V)-Atto647 (presence of acetate, 50 mM, pH 7.0; Δ*cheZ* background, strain EW669). 40 cells were recorded with a total of 2660 trajectories. **F,** Response of tethered *fliM*_ΔN_ Δ*cheA* cells (strain EW634) to acetate (50 mM, pH 7.0) at different CheY concentrations. Lines and shaded regions are mean ± SEM. **G,** Response of tethered *fliM*_ΔN_ Δ*cheZ* cells (strain EW635) to acetate. Details as in F. **H,** Contribution of acetylation and phosphorylation to clockwise rotation. The points are the mean clockwise rotation calculated from F and G at time segments 0-20 s and 160-180 s.

The survival distributions in all experiments appeared to be biphasic, comprised of fast and slow exponential decay processes (note the logarithmic scale of the ordinate in Figure 6B-E). To determine the effect of phosphorylation and acetylation on each of the phases of dwell time, we fitted each of the distributions with a bi-exponential expression [*A_1_*e^(−k_1_t)^ + *A*_2_e^(−k_2_t)^ with pre-exponential factors A_1_> A_2_ and *k_1_*, *k_2_* being the rate constants of the fast and slow process, respectively; t is time] and calculated the decay rate constant of each process (Figure 6B-E). Unmodified CheY exhibited the fastest decay rate of both processes (*k_1_* and *k_2_* being 99.5 and 9.7 s^−1^, respectively; Figure 6B). Phosphorylation markedly decreased the rate of the first decay process, but did not change the second process (*k_2_* and *k_2_* being 45.3 and 11.1 s^−1^, respectively; Figure 6D). Acetylation alone decreased the decay rate of both processes (*k_2_* and *k_2_* being 32.8 and 5.6 s^−1^, respectively; Figure 6C). Comparable values were obtained by phosphorylation and acetylation combined (*k_1_* and *k_2_* being 28.7 and 4.9 s^−1^, respectively; Figure 6E). Thus, it seems that phosphorylation mostly affects the fast phase of CheY dissociation from the motor and that acetylation likely affects both phases.

The results obtained thus far suggest that the first and second clockwise phases are due to CheY interaction with FliN and FliM_M_, respectively, and that, apparently, CheY^~^P and CheY^~^Ac preferentially bind to the former and latter, respectively. To investigate whether, indeed, CheY^~^P and CheY^~^Ac preferentially promote the first and second phase, respectively, we determined whether each of these covalent modifications promotes a different motor response and whether their effects are additive. To this end, we measured the concentration-dependent response of FliM_ΔN_ motors to CheY^~^P, CheY^~^Ac, and CheY stimulated to have both modifications combined. Unmodified CheY did not produce clockwise rotation (pre-acetate stimulation in Figure 6F; purple curve in Figure 6H). CheY acetylation achieved by acetate addition increased, after a short delay, the fraction of time spent in clockwise rotation (Figure 6F). The extent of the response was sigmoidally dependent on the CheY level (Figure 6H, red). Increasing concentrations of CheY^~^P (CheY in a Δ*cheZ* background) enhanced clockwise rotation up to saturation (Figure 6H, blue). Acetylation of CheY^~^P further enhanced clockwise rotation (Figure 6G; Figure 6H, green). Notably, the clockwise curve of CheY^~^P (Figure 6H, blue) resembled the first binding phase (Figure 3E, blue), including the CheY^~^P concentration required for achieving saturation (around 20 μM). Likewise, the clockwise curve of CheY^~^Ac (Figure 6H, red) resembled the second phase of binding (Figure 3E, red). These resemblances are in-line with the suggested conclusion, made above, that the first phase of clockwise production is more associated with CheY^~^P binding and the second phase is more associated with CheY^~^Ac binding.

### The first phase of clockwise generation enables directional persistence in swimming

The physiological relevance of the second phase of clockwise generation, which generates long clockwise intervals, appears obvious as it leads to tumbling^8,9^, which results in an abrupt change in the swimming direction. However, the physiological relevance of the first phase, which leads to short clockwise intervals, is not at all obvious. To address the question of the physiological contribution of the first phase to the swimming behavior of the cells, we measured the angular shift (i.e., not tumbling behavior; see Figure 7A) in 1 s trajectories of *fliM_ΔN_ΔcheA* cells expressing FliM_N_-CheY from a leaky promoter (strain EW696), in the absence or presence of acetate (Figure 7B, blue and red, respectively). We chose this strain because, in the absence of any other activation, it has limited clockwise-generation activity (Figure 2A, orange) and generates short clockwise intervals only (Figure S5B), suggesting that unmodified FliM_N_-CheY is less likely to activate the second phase of clockwise generation. We used a leaky promoter because even slight overexpression resulted in major or complete tumbling behavior. As a control for noise, we made similar measurements in cells that do not express CheY and, therefore, do not generate clockwise rotation (Figure 7B; green and purple for the absence or presence of acetate, respectively). The experimental distributions were significantly different from the control distributions (*P* = 6×10^−9^ and 4×10^−26^ for the absence and presence of acetate, respectively, according to a two-sample Kolmogorov-Smirnov test). In the absence of CheY, turning angles can be related to noise. The same noise is expected to appear also in the experimental distributions. To de-noise the experimental distributions, we subtracted the FliM_N_-CheY measurements from the Δ*cheY* measurements and found that the subtracted distribution peaked at 20° (Figure 7C). This angle is the angular deflection expected to maintain directional persistence of swimming^38^. It is worth noting that, under some assumptions, a peak distribution of 20° turning angle was calculated to be the product of an average clockwise intervals of 50 ms^38^, which is approximately the average clockwise interval length of the first clockwise-generation phase in our measurements (Figure 4B and Figure S5B). Thus, the first phase of clockwise generation appears to produce angular deflection while maintaining directional persistence.

**Figure 7.**
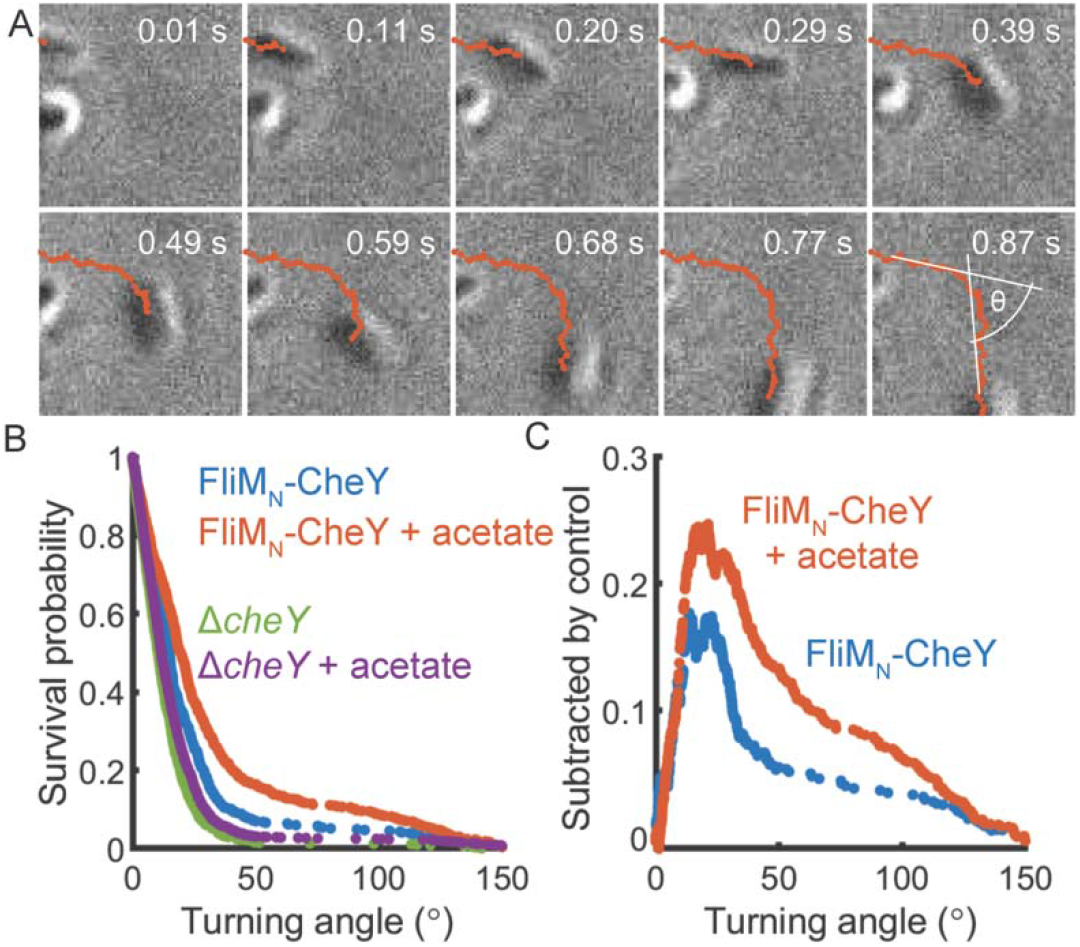
Small deflection angles of swimming in cells that predominantly exhibit the first clockwise-generating phase. A, Montage of a swimming cell, demonstrating deflection measurement. Cell is in black. The trajectory taken by the timestamp is in red. The measured deflection angle is shown in the last frame of the montage. **B,** angle Distributions of angular deflection. Red and blue, *fliM_ΔN_* Δ*cheA* cells expressing FliM_N_-CheY to low levels (strain EW696; leaky promoter expression), in the presence or absence of acetate (10 mM, pH 7), respectively. A total of 1407 and 2135 angles from 32 and 31 different 30 s recordings were collected for the absence and presence of acetate, respectively. Purple and green, Δ*cheY* cells (strain UU1631), in the presence or absence of acetate (10 mM, pH 7), respectively. A total of 1928 and 2977 angles from 22 different 30 s recordings in each case were collected for the absence and presence of acetate, respectively. **C,** Subtracted distributions of A.

## Discussion

Of the many known biological switches, the switch of the bacterial flagellar motor has been a focus of great interest due to its unique properties. It has also been a source of frustration due to lack of success in resolving its molecular mechanism. In the current study we revealed the functions of each of the three CheY-binding sites at the switch, FliM_N_, FliM_M_ and FliN. This led to uncovering processes at the switch that result in brief and long events of switching, their regulation, and physiological significance. All these are discussed below.

### Functions of FliM_N_

We found that the first site to which CheY binds, a high-affinity site at FliM_N_, is not essential for clockwise generation (Figures 1D, 2A). Yet, the order-of-magnitude higher CheY concentration needed for a response in the absence of FliM_N_ (Figure 1D) indicates that FliM_N_ is required for maximal sensitivity of the switch. FliM_N_ does it by tethering CheY to the switch and activating it (Figure 2). To the best of our knowledge, this is the first system in which a ligand (CheY) is activated by its receptor (FliM at the switch), rather than *vice versa.*

The conclusion that FliM_N_ activates CheY is based on our studies with FliM_N_-CheY fusion protein (Figure 2). The published observation that CheY binds to FliM_N_ (*K_d_* = 27 and 680 μM for CheY^~^P and CheY, respectively^39^) strongly suggests that this activation also occurs *in vivo* with non-fused proteins. CheY activation at the switch may be a preliminary step in the process of clockwise generation (Figure 8A), which may have evolved to filter out crosstalk with proteins having CheY-like topology in other two-component signaling pathways. Binding of such proteins to FliM_N_ is expected to be futile.

**Figure 8.**
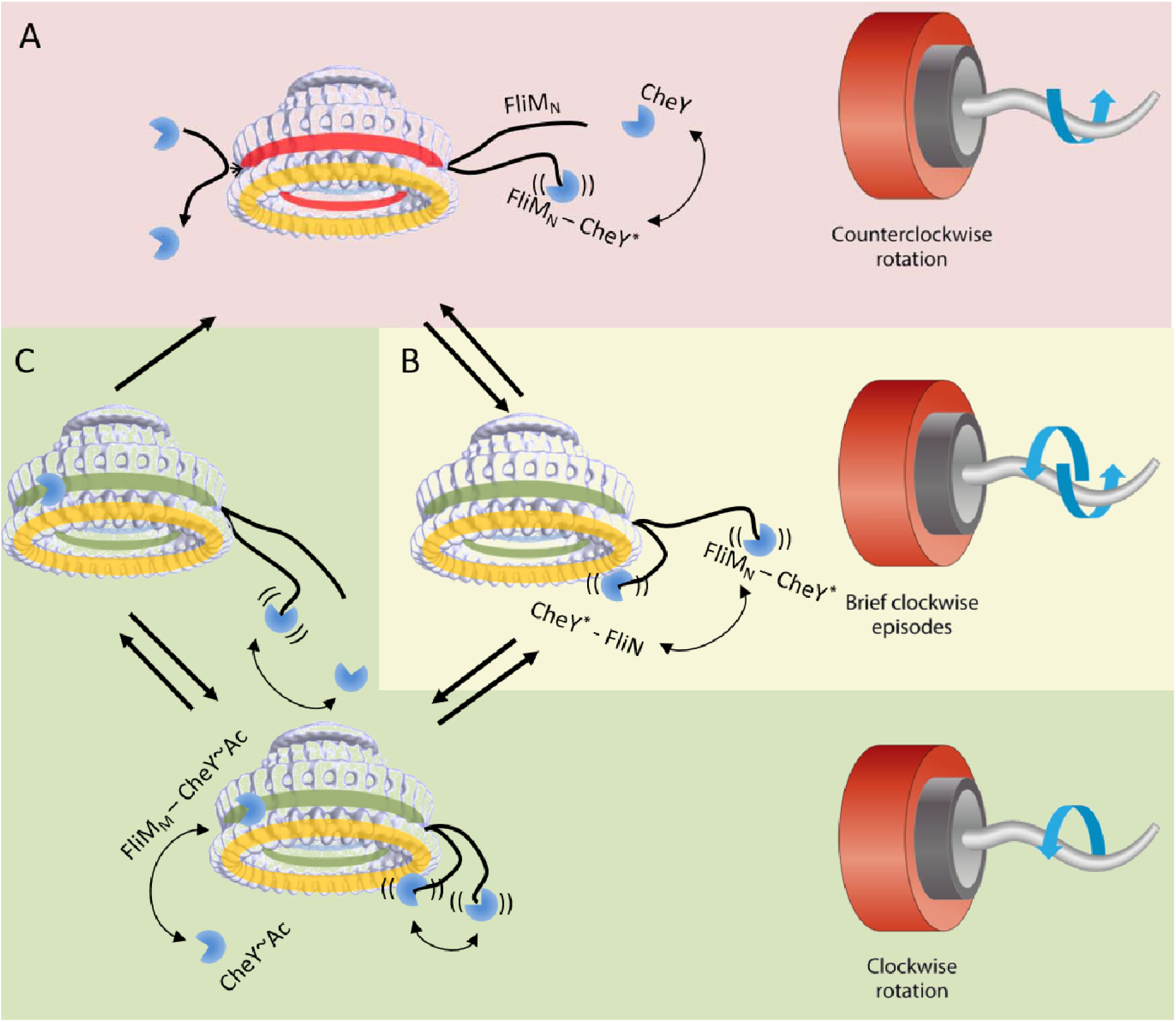
A model for clockwise generation by CheY^~^P and CheY^~^Ac. The model suggests the most likely interactions of CheY at the switch. Yellow ring, FliN. Green or red ring, FliM in a conformation to which CheY can or cannot bind, respectively. Red background region is counterclockwise rotation. Yellow background region is frequent switching. Green shaded region is consecutive clockwise rotation. See text for details.

### The switching mechanism: an apparent gating mechanism

Here we showed that, in FliM_ΔN_ motors, CheY binding to the motor and the motor response to CheY are both biphasic (Figures 3C-F, 4, and 6H). Mutations in FliN and FliM_M_ affected the first and second phases of clockwise generation, respectively (Figures 4–5). On the basis of this it is reasonable to assume that CheY binding to FliN underlies the first phase of binding and clockwise generation, whereas CheY binding to FliM_M_ underlies the second phase of these processes. The generation of solely brief clockwise intervals when CheY can only bind to FliN (Figure 5E), the lack of such intervals when CheY cannot bind to FliN (Figure 4G), and the short dwell time of CheY^~^P at the motor (Figure 6B) suggest that CheY^~^P binding to FliN transiently switches the motor to clockwise rotation. Likewise, the association of long clockwise intervals with FliM_M_ (Figures 4G, 5E) and the relatively long dwell time of CheY^~^Ac at the motor (Figure 6) suggest that CheY^~^Ac preferentially binds to FliM_M_ to produce stable clockwise rotation. We propose that CheY^~^P mainly binds to FliN to make the first catalytic step of clockwise generation following CheY activation by FliM_N_, producing brief episodes of clockwise rotation (Figure 8B), and that CheY^~^Ac mainly binds to FliM_M_ and makes the second catalytic step of clockwise generation, stabilizing the rotation in the clockwise direction (Figure 8C). It is probable that CheY^~^P and CheY^~^Ac can also bind on FliM_M_ and FliN, respectively, though at much lesser affinity. We assume that activated CheY can bind to FliN at any time, but it can bind FliM_M_ only when the motor already rotates clockwise to some extent. This assumption relies on the observation that the orientation of a FliM subunit within the motor is different in the counterclockwise and clockwise states^40^, which may cause FliM_M_ to be sterically blocked for CheY binding in the counterclockwise conformation. Thus, clockwise rotation, which results from CheY binding to FliN, exposes the CheY-binding site at FliM_M_. We term such a mechanism that involves a conditional binding a ‘gating mechanism’.

### Gating can affect ultrasensitivity

The response of wild-type motors to CheY is ultrasensitive^5^. The observation that CheY binding to the switch is non-cooperative^6,7^ suggests that the ultrasensitivity is the outcome of switching steps subsequent to the step of CheY binding to FliM_N_. However, on face value, the current study appears to contradict this suggestion because the ultrasensitive response of the motor was apparently lost when FliM_N_ was removed (Figure 2A). The gating mechanism and the observation of this study that CheY potency to act on the motor is dependent on FliM_N_ and is much reduced in its absence can resolve this apparent inconsistency. Thus, the loss of ultrasensitivity in FliM_ΔN_ motors can be readily explained by differential affinity in the binding of CheY to two motor sites. Conceptually, when CheY binding to the second site is much stronger than to the first site, once clockwise rotation is generated by CheY binding to the first site, the second site would be immediately saturated, producing an ultrasensitive response curve. However, when the affinity of CheY for the first site is much higher, the response curve would break to two saturation curves.

To demonstrate this, we constructed a simplified kinetic model of the switch in which CheY-FliM_M_ binding occurs only when FliN is already CheY-bound. The simplified model consists of a single FliN-binding site and two FliM_M_-binding sites (Figure 9A). When the dissociation constant *(K_d_)* of CheY binding to FliN was much lower than that to FliM_M_, the modeled binding was multiphasic, but not ultrasensitive (Figure 9B, blue). When the *K_d_* of CheY binding to FliN was much higher than the other, the binding was ultrasensitive (Figure 9B, orange). When the *K_d_’s* were comparable, the binding was a simple saturation process (Figure 9B, gray; note the logarithmic scale). Consistently, the apparent Hill coefficients of the curves shifted from positive cooperativity (Figure 9B, insert, orange), through zero cooperativity (Figure 9B, insert; gray), to negative cooperativity (Figure 9B, insert, blue). Thus, the gating mechanism, with some constrains, can explain the gain and loss of the motor’s ultrasensitivity.

**Figure 9.**
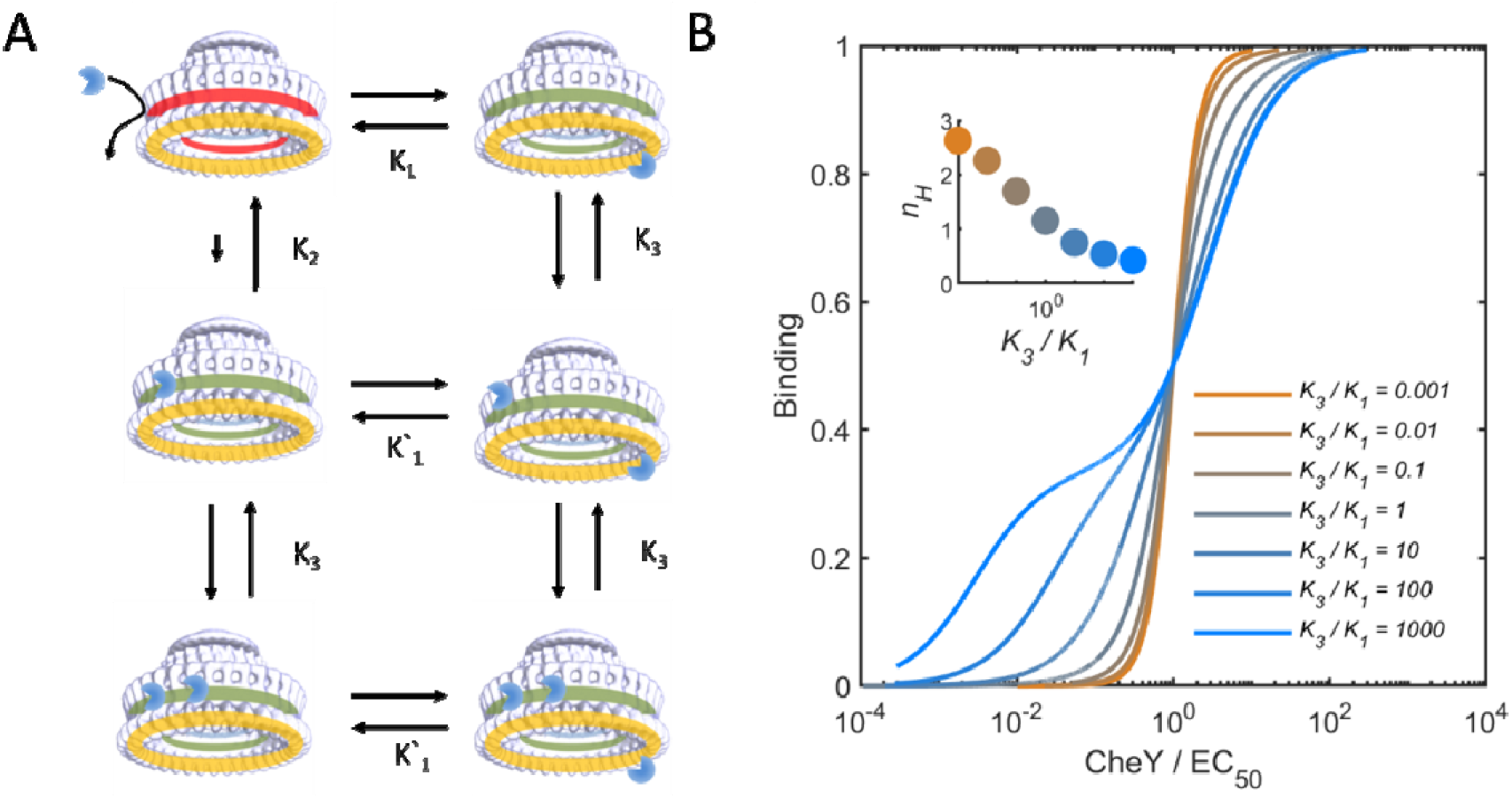
A kinetic model demonstrating varying levels of ultrasensitivity produced by a gating system. **A,** The kinetic model. See text for details. For color codes, see legend to Figure 8. **B,** Binding curve solution of the kinetic model for different *K_3_* values (binding to both FliN and FliM_M_). *K_1_* was set as 1. *K_2_* was set as 10^6^ *K_1_*. Different colors are solutions for different *K_3_* values. Ligand levels were normalized to half-saturation point (EC_50_) of each curve. Note that normalized ligand concentrations are shown in a logarithmic scale. The solution of the model was generated in Mathematica. *Insert,* Calculated apparent Hill-coefficient for the curves in the main panel. Apparent Hill-coefficient was calculated as log_10_(81)/log_10_(EC_90_/EC_10_)^46^.

### Gating supports efficient swimming

When some of the motors in a given bacterial cell exhibit extremely brief episodes of clockwise rotation, the swimming direction is maintained at large, and the outcome is directionally-persistent migration of the cell population. Such directional persistence may markedly improve collective migration. On the other hand, tumbling behavior, which reorients the swimming direction to support the classical run-and-tumble view of bacterial migration^12,13^, requires relatively long clockwise rotation intervals. The gating mechanism can generate both types of swimming behavior because it can produce both short and long clockwise intervals. As discussed above, the very short intervals of clockwise rotation required for directional persistence^11^ can be provided by CheY^~^P binding to FliN (Figure 7) whereas high levels of CheY^~^P, or adequate levels of CheY^~^Ac would produce tumbling by interaction with FliM_M_. This predicts that the bias of the population to choose between directional persistence and tumbling would be very much dependent on the levels of CheY^~^Ac in the cell, which is wired to the cell’s metabolic state. Thus, the gating mechanism apparently optimizes chemotactic performance according to the ambient conditions.

### Unique features of the switching mechanism

The switching mechanism, revealed in this study, has a number of unique features. On the one hand, it is tightly regulated by three distinct binding sites and by two different covalent modifications. Binding to the second site, FliN, depends on phosphorylation of CheY, and binding to the third site, FliM_M_ — on acetylation. This binding can apparently occur only if it is preceded by CheY^~^P binding to FliN. On the other hand, this switching mechanism endows the motor with flexibility, as the intermediate stage at which CheY^~^P is bound to FliN provides a “go/no go” situation, in which the motor can either proceed to a stable clockwise rotation due to binding to FliM_M_ or shift back to counterclockwise rotation. Furthermore, the distinct dependence of each binding step on a different covalent modification suggests that the switching mechanism is wired to inputs from two different signaling pathways: One involves the chemotaxis machinery that regulates CheY phosphorylation, and the other involves the cellular metabolic pathway that regulates CheY acetylation. With this unique combination of seemingly conflicting, but complementing properties of the switching mechanism, it would not be surprising if similar mechanisms are found in the future in the output of other signaling systems.

## Methods

### Strains and plasmids

To produce *fliM_N_* truncation, *fliM-YPet* was cloned from the strain JPA945^41^ (kindly provided by J. Armitage) with genomic flanks of 500 base pairs to the Pst1 sites of pDS132 suicide plasmid^42^. FliM residues 1–16 were truncated from the resulting plasmid by RF-cloning. The constructed plasmid was used to perform allelic exchange with strain JPA945 to produce strain EW566 bearing *flim*Δ(1-16)-*YPet* genomic mutation. The mutation was verified by PCR sequencing. Plasmidic mutations were produced by RF cloning. The genomic deletion mutations of *cheA* and *cheZ* were produced by subjecting cells to P1 transduction with phage containing the genetic background of strain JW1870 or JW1877, respectively. See Tables S2-3 for detailed strains and plasmids list.

### Growth conditions

Strains, diluted 1:100 from overnight cultures, were cultured to mid-late exponential phase at 30°C in tryptone broth with appropriate antibiotics to maintain the plasmids. For CheY expression, cells were grown to mid-late exponential phase and induced by IPTG for 3-4 h. It appeared that dilution of overnight cultures, which spent 1–2 days on the bench at room temperature, produced better responses to acetate.

### Analysis of the direction of flagellar rotation

The direction of flagellar rotation was determined by the tethering assay^43^. Analysis of the rotation was done automatically by an in-house custom-made MATLAB program. For interval analysis, each subjected interval was considered valid if it was flanked by two other valid intervals or by rotation in the opposite direction. See Supplementary Methods for details and other considerations.

### Interval length analysis

We defined clockwise intervals as intervals that have at least 4 consecutive frames (52 ms) of clockwise rotation at a rate higher than 1 Hz, which is the measured rotation rate of fluctuations of non-rotating cells distributed around 0±0.5 Hz (±SD). This 4-frame threshold excluded short events that might not be clockwise rotation mediated by CheY. This is according to a negative control consisting of a Δ*cheY* strain, in which the generation of clockwise rotation cannot be mediated by CheY. In this negative-control strain we studied 1 min recordings of 344 cells, and found ^~^100 clockwise intervals, all shorter than 50 ms. To avoid artefacts that might have been caused by interactions between the cell body and the surface, we also excluded from the analysis intervals that were flanked by pause events. We grouped clockwise intervals from cells that had similar average clockwise levels and produced from each group a distribution of interval lengths.

### In vivo FRET

Cells were immobilized onto a polylysine-coated 8-well glass bottom (ibidi, Germany), washed 4 times with motility buffer and visualized using an inverted confocal microscope set (IX81, Olympus, Japan) with an UPLSAPO 60×0 NA:1.35 objective equipped and excitation lasers at 488 and 559 nm for YPet and mCherry, respectively. FRET was measured by acceptor photobleaching with high intensity 559 nm laser. FRET efficiency was estimated as (I_Post bleaching_ – I_prior bleaching_) / I_post bleaching_. See Supplementary Methods for details.

### Single molecule analysis

The locations of CheY(I95V)-Atto647 and FliM-YPet were automatically estimated by a custom-made MATLAB script. Switch locations were identified as peaks of fluorescence. The exact switch location was determined by identifying the maxima location of a 2D Gaussian, fitted to each switch spot. For the identification of switches, several images were acquired, normalized and averaged before Atto647 imaging. The location of CheY(I95V)-Atto647 was determined similarly to switch location in each frame. CheY(I95V)-Atto647 was considered bound to the switch when it was within 75 nm of a switch location for at least 30 ms (3 frames). Shorter colocalizations were excluded to avoid false positive measurements (e.g., molecules diffusing near the switch). The value of 30 ms was chosen as the half of the expected mean dwell time of CheY(I95V)-Atto647 with FliM_N_ [^~^60 ms given a *K_d_* = 3.9 μM for CheY(I95V)-FliM_N_ interaction and an estimated *k_on_* = 4×10^6^ M^−1^s^−1 6,37^]. The mean-square displacement in 2 dimensions due to free diffusion is <*r*^2^> = *4Dt*. Setting <*r*^2^> = (75 nm)^2^ and *t* = 30 ms gives *D* = 0.047 μm^2^ s^−1^, which is lower than the diffusion coefficient of CheY in the cell. Thus, the falsepositive rate due to diffusing molecules is expected to be negligible. See Supplementary Methods for a detailed description of protein labeling and optical setup.

## Acknowledgments

We are indebted to Dr. David Blair and Paige Wheatley in his group for numerous intensive discussions, for critical reading of earlier versions of the manuscript, for disclosing their unpublished data, and for providing some of the mutants used in this study. We thank Dr. James E. Ferrell for his contribution in the construction of the mathematical model, Dr. Ady Vaknin and Vered Frank in his group for allowing us to measure FRET in their laboratory and for their useful suggestions and technical assistance, Dr. S. Roy Caplan for helpful discussions, Vladimir Kiss for assistance with operating the confocal microscope, Dr. Judith Armitage for supplying bacterial strains, and Yana Gurevich for technical assistance. This study was supported by grant no. 2013197 from the US-Israel Binational Science Foundation and by grant no. 66/14 from the Israel Science Foundation to M.E., by a UK BBSRC grant no. BB/H01795X/1 and an ERC Starter grant no. ERC 261227 to A.N.K., and by a BBSRC grant to R.M.B. D.D.P. was supported by a UK EPSRC DTC studentship. A.P. was supported by a UK EPSRC DTA studentship and the German National Academic Foundation (Studienstiftung) and the Phizackerley Senior Scholarship in Medical Sciences at Balliol College, Oxford. O.A. was supported by an EMBO short term travel fellowship to Oxford.

## Author contribution

O. A. performed and analyzed the experiments, O. A. and M. Eisenbach designed the experiments, interpreted the results, and wrote the manuscript, O. A. and M. Eisenstein performed the docking analysis, K. L. did some of the experiments and provided technical assistance, D. D. P. and O. A. carried out and analyzed the singlemolecule electroporation experiments, A. N. K. provided the TIRF setup used for them, R. M. B. offered insight interpreting the single-molecule results and provided important input for the data analysis of the tethering experiments and for writing the manuscript, and A. P. shared a script that helped with image alignment.

