## Supplementary Information for "Tightly regulated, yet flexible, directional switching mechanism of a rotary motor"

### Supplementary methods

#### Measurement of the direction of flagellar rotation

The direction of flagellar rotation was determined by the tethering assay [1]. Subsequent to flagellar partial truncation by passing the bacterial culture in a syringe several times, the resulting suspension was usually washed three times in motility buffer (10 mM $KP_i$ pH 7.0, 0.1 mM EDTA). The cells were tethered to a coverslip in a flow chamber as described [2] and washed with motility buffer in the flow chamber for roughly 5 min prior to measurement. All chemicals used in behavioral assays were dissolved in the motility buffer. Documentation was done with a uEye digital camera on top of a Zeiss phase-contrast microscope. Recording was typically done at 75 frames/s.

#### Measurement of swimming cells

Cells were gently washed three times in motility buffer (10 mM $KP_i$ pH 7.0, 0.1 mM EDTA). Acetate, when supplemented, was added to the last wash of the cells. Documentation was done at the focal point nearest to glass with a uEye digital camera on top of a Zeiss phase-contrast microscope. Recording was typically done at 75 frames/s. Each field of cells was typically recorded for 30 s.

#### Automated analysis of flagellar motor direction of rotation

All the analyses were done with a pack of Matlab scripts prepared for this study. These are freely available at https://github.com/OshriAfanzar/Afanzar-et-al-2019. All samples of all experiments were analyzed in the exact same way and codes.

*Frame and movie processing*

Each frame was processed by the following scheme: Image binarization → Identification of connected pixels → Ellipse fitting to bodies of connected pixels. Image binarization process was written in Matlab as the following code lines:

```
Pixels = vidFrame(:);

SortedPixels = sort(Pixels);

LinearBase = linspace(min(Pixels),max(Pixels),length(Pixels));

SubtractPixels= SortedPixels - LinearBase;

Tresh = mean(SortedPixels (find(SubtractPixels,min(SubtractPixels))));

vidFrame = (vidFrame – Tresh)>0;
```

Connected pixels were extracted by the ‘bwconncomp’ function of Matlab and ellipse fitting was done using the ‘regionprops’ function of Matlab. Trajectories of rotating cells were composed by identifying ellipses that shared common pixels area for at least 95% of the recording time. The value of 95% was chosen because during the recording there were sometimes events of missing acquisitions (e.g., out of focus events) and we reasoned that we can trust a cell recording only when it could be identified for at least 95% of the recording time. Trajectories in which the rotation was not smooth (e.g., in cases where the rotation paused due to cell body contacting the glass) and, therefore, contained overrepresented rotation angles, were spotted and discarded. For this, the rotation angles of each of the trajectories were binned in a histogram using 10° spacing. A trajectory was trusted if the number of counts in each of the bins was no more than (total number of counts)*2/(number of bins). Thus, in the case of 10° spacing, the number of bins is 36, meaning that if the total number of angles is 1000, when the number of counts in one of the bins exceeded 56 (1000*2/36), the trajectory was discarded. This value was chosen because we observed that it is close to a flat distribution, expected in the case of random angles, but, yet, enables enough deviation from a flat distribution to accommodate empirical rotation data.

*Analysis of rotation*

Angles extracted from the fitted ellipses were used to calculate rotation velocity as the difference in cell angle with respect to the frame rate. To ensure high quality data, events in which the fitted ellipse met the following stringent conditions were marked as erroneous: 1) Major-axis/Minor Axis < 1.25. 2) Major axis < 7 pixels. 3) Major axis < mean(Major axis)-2SD(Major axis). 4) Major axis > mean(Major axis)+2SD(Major axis). 5) Angular displacement was < 360°/frame rate (e.g., 360/75 for 75 frames per second) or > 10*360°/frame rate (e.g., 3600/75 for acquisition frequency of 75 frames per second). When a single erroneous event was flanked by intervals of a different type, the erroneous event was replaced by an event whose identity was determined by the calculated rate of rotation. For example, for clockwise = →, counterclockwise = ← and erroneous interval = E, for the sequence ←←←←E→→→→ which corresponds to the rotation rates (-5)(-5)(-5)(-5)(0.4)(5)(5)(5)(5) Hz, E would be replaced by → because the erroneous interval 0.4 is positive, i.e., with clockwise tendency.

*Calculation of the fraction of time spent in clockwise rotation*

The fraction of time that a motor spent in clockwise rotation was calculated as the sum of clockwise events in a unit time. For steady state rotation, the fraction of time spent in clockwise rotation was calculated for the whole time of acquisition (typically, 30 s) and averaged over all cells (see illustration in Figure S9A below). For time-resolved rotation (as in the response to stimuli), clockwise rotation of single cells was averaged in 1 s intervals (see illustration in Figure S9B).

*Analysis of interval duration*

Intervals were measured only when they were flanked by intervals of the opposite direction, and were not flanked by erroneous intervals. For example, for clockwise = →, counterclockwise = ←, and erroneous interval = E, the sequence of intervals ←←←←→→EE←←→→→←←← would resolve in only one interval of clockwise rotation at the length of 3 frames. When a single erroneous event was flanked by intervals of the same type, the erroneous event was replaced by an event of the same type as the flanks. For example, ←←←←E←←←← would be replaced by ←←←←←←←←←.

*Analysis of reversal frequency*

The number of zero-velocity crossing events per second (after filtering for noise).

#### Automated analysis of cells swimming

The localization of swimming cells was done as described for tethered cells. For every 1 s of recording, the center of mass of the cells was used to construct a trajectory of cell swimming. In a trajectory with a turning point, the scatter of localization points can correspond to two different straight lines, one before and one after the turn. To find the turning point and its angle within the trajectory, the data points of each of the trajectories were fitted with two linear expressions. Trajectories in which one or both fits had $R^2<0.9$ were discarded. Adequate fits were selected as those having the best average $R^2$ score. For each trajectory, the angle of deflection was calculated as the angle between each of the fitted straight lines. As a control for false angles resulting from localization errors or wobbly swimming, we employed a Δ*cheY* strain.

#### Binding measurements *in vivo* employing photobleaching FRET

Cells with genomic mutations for Δ*cheZ* and *fliM*Δ(1-16)-*YPet*, expressing either CheY-mCherry or mCherry from an IPTG-inducible expression plasmid (*E. coli* strains EW575 and EW569, respectively) were grown to mid-late exponential phase, and incubated at various IPTG concentrations for ~6 h. Cells were immobilized onto a polylysine-coated µ-Slide 8-well glass bottom (ibidi, Germany), washed 4 times with motility buffer and visualized using an inverted confocal microscope (IX81, Olympus, Japan) with an UPLSAPO 60x0 NA:1.35 objective equipped with excitation lasers at 488 and 559 nm for YPet and mCherry, respectively. A field of cells was sequentially scanned twice, bleached by high-intensity 559 nm laser, and sequentially scanned again twice more. High-intensity fluorescence regions in the YPet channel were automatically segmented using a Matlab script having a similar principle to that described in the *Frame and movie processing* section above. Mean fluorescence intensity values of the resulting segments were calculated for the YPet channel prior- and post-bleaching. For each segment, FRET efficiency was estimated as $(I_{Post\ bleaching} - I_{Prior\ bleaching}) / I_{Post\ bleaching}$, and the mCherry fluorescence value was defined as the average fluorescence in the two frames preceding bleaching. The FRET efficiency values were sorted to groups of 100 measurements with similar mCherry fluorescence values. Each of the groups was averaged for mCherry levels and FRET efficiency. An illustration of the experimental setup is shown in Figure S10.

#### Binding measurements *in vivo* employing FRET, in response to stimuli

Cells of various genetic background, containing either *fliM*Δ(1-16)-*YPet* or *fliM-YPet* and expressing either CheY-mCherry or mCherry from an IPTG-inducible expression plasmid, were grown to mid-late exponential phase, and incubated at various IPTG concentrations for ~4 h. The cells were washed with motility buffer and were immobilized onto a polylysine-coated round coverslips. Coverslips were mounted onto a flow chamber and measured for FRET in response to stimuli as described [3]. All chemicals for the assays were dissolved in motility buffer.

#### Measurements of CheY expression levels

To assess CheY levels in the cytoplasm, cells expressing CheY-mCherry or mCherry from a plasmid (strains EW575 and EW569, respectively) were grown to mid-late exponential phase and induced by various IPTG concentrations for 4 h or 6 h. The cells were washed three times in NaPi (10 mM, pH 7.6). A sample of the cells was plated in serial dilutions on LB-agar plates to estimate, by colony forming units, the number of cells in the culture. The rest of the sample was sonicated and loaded in 200 µl aliquots to a 96-well plate. In the same plate, a purified CheY-mCherry protein at a known concentration was loaded in different dilutions. The plate was read by a Cytation-5 plate reader (excitation and emission filters were 570 ± 20 nm and 610 ± 20 nm, respectively). The calculation of the cellular concentration of the expressed proteins assumed that the average volume of a single cell is 1 fl.

**Single molecule observations and analysis**

The protein 6xHis-CheY(I95V) was purified by Protino Ni-TED beads and labeled with a maleimide modification of the organic dye Atto647, previously shown to be bright, photo-stable, not hydrophobic and, therefore, highly compatible with observation of single molecules *in vivo* [4]. The labeled protein was separated from the residual unreacted dye by size-exclusion chromatography and was then electroporated into EW669 cells in a low-salinity buffer. The electroporation approach was carried out as described [5]. The electroporated cells were recovered for 5 min in Super Optimal broth with Catabolite repression, washed 4-5 times in motility buffer, and visualized on agar pads on a customized inverted Olympus IX-71 microscope equipped with two lasers, a 637 nm diode laser (Vortran Stradus; Vortran Laser Technology, Sacramento, CA, USA) and a 532 nm DPSS laser (MGL-III-532nm-100mW, CNI) (Figure S11A). Laser light was combined into a single-mode optical fibre (Thorlabs, Newton, NJ, USA) and collimated before focusing on the objective. Highly inclined and laminated optical sheet (HILO) and total internal reflection fluorescence illuminations were achieved by adjusting the position of the focused excitation light on the back focal plane of the objective (UPLSAPO, 100x, NA 1.4, Olympus). Cellular fluorescence was collected through the same objective, filtered to remove excitation light through a long-pass filter (HQ545LP; Chroma, Taoyuan Hsien, Taiwan) and a notch filter (NF02-633S; Semrock, Rochester, NY, USA), and spectrally separated by a dichroic mirror (630DRLP, Omega, Brattleboro, VT, USA). Each channel was imaged onto separate halves of the chip of an EMCCD camera (iXon+, BI-887, Andor, Belfast, UK). The illumination for bright-field images comprised a white-light lamp (IX2-ILL100; Olympus, Shinjuku, Tokyo, Japan) and condenser (IX2-LWUCD; Olympus) attached to the microscope. Movies and images were recorded using manufacturer's software (Andor). All measurements were carried out in continuous wave mode for both green and red lasers. For all the experiments, the exposure time was 10 ms and the intensities used for YPet and Atto647 were 400 nW/μm$^2$ and 1 μW/μm$^2$, respectively. The locations of CheY(I95V)-Atto647 and FliM-YPet were automatically estimated by a custom-made Matlab script. CheY(I95V)-Atto647 was considered as interacting with the motor when it was in a radius <75 nm away from FliM-YPet spots. Dwell times <30 ms were excluded to avoid false positive measurements (e.g., molecules diffusing near the motor). The value of 30 ms (3 frames) was chosen as the half of the expected mean dwell time of CheY(I95V)-Atto647 with FliM$_N$ [~60 ms given a $K_d$ = 3.9 μM for CheY(I95V)-FliM$_N$ interaction and an estimated $k_{on}$ = 4x10$^6$ M$^{-1}$s$^{-1}$) [6,7]]. To account for blinking events and possible errors in localization, we ignored discontinuity of 10 ms (one frame) in trajectories of motor binding. The accumulative recording of all cells (30 s each) in all experiments enabled identification of ~10$^5$ localizations of single CheY(I95V)-Atto647 molecules. Trajectories from the same type of experiments were joined for survival analysis (Figure S11B). The number of cells recorded was 82 (Δ*cheZ* no acetate), 40 (Δ*cheZ* with acetate), 63 (Δ*cheA* no acetate) and 56 (Δ*cheA* with acetate). The number of trajectories in which CheY was found to interact with FliM was 2414 (Δ*cheZ* no acetate), 2660 (Δ*cheZ* with acetate), 1316 (Δ*cheA* no acetate) and 1904 (Δ*cheA* with acetate).

**In silico docking model**

The structure of *E. coli*'s CheY in different activation states has been determined by X-ray crystallography. However, the structure of *E. coli* FliM$_M$ is unavailable. We, therefore, proceeded in two ways: (1) Docking the available structures of CheY and FliM$_M$ from *T. maritima* and superposition of the corresponding *E.*

*coli* proteins, one of which was modeled, on the resultant docking models. (2) Docking of *E. coli* CheY and a model structure of *E. coli* $FliM_M$. The first procedure circumvents the effect of inaccuracies in the modeled structure of *E. coli* $FliM_M$ on the docking results. However, it assumes that the CheY-$FliM_M$ interaction in *E. coli* and *T. maritima* is very similar.

Several structures of *T. maritima* CheY are available (Protein Data Bank [PDB] codes 1TMY, 2TMY, 3TMY and 4TMY). The structure 4TMY was used in the docking computations. The structure of $FliM_M$ from *T. maritima* is available as a free protein and in complex with FliG middle and middle plus C-terminal domains (PDB codes 2HP7, 3SOH, 4FHR and 4QRM). Of these, structure 4FHR is of high resolution and its chain A is complete and includes loop 134-137, which is not resolved in the free protein. The conformation of this loop should not interfere with the binding of CheY because it binds next to $FliG_M$, which does not compete with $FliM_M$ for CheY binding.

The structure of CheY from *E. coli* was studied extensively. We used in the docking the structure of CheY in complex with $Mg^{2+}$ (PDB code 1CHN) and the structure of active CheY with bound $Mg^{2+}$ extracted from the complex with CheZ (PDB code 1KMI). $FliM_M$ from *E. coli* was modeled based on the structures of $FliM_M$ from *Helicobacter pylori* (PDB code 4FQ0), using the program Modeller. The two sequences have 27% identity and 50% similarity.

Docking was executed with the geometric-electrostatic-hydrophobic (GEH) version of MolFit [8]. MolFit performs a 6-dimensional scan in which one molecule is fixed and the other molecule is rotated and translated in steps. For each rotation/translation pose a GEH score is calculated, which reflects the quality of the geometric and chemical complementarity. The molecules are treated as “soft bodies”; hence incorrect conformations of side chains may lower score of the near native docking models but these models are not discarded. The scan is followed with a post-scan filter (P&S filter) in which additional measures are used to evaluate the predicted interface: solvation energy, statistical preference of residues to be in the interface and statistical preference of pairwise interactions. The P&S filter discards a large proportion of false models, yet retains correct and approximately correct models.

Experimental information regarding residues that are likely to be at the CheY/$FliM_M$ interface, was introduced in two ways: (1) In the 6-dimensional scan, positive weights were given to putative interface residues, increasing the scores and elevating the ranking of poses in which these residues are at the interface. (2) In a post scan test that selects poses in which the putative interface region of one molecule makes physical contact with the putative interface region of the other molecule. Experimental data were derived from the NMR spectra for $FliM_M$ and CheY of *T. maritime* [9]. We used UCSF-Chimera to view the results. The docking of $FliM_M$ and CheY from *T. maritima* produced two models of similar quality in terms of the MolFit geometric-electrostatic-hydrophobic score [8]. However, one of the models corresponded significantly better to the NMR results [9]. In this model residues F13, M14, M17, P105 and Q107 of CheY bind in a depression on the surface of $FliM_M$ and interact with the surrounding circular ridge formed by residues S83, D85, E91, S95, M97, R119 and F210. These ridge residues were identified as CheY binding residues in the NMR experiment [9]. CheY is also predicted to interact with the bottom of the depression, which is lined with the side chains of residues F92, V96, E117, L197, T199 and N212. A deeper cavity is located near $FliM_M$ residue V96 where in our model the side chains of CheY F13 and M14 are accommodated. Notably, very large perturbations in the NMR spectra upon binding were observed for $FliM_M$ V96 and CheY M14, which are predicted here to be in direct contact. Analysis of the conservation of the predicted binding surface of $FliM_M$, using ConSurf [10], indicates that positions 83 and 95 are conserved aliphatic-polar, position 91 is conserved negative, positions 96 and 97 are conserved aliphatichydrophobic and F92 and N212 are almost absolutely conserved. The predicted binding surface of CheY is conserved too. Thus, M14 and P105 are highly conserved and Q107 is conserved aliphatic-polar or negatively charged.

### Supplementary Tables

***Table S1. Measured parameters of clockwise rotation***

| Strain | IPTG (μM) | Frac. time spent in CW | STD | Number of cells | Batch | Clockwise distribution (by 0.1 bins) | | | | | | | | | |
|---|---|---|---|---|---|---|---|---|---|---|---|---|---|---|---|
| | | | | | | 0-0.1 | 0.1-0.2 | 0.2-0.3 | 0.3-0.4 | 0.4-0.5 | 0.5-0.6 | 0.6-0.7 | 0.7-0.8 | 0.8-0.9 | 0.9-1 |
| RP437 *fliM*$_{\Delta N}$-*YPet* Δ*cheA* transformed with *cheY*-pTRC99a-FF4 (EW693) | 0 | 0.01 | 0.03 | 67 | 1 | 0.99 | 0.01 | 0.00 | 0.00 | 0.00 | 0.00 | 0.00 | 0.00 | 0.00 | 0.00 |
| | 50 | 0.01 | 0.02 | 29 | 1 | 1.00 | 0.00 | 0.00 | 0.00 | 0.00 | 0.00 | 0.00 | 0.00 | 0.00 | 0.00 |
| | 100 | 0.01 | 0.02 | 96 | 1 | 0.99 | 0.01 | 0.00 | 0.00 | 0.00 | 0.00 | 0.00 | 0.00 | 0.00 | 0.00 |
| | 200 | 0.00 | 0.01 | 64 | 1 | 1.00 | 0.00 | 0.00 | 0.00 | 0.00 | 0.00 | 0.00 | 0.00 | 0.00 | 0.00 |
| | 400 | 0.01 | 0.01 | 98 | 1 | 1.00 | 0.00 | 0.00 | 0.00 | 0.00 | 0.00 | 0.00 | 0.00 | 0.00 | 0.00 |
| | 800 | 0.02 | 0.03 | 303 | 1 | 0.97 | 0.02 | 0.00 | 0.00 | 0.00 | 0.00 | 0.00 | 0.00 | 0.00 | 0.00 |
| | 0 | 0.01 | 0.02 | 177 | 2 | 0.99 | 0.01 | 0.00 | 0.00 | 0.00 | 0.00 | 0.00 | 0.00 | 0.00 | 0.00 |
| | 50 | 0.01 | 0.02 | 152 | 2 | 0.99 | 0.01 | 0.00 | 0.00 | 0.00 | 0.00 | 0.00 | 0.00 | 0.00 | 0.00 |
| | 100 | 0.01 | 0.01 | 11 | 2 | 1.00 | 0.00 | 0.00 | 0.00 | 0.00 | 0.00 | 0.00 | 0.00 | 0.00 | 0.00 |

| | | | | | | | | | | | | | | | |
|---|---|---|---|---|---|---|---|---|---|---|---|---|---|---|---|
| | 200 | 0.01 | 0.03 | 142 | 2 | 0.99 | 0.01 | 0.01 | 0.00 | 0.00 | 0.00 | 0.00 | 0.00 | 0.00 | 0.00 |
| | 400 | 0.01 | 0.03 | 161 | 2 | 0.96 | 0.04 | 0.00 | 0.00 | 0.00 | 0.00 | 0.00 | 0.00 | 0.00 | 0.00 |
| | 800 | 0.01 | 0.03 | 251 | 2 | 0.97 | 0.03 | 0.00 | 0.00 | 0.00 | 0.00 | 0.00 | 0.00 | 0.00 | 0.00 |

| | | | | | | | | | | | | | | | |
|---|---|---|---|---|---|---|---|---|---|---|---|---|---|---|---|
| RP437 *fliM*$_{\Delta N}$-*YPet* Δ*cheZ* transformed with *cheY*-pTRC99a-FF4 (EW694) | 0 | 0.03 | 0.04 | 112 | 1 | 0.91 | 0.08 | 0.01 | 0.00 | 0.00 | 0.00 | 0.00 | 0.00 | 0.00 | 0.00 |
| | 50 | 0.12 | 0.13 | 103 | 1 | 0.59 | 0.13 | 0.16 | 0.11 | 0.02 | 0.00 | 0.00 | 0.00 | 0.00 | 0.00 |
| | 100 | 0.25 | 0.14 | 93 | 1 | 0.20 | 0.13 | 0.29 | 0.23 | 0.13 | 0.02 | 0.00 | 0.00 | 0.00 | 0.00 |
| | 200 | 0.30 | 0.20 | 140 | 1 | 0.25 | 0.09 | 0.15 | 0.15 | 0.21 | 0.11 | 0.01 | 0.02 | 0.00 | 0.01 |
| | 400 | 0.30 | 0.19 | 64 | 1 | 0.19 | 0.17 | 0.16 | 0.14 | 0.22 | 0.09 | 0.02 | 0.00 | 0.00 | 0.02 |
| | 800 | 0.44 | 0.19 | 27 | 1 | 0.07 | 0.07 | 0.04 | 0.07 | 0.41 | 0.15 | 0.15 | 0.00 | 0.04 | 0.00 |
| | 0 | 0.02 | 0.03 | 42 | 2 | 0.95 | 0.05 | 0.00 | 0.00 | 0.00 | 0.00 | 0.00 | 0.00 | 0.00 | 0.00 |
| | 50 | 0.03 | 0.07 | 42 | 2 | 0.90 | 0.02 | 0.05 | 0.02 | 0.00 | 0.00 | 0.00 | 0.00 | 0.00 | 0.00 |
| | 100 | 0.03 | 0.08 | 55 | 2 | 0.89 | 0.04 | 0.04 | 0.04 | 0.00 | 0.00 | 0.00 | 0.00 | 0.00 | 0.00 |
| | 200 | 0.03 | 0.06 | 59 | 2 | 0.88 | 0.08 | 0.03 | 0.00 | 0.00 | 0.00 | 0.00 | 0.00 | 0.00 | 0.00 |
| | 400 | 0.15 | 0.17 | 46 | 2 | 0.54 | 0.20 | 0.09 | 0.09 | 0.04 | 0.02 | 0.00 | 0.00 | 0.02 | 0.00 |
| | 800 | 0.31 | 0.26 | 25 | 2 | 0.20 | 0.24 | 0.12 | 0.12 | 0.20 | 0.04 | 0.00 | 0.00 | 0.00 | 0.08 |

| | | | | | | | | | | | | | | | |
|---|---|---|---|---|---|---|---|---|---|---|---|---|---|---|---|
| RP437 *fliM*$_{\Delta N}$-*YPet* Δ*cheA* transformed with *fliM*$_{N}$-*cheY*-pTRC99a-FF4 (EW696) | 0 | 0.01 | 0.03 | 81 | 1 | 0.98 | 0.02 | 0.00 | 0.00 | 0.00 | 0.00 | 0.00 | 0.00 | 0.00 | 0.00 |
| | 50 | 0.01 | 0.03 | 102 | 1 | 0.98 | 0.01 | 0.01 | 0.00 | 0.00 | 0.00 | 0.00 | 0.00 | 0.00 | 0.00 |
| | 100 | 0.03 | 0.05 | 109 | 1 | 0.95 | 0.03 | 0.01 | 0.00 | 0.01 | 0.00 | 0.00 | 0.00 | 0.00 | 0.00 |
| | 200 | 0.05 | 0.07 | 88 | 1 | 0.82 | 0.11 | 0.07 | 0.00 | 0.00 | 0.00 | 0.00 | 0.00 | 0.00 | 0.00 |
| | 400 | 0.07 | 0.09 | 94 | 1 | 0.77 | 0.14 | 0.04 | 0.05 | 0.00 | 0.00 | 0.00 | 0.00 | 0.00 | 0.00 |
| | 800 | 0.16 | 0.15 | 382 | 1 | 0.46 | 0.23 | 0.17 | 0.08 | 0.04 | 0.02 | 0.01 | 0.01 | 0.00 | 0.00 |
| | 0 | 0.01 | 0.02 | 117 | 2 | 0.99 | 0.01 | 0.00 | 0.00 | 0.00 | 0.00 | 0.00 | 0.00 | 0.00 | 0.00 |
| | 50 | 0.03 | 0.05 | 116 | 2 | 0.92 | 0.06 | 0.01 | 0.01 | 0.00 | 0.00 | 0.00 | 0.00 | 0.00 | 0.00 |
| | 100 | 0.03 | 0.05 | 138 | 2 | 0.93 | 0.07 | 0.00 | 0.01 | 0.00 | 0.00 | 0.00 | 0.00 | 0.00 | 0.00 |
| | 200 | 0.05 | 0.07 | 118 | 2 | 0.82 | 0.14 | 0.01 | 0.03 | 0.00 | 0.00 | 0.00 | 0.00 | 0.00 | 0.00 |
| | 400 | 0.11 | 0.12 | 88 | 2 | 0.63 | 0.17 | 0.11 | 0.05 | 0.05 | 0.00 | 0.00 | 0.00 | 0.00 | 0.00 |
| | 800 | 0.19 | 0.17 | 142 | 2 | 0.40 | 0.15 | 0.14 | 0.18 | 0.08 | 0.03 | 0.01 | 0.00 | 0.01 | 0.00 |
| | 0 | 0.03 | 0.05 | 59 | 3 | 0.95 | 0.03 | 0.00 | 0.02 | 0.00 | 0.00 | 0.00 | 0.00 | 0.00 | 0.00 |
| | 50 | 0.05 | 0.08 | 8 | 3 | 0.75 | 0.13 | 0.13 | 0.00 | 0.00 | 0.00 | 0.00 | 0.00 | 0.00 | 0.00 |
| | 100 | 0.04 | 0.06 | 118 | 3 | 0.91 | 0.07 | 0.01 | 0.02 | 0.00 | 0.00 | 0.00 | 0.00 | 0.00 | 0.00 |
| | 200 | 0.04 | 0.06 | 118 | 3 | 0.86 | 0.12 | 0.02 | 0.01 | 0.00 | 0.00 | 0.00 | 0.00 | 0.00 | 0.00 |
| | 400 | 0.10 | 0.12 | 132 | 3 | 0.67 | 0.19 | 0.07 | 0.05 | 0.02 | 0.00 | 0.00 | 0.01 | 0.00 | 0.00 |
| | 800 | 0.12 | 0.11 | 371 | 3 | 0.54 | 0.27 | 0.12 | 0.05 | 0.01 | 0.01 | 0.00 | 0.00 | 0.00 | 0.00 |

| | | | | | | | | | | | | | | | |
|---|---|---|---|---|---|---|---|---|---|---|---|---|---|---|---|
| RP437 *fliM*$_{\Delta N}$-*YPet* Δ*cheA* transformed with *fliM*$_{N}$-*cheY*-pTRC99a-FF4 (EW696) + 10 mM acetate | 60 | 0.21 | 0.15 | 204 | 1 | 0.28 | 0.23 | 0.24 | 0.16 | 0.05 | 0.01 | 0.01 | 0.00 | 0.00 | 0.00 |
| | 90 | 0.29 | 0.19 | 151 | 1 | 0.17 | 0.19 | 0.19 | 0.23 | 0.15 | 0.03 | 0.01 | 0.02 | 0.01 | 0.01 |
| | 120 | 0.32 | 0.19 | 146 | 1 | 0.16 | 0.12 | 0.15 | 0.29 | 0.16 | 0.05 | 0.03 | 0.01 | 0.02 | 0.01 |
| | 150 | 0.39 | 0.25 | 152 | 1 | 0.14 | 0.10 | 0.14 | 0.16 | 0.21 | 0.11 | 0.04 | 0.03 | 0.02 | 0.06 |
| | 180 | 0.42 | 0.21 | 182 | 1 | 0.05 | 0.11 | 0.12 | 0.18 | 0.24 | 0.14 | 0.05 | 0.05 | 0.02 | 0.04 |
| | 210 | 0.47 | 0.27 | 164 | 1 | 0.09 | 0.06 | 0.10 | 0.16 | 0.16 | 0.16 | 0.09 | 0.03 | 0.04 | 0.12 |
| | 300 | 0.50 | 0.31 | 105 | 1 | 0.13 | 0.07 | 0.09 | 0.12 | 0.12 | 0.14 | 0.08 | 0.03 | 0.01 | 0.21 |
| | 0 | 0.08 | 0.09 | 153 | 2 | 0.67 | 0.23 | 0.07 | 0.02 | 0.01 | 0.00 | 0.00 | 0.00 | 0.00 | 0.00 |
| | 20 | 0.16 | 0.15 | 51 | 2 | 0.47 | 0.12 | 0.24 | 0.10 | 0.06 | 0.02 | 0.00 | 0.00 | 0.00 | 0.00 |
| | 40 | 0.15 | 0.20 | 57 | 2 | 0.60 | 0.14 | 0.11 | 0.09 | 0.04 | 0.00 | 0.00 | 0.00 | 0.02 | 0.02 |
| | 60 | 0.18 | 0.15 | 68 | 2 | 0.43 | 0.16 | 0.19 | 0.10 | 0.10 | 0.01 | 0.00 | 0.00 | 0.00 | 0.00 |
| | 100 | 0.40 | 0.22 | 148 | 2 | 0.08 | 0.07 | 0.15 | 0.25 | 0.22 | 0.10 | 0.01 | 0.03 | 0.05 | 0.03 |
| | 120 | 0.39 | 0.24 | 109 | 2 | 0.10 | 0.11 | 0.18 | 0.17 | 0.14 | 0.14 | 0.05 | 0.03 | 0.03 | 0.06 |
| | 140 | 0.52 | 0.29 | 72 | 2 | 0.08 | 0.04 | 0.07 | 0.13 | 0.25 | 0.13 | 0.03 | 0.04 | 0.06 | 0.18 |
| | 160 | 0.37 | 0.28 | 166 | 2 | 0.24 | 0.09 | 0.10 | 0.12 | 0.17 | 0.10 | 0.04 | 0.03 | 0.04 | 0.07 |
| | 180 | 0.47 | 0.27 | 214 | 2 | 0.09 | 0.07 | 0.08 | 0.17 | 0.23 | 0.09 | 0.06 | 0.04 | 0.06 | 0.11 |
| | 0 | 0.03 | 0.03 | 28 | 3 | 0.96 | 0.04 | 0.00 | 0.00 | 0.00 | 0.00 | 0.00 | 0.00 | 0.00 | 0.00 |
| | 10 | 0.07 | 0.16 | 43 | 3 | 0.81 | 0.09 | 0.05 | 0.02 | 0.00 | 0.00 | 0.00 | 0.00 | 0.00 | 0.02 |

| | | | | | | | | | | | | | | | |
|---|---|---|---|---|---|---|---|---|---|---|---|---|---|---|---|
| (cont.) | 30 | 0.11 | 0.13 | 114 | 3 | 0.65 | 0.18 | 0.10 | 0.04 | 0.03 | 0.01 | 0.00 | 0.00 | 0.01 | 0.00 |
| | 40 | 0.12 | 0.15 | 157 | 3 | 0.61 | 0.17 | 0.08 | 0.08 | 0.03 | 0.02 | 0.01 | 0.01 | 0.00 | 0.00 |
| | 50 | 0.13 | 0.16 | 159 | 3 | 0.58 | 0.19 | 0.13 | 0.04 | 0.03 | 0.00 | 0.01 | 0.00 | 0.01 | 0.01 |
| | 60 | 0.20 | 0.22 | 111 | 3 | 0.41 | 0.24 | 0.14 | 0.08 | 0.05 | 0.01 | 0.01 | 0.03 | 0.01 | 0.03 |
| | 70 | 0.18 | 0.21 | 155 | 3 | 0.46 | 0.21 | 0.10 | 0.13 | 0.03 | 0.01 | 0.01 | 0.01 | 0.01 | 0.03 |
| | 90 | 0.31 | 0.24 | 223 | 3 | 0.21 | 0.16 | 0.19 | 0.13 | 0.13 | 0.06 | 0.04 | 0.02 | 0.01 | 0.04 |
| | 100 | 0.32 | 0.25 | 277 | 3 | 0.23 | 0.17 | 0.10 | 0.16 | 0.12 | 0.08 | 0.05 | 0.03 | 0.02 | 0.04 |
| | 800 | 0.94 | 0.08 | 71 | 4 | 0.00 | 0.00 | 0.00 | 0.00 | 0.00 | 0.00 | 0.03 | 0.03 | 0.17 | 0.77 |
| | 0 | 0.01 | 0.03 | 7 | 5 | 1.00 | 0.00 | 0.00 | 0.00 | 0.00 | 0.00 | 0.00 | 0.00 | 0.00 | 0.00 |
| | 50 | 0.31 | 0.33 | 20 | 5 | 0.35 | 0.10 | 0.15 | 0.10 | 0.05 | 0.05 | 0.00 | 0.10 | 0.00 | 0.10 |
| | 100 | 0.63 | 0.39 | 10 | 5 | 0.20 | 0.00 | 0.00 | 0.10 | 0.10 | 0.00 | 0.10 | 0.00 | 0.10 | 0.40 |
| | 200 | 0.89 | 0.16 | 31 | 5 | 0.00 | 0.00 | 0.00 | 0.03 | 0.00 | 0.06 | 0.03 | 0.10 | 0.16 | 0.61 |
| | 400 | 0.91 | 0.10 | 41 | 5 | 0.00 | 0.00 | 0.00 | 0.00 | 0.02 | 0.00 | 0.02 | 0.07 | 0.22 | 0.66 |
| | 800 | 0.91 | 0.13 | 147 | 5 | 0.00 | 0.01 | 0.01 | 0.01 | 0.00 | 0.01 | 0.03 | 0.03 | 0.20 | 0.70 |
| | 0 | 0.03 | 0.07 | 70 | 6 | 0.89 | 0.03 | 0.07 | 0.01 | 0.00 | 0.00 | 0.00 | 0.00 | 0.00 | 0.00 |
| | 50 | 0.23 | 0.31 | 44 | 6 | 0.50 | 0.16 | 0.09 | 0.02 | 0.05 | 0.05 | 0.00 | 0.02 | 0.02 | 0.09 |
| | 100 | 0.38 | 0.31 | 34 | 6 | 0.32 | 0.03 | 0.06 | 0.09 | 0.24 | 0.06 | 0.03 | 0.06 | 0.03 | 0.09 |
| | 200 | 0.35 | 0.37 | 22 | 6 | 0.45 | 0.05 | 0.00 | 0.09 | 0.00 | 0.05 | 0.09 | 0.09 | 0.14 | 0.05 |
| | 400 | 0.80 | 0.28 | 26 | 6 | 0.08 | 0.00 | 0.00 | 0.00 | 0.12 | 0.00 | 0.00 | 0.04 | 0.27 | 0.50 |

| | | | | | | | | | | | | | | | |
|---|---|---|---|---|---|---|---|---|---|---|---|---|---|---|---|
| | 800 | 0.70 | 0.40 | 25 | 6 | 0.16 | 0.04 | 0.04 | 0.04 | 0.04 | 0.00 | 0.00 | 0.04 | 0.00 | 0.64 |

| | | | | | | | | | | | | | | | |
|---|---|---|---|---|---|---|---|---|---|---|---|---|---|---|---|
| RP437 *fliM*$_{\Delta N}$*-YPet* Δ*cheZ* transformed with *fliM*$_{N}$-*cheY*-pTRC99a-FF4 (EW697) | 0 | 0.16 | 0.11 | 18 | 1 | 0.39 | 0.11 | 0.33 | 0.17 | 0.00 | 0.00 | 0.00 | 0.00 | 0.00 | 0.00 |
| | 50 | 0.21 | 0.18 | 175 | 1 | 0.37 | 0.17 | 0.16 | 0.17 | 0.08 | 0.03 | 0.01 | 0.00 | 0.02 | 0.00 |
| | 100 | 0.27 | 0.23 | 153 | 1 | 0.31 | 0.17 | 0.11 | 0.11 | 0.16 | 0.05 | 0.04 | 0.03 | 0.00 | 0.03 |
| | 200 | 0.42 | 0.25 | 149 | 1 | 0.10 | 0.10 | 0.16 | 0.17 | 0.14 | 0.10 | 0.08 | 0.05 | 0.03 | 0.07 |
| | 400 | 0.52 | 0.24 | 177 | 1 | 0.03 | 0.08 | 0.09 | 0.10 | 0.19 | 0.19 | 0.10 | 0.08 | 0.06 | 0.08 |
| | 800 | 0.63 | 0.24 | 111 | 1 | 0.02 | 0.03 | 0.06 | 0.05 | 0.16 | 0.18 | 0.13 | 0.11 | 0.05 | 0.23 |
| | 0 | 0.07 | 0.09 | 118 | 2 | 0.78 | 0.13 | 0.03 | 0.06 | 0.00 | 0.00 | 0.00 | 0.00 | 0.00 | 0.00 |
| | 50 | 0.19 | 0.16 | 127 | 2 | 0.38 | 0.21 | 0.14 | 0.16 | 0.08 | 0.02 | 0.01 | 0.01 | 0.00 | 0.00 |
| | 100 | 0.25 | 0.19 | 105 | 2 | 0.31 | 0.10 | 0.18 | 0.10 | 0.21 | 0.07 | 0.01 | 0.00 | 0.01 | 0.00 |
| | 200 | 0.52 | 0.24 | 83 | 2 | 0.04 | 0.05 | 0.11 | 0.14 | 0.14 | 0.19 | 0.10 | 0.06 | 0.07 | 0.10 |
| | 400 | 0.65 | 0.27 | 127 | 2 | 0.06 | 0.03 | 0.02 | 0.06 | 0.11 | 0.13 | 0.09 | 0.15 | 0.09 | 0.26 |
| | 800 | 0.65 | 0.27 | 49 | 2 | 0.06 | 0.00 | 0.00 | 0.14 | 0.12 | 0.08 | 0.18 | 0.08 | 0.06 | 0.27 |
| | 0 | 0.02 | 0.05 | 53 | 3 | 0.96 | 0.00 | 0.04 | 0.00 | 0.00 | 0.00 | 0.00 | 0.00 | 0.00 | 0.00 |
| | 50 | 0.05 | 0.08 | 47 | 3 | 0.87 | 0.09 | 0.00 | 0.02 | 0.02 | 0.00 | 0.00 | 0.00 | 0.00 | 0.00 |
| | 100 | 0.10 | 0.12 | 45 | 3 | 0.60 | 0.24 | 0.07 | 0.07 | 0.00 | 0.02 | 0.00 | 0.00 | 0.00 | 0.00 |
| | 200 | 0.25 | 0.24 | 43 | 3 | 0.30 | 0.23 | 0.16 | 0.05 | 0.14 | 0.02 | 0.05 | 0.00 | 0.00 | 0.05 |
| | 400 | 0.36 | 0.26 | 44 | 3 | 0.16 | 0.23 | 0.05 | 0.27 | 0.07 | 0.07 | 0.02 | 0.05 | 0.05 | 0.05 |

| | | | | | | | | | | | | | | | |
|---|---|---|---|---|---|---|---|---|---|---|---|---|---|---|---|
| | 800 | 0.59 | 0.29 | 11 | 3 | 0.00 | 0.00 | 0.18 | 0.09 | 0.18 | 0.09 | 0.09 | 0.00 | 0.18 | 0.18 |

| | | | | | | | | | | | | | | | |
|---|---|---|---|---|---|---|---|---|---|---|---|---|---|---|---|
| RP437 *fliM*$_{\Delta N}$-*YPet* Δ*cheA* transformed with *cheY*(D13K)-pTRC99a-FF4 (EW737) | 50 | 0.28 | 0.13 | 231 | 1 | 0.10 | 0.12 | 0.32 | 0.32 | 0.11 | 0.02 | 0.00 | 0.00 | 0.00 | 0.01 |
| | 75 | 0.36 | 0.15 | 163 | 1 | 0.02 | 0.12 | 0.19 | 0.32 | 0.21 | 0.10 | 0.02 | 0.01 | 0.01 | 0.01 |
| | 100 | 0.39 | 0.19 | 121 | 1 | 0.02 | 0.10 | 0.18 | 0.31 | 0.22 | 0.06 | 0.04 | 0.02 | 0.02 | 0.04 |
| | 200 | 0.52 | 0.21 | 66 | 1 | 0.00 | 0.02 | 0.05 | 0.30 | 0.29 | 0.06 | 0.06 | 0.11 | 0.03 | 0.09 |
| | 400 | 0.48 | 0.22 | 61 | 1 | 0.02 | 0.05 | 0.08 | 0.25 | 0.33 | 0.07 | 0.03 | 0.05 | 0.07 | 0.07 |
| | 0 | 0.00 | 0.01 | 101 | 2 | 1.00 | 0.00 | 0.00 | 0.00 | 0.00 | 0.00 | 0.00 | 0.00 | 0.00 | 0.00 |
| | 200 | 0.06 | 0.07 | 28 | 2 | 0.79 | 0.14 | 0.04 | 0.04 | 0.00 | 0.00 | 0.00 | 0.00 | 0.00 | 0.00 |
| | 400 | 0.22 | 0.13 | 53 | 2 | 0.17 | 0.26 | 0.32 | 0.17 | 0.06 | 0.02 | 0.00 | 0.00 | 0.00 | 0.00 |
| | 0 | 0.00 | 0.00 | 29 | 3 | 1.00 | 0.00 | 0.00 | 0.00 | 0.00 | 0.00 | 0.00 | 0.00 | 0.00 | 0.00 |
| | 50 | 0.15 | 0.12 | 27 | 3 | 0.41 | 0.30 | 0.15 | 0.11 | 0.04 | 0.00 | 0.00 | 0.00 | 0.00 | 0.00 |
| | 75 | 0.25 | 0.19 | 48 | 3 | 0.23 | 0.21 | 0.25 | 0.15 | 0.10 | 0.02 | 0.00 | 0.00 | 0.02 | 0.02 |
| | 100 | 0.21 | 0.16 | 32 | 3 | 0.28 | 0.28 | 0.16 | 0.13 | 0.13 | 0.03 | 0.00 | 0.00 | 0.00 | 0.00 |
| | 200 | 0.27 | 0.18 | 35 | 3 | 0.14 | 0.20 | 0.26 | 0.26 | 0.06 | 0.03 | 0.00 | 0.03 | 0.03 | 0.00 |
| | 400 | 0.27 | 0.17 | 28 | 3 | 0.11 | 0.25 | 0.25 | 0.25 | 0.07 | 0.00 | 0.04 | 0.04 | 0.00 | 0.00 |
| | 0 | 0.01 | 0.04 | 40 | 4 | 0.98 | 0.00 | 0.03 | 0.00 | 0.00 | 0.00 | 0.00 | 0.00 | 0.00 | 0.00 |
| | 50 | 0.14 | 0.12 | 71 | 4 | 0.46 | 0.23 | 0.15 | 0.15 | 0.00 | 0.00 | 0.00 | 0.00 | 0.00 | 0.00 |
| | 75 | 0.22 | 0.13 | 56 | 4 | 0.20 | 0.29 | 0.29 | 0.13 | 0.09 | 0.00 | 0.02 | 0.00 | 0.00 | 0.00 |

| | | | | | | | | | | | | | | | |
|---|---|---|---|---|---|---|---|---|---|---|---|---|---|---|---|
| | 100 | 0.23 | 0.16 | 104 | 4 | 0.22 | 0.24 | 0.26 | 0.17 | 0.05 | 0.04 | 0.01 | 0.00 | 0.00 | 0.01 |
| | 200 | 0.31 | 0.22 | 91 | 4 | 0.15 | 0.14 | 0.24 | 0.18 | 0.13 | 0.05 | 0.04 | 0.01 | 0.01 | 0.03 |
| | 400 | 0.40 | 0.19 | 76 | 4 | 0.04 | 0.07 | 0.16 | 0.34 | 0.20 | 0.05 | 0.07 | 0.04 | 0.01 | 0.03 |

| | | | | | | | | | | | | | | | |
|---|---|---|---|---|---|---|---|---|---|---|---|---|---|---|---|
| RP437 *fliM*$_{\Delta N}$-*YPet* Δ*cheA* transformed with *fliM*$_{N}$-*cheY*(D13K)-pTRC99a-FF4 (EW739) | 0 | 0.35 | 0.23 | 123 | 1 | 0.12 | 0.09 | 0.24 | 0.26 | 0.13 | 0.05 | 0.02 | 0.01 | 0.03 | 0.05 |
| | 200 | 1.00 | 0.01 | 48 | 1 | 0.00 | 0.00 | 0.00 | 0.00 | 0.00 | 0.00 | 0.00 | 0.00 | 0.00 | 1.00 |
| | 400 | 0.87 | 0.32 | 16 | 1 | 0.13 | 0.00 | 0.00 | 0.00 | 0.00 | 0.00 | 0.00 | 0.00 | 0.00 | 0.88 |
| | 0 | 0.25 | 0.16 | 79 | 2 | 0.14 | 0.27 | 0.28 | 0.18 | 0.09 | 0.03 | 0.01 | 0.00 | 0.00 | 0.01 |
| | 0 | 0.21 | 0.15 | 60 | 2 | 0.27 | 0.25 | 0.22 | 0.17 | 0.08 | 0.00 | 0.00 | 0.00 | 0.02 | 0.00 |
| | 1 | 0.33 | 0.22 | 119 | 2 | 0.12 | 0.18 | 0.22 | 0.20 | 0.13 | 0.04 | 0.03 | 0.02 | 0.03 | 0.04 |
| | 10 | 0.92 | 0.20 | 57 | 2 | 0.00 | 0.04 | 0.00 | 0.02 | 0.04 | 0.00 | 0.02 | 0.00 | 0.04 | 0.86 |
| | 100 | 1.00 | 0.02 | 51 | 2 | 0.00 | 0.00 | 0.00 | 0.00 | 0.00 | 0.00 | 0.00 | 0.00 | 0.00 | 1.00 |
| | 0 | 0.30 | 0.24 | 85 | 3 | 0.21 | 0.16 | 0.26 | 0.12 | 0.11 | 0.06 | 0.00 | 0.00 | 0.02 | 0.06 |
| | 10 | 0.95 | 0.14 | 69 | 3 | 0.00 | 0.00 | 0.01 | 0.01 | 0.00 | 0.00 | 0.01 | 0.04 | 0.07 | 0.84 |
| | 50 | 0.99 | 0.01 | 71 | 3 | 0.00 | 0.00 | 0.00 | 0.00 | 0.00 | 0.00 | 0.00 | 0.00 | 0.00 | 1.00 |
| | 75 | 0.99 | 0.02 | 76 | 3 | 0.00 | 0.00 | 0.00 | 0.00 | 0.00 | 0.00 | 0.00 | 0.00 | 0.01 | 0.99 |
| | 100 | 1.00 | 0.01 | 60 | 3 | 0.00 | 0.00 | 0.00 | 0.00 | 0.00 | 0.00 | 0.00 | 0.00 | 0.00 | 1.00 |
| | 0 | 0.33 | 0.27 | 21 | 4 | 0.14 | 0.14 | 0.33 | 0.19 | 0.00 | 0.05 | 0.00 | 0.05 | 0.00 | 0.10 |
| | 1 | 0.33 | 0.33 | 40 | 4 | 0.25 | 0.18 | 0.33 | 0.05 | 0.00 | 0.00 | 0.00 | 0.03 | 0.00 | 0.18 |
| | 2 | 0.58 | 0.35 | 30 | 4 | 0.03 | 0.10 | 0.20 | 0.13 | 0.07 | 0.03 | 0.00 | 0.00 | 0.07 | 0.37 |
| | 5 | 0.61 | 0.30 | 45 | 4 | 0.04 | 0.00 | 0.09 | 0.20 | 0.13 | 0.04 | 0.02 | 0.09 | 0.13 | 0.24 |

| | | | | | | | | | | | | | | | |
|---|---|---|---|---|---|---|---|---|---|---|---|---|---|---|---|
| RP437 Δ*fliM* Δ*fliN* Δ*cheZ* transformed with *fliM*$_{N}$-*cheY*-pTRC99a-FF4 and *fliM*$_{\Delta N}$ *fliN*(A93D)-pKG116 (EW714) + 10 mM Acetate | 500 | 0.15 | 0.17 | 40 | 1 | 0.53 | 0.10 | 0.18 | 0.10 | 0.08 | 0.00 | 0.03 | 0.00 | 0.00 | 0.00 |
| | 1000 | 0.37 | 0.25 | 36 | 1 | 0.11 | 0.17 | 0.14 | 0.25 | 0.11 | 0.06 | 0.00 | 0.06 | 0.08 | 0.03 |
| | 2000 | 0.56 | 0.39 | 12 | 1 | 0.08 | 0.17 | 0.17 | 0.00 | 0.08 | 0.08 | 0.00 | 0.00 | 0.08 | 0.33 |
| | 4000 | 0.48 | 0.34 | 27 | 1 | 0.15 | 0.07 | 0.15 | 0.11 | 0.11 | 0.04 | 0.04 | 0.07 | 0.07 | 0.19 |
| | 8000 | 0.53 | 0.31 | 38 | 1 | 0.08 | 0.05 | 0.13 | 0.11 | 0.13 | 0.05 | 0.16 | 0.03 | 0.08 | 0.18 |
| | 125 | 0.42 | 0.25 | 10 | 2 | 0.00 | 0.20 | 0.10 | 0.30 | 0.20 | 0.00 | 0.00 | 0.00 | 0.20 | 0.00 |
| | 250 | 0.20 | 0.23 | 5 | 2 | 0.40 | 0.20 | 0.20 | 0.00 | 0.00 | 0.20 | 0.00 | 0.00 | 0.00 | 0.00 |
| | 500 | 0.37 | 0.38 | 23 | 2 | 0.39 | 0.04 | 0.09 | 0.09 | 0.04 | 0.04 | 0.00 | 0.13 | 0.00 | 0.17 |
| | 1000 | 0.18 | 0.32 | 17 | 2 | 0.65 | 0.12 | 0.06 | 0.00 | 0.00 | 0.00 | 0.06 | 0.00 | 0.06 | 0.06 |
| | 2000 | 0.33 | 0.32 | 21 | 2 | 0.24 | 0.19 | 0.19 | 0.10 | 0.00 | 0.05 | 0.05 | 0.10 | 0.00 | 0.10 |
| | 4000 | 0.59 | 0.23 | 17 | 2 | 0.00 | 0.06 | 0.06 | 0.00 | 0.18 | 0.18 | 0.29 | 0.06 | 0.06 | 0.12 |
| | 8000 | 0.67 | 0.40 | 17 | 2 | 0.24 | 0.00 | 0.00 | 0.00 | 0.06 | 0.00 | 0.00 | 0.12 | 0.06 | 0.53 |
| | 2000 | 0.27 | 0.22 | 57 | 3 | 0.25 | 0.14 | 0.21 | 0.19 | 0.09 | 0.05 | 0.00 | 0.04 | 0.02 | 0.02 |
| | 4000 | 0.28 | 0.23 | 69 | 3 | 0.30 | 0.04 | 0.23 | 0.14 | 0.14 | 0.06 | 0.01 | 0.01 | 0.01 | 0.03 |
| | 8000 | 0.22 | 0.24 | 103 | 3 | 0.41 | 0.15 | 0.18 | 0.11 | 0.06 | 0.01 | 0.03 | 0.00 | 0.03 | 0.03 |
| | 500 | 0.28 | 0.26 | 106 | 4 | 0.34 | 0.11 | 0.11 | 0.16 | 0.10 | 0.06 | 0.03 | 0.03 | 0.04 | 0.02 |
| | 2000 | 0.12 | 0.20 | 63 | 4 | 0.68 | 0.08 | 0.11 | 0.05 | 0.03 | 0.02 | 0.00 | 0.00 | 0.03 | 0.00 |
| | 8000 | 0.45 | 0.31 | 55 | 4 | 0.20 | 0.04 | 0.11 | 0.15 | 0.16 | 0.04 | 0.07 | 0.04 | 0.09 | 0.11 |

| | | | | | | | | | | | | | | | |
|---|---|---|---|---|---|---|---|---|---|---|---|---|---|---|---|
| | 50 | 0.01 | 0.02 | 16 | 5 | 1.00 | 0.00 | 0.00 | 0.00 | 0.00 | 0.00 | 0.00 | 0.00 | 0.00 | 0.00 |
| | 100 | 0.20 | 0.18 | 6 | 5 | 0.33 | 0.33 | 0.00 | 0.17 | 0.17 | 0.00 | 0.00 | 0.00 | 0.00 | 0.00 |
| | 500 | 0.08 | 0.10 | 21 | 5 | 0.71 | 0.10 | 0.19 | 0.00 | 0.00 | 0.00 | 0.00 | 0.00 | 0.00 | 0.00 |
| | 2000 | 0.26 | 0.19 | 9 | 5 | 0.22 | 0.22 | 0.22 | 0.11 | 0.11 | 0.11 | 0.00 | 0.00 | 0.00 | 0.00 |
| | 8000 | 0.09 | 0.17 | 37 | 5 | 0.81 | 0.03 | 0.08 | 0.05 | 0.00 | 0.00 | 0.00 | 0.00 | 0.03 | 0.00 |
| | 8000 | 0.34 | 0.22 | 80 | 6 | 0.19 | 0.09 | 0.14 | 0.25 | 0.13 | 0.06 | 0.09 | 0.03 | 0.03 | 0.01 |
| | 8000 | 0.39 | 0.23 | 80 | 7 | 0.11 | 0.09 | 0.13 | 0.24 | 0.13 | 0.16 | 0.06 | 0.01 | 0.04 | 0.04 |
| | 8000 | 0.23 | 0.22 | 51 | 8 | 0.39 | 0.06 | 0.16 | 0.16 | 0.16 | 0.04 | 0.02 | 0.00 | 0.00 | 0.02 |
| | 8000 | 0.27 | 0.18 | 65 | 9 | 0.26 | 0.05 | 0.22 | 0.23 | 0.15 | 0.05 | 0.02 | 0.03 | 0.00 | 0.00 |
| | 8000 | 0.56 | 0.28 | 15 | 10 | 0.00 | 0.07 | 0.07 | 0.13 | 0.20 | 0.20 | 0.07 | 0.00 | 0.07 | 0.20 |
| | 8000 | 0.26 | 0.16 | 32 | 11 | 0.19 | 0.09 | 0.31 | 0.22 | 0.13 | 0.06 | 0.00 | 0.00 | 0.00 | 0.00 |
| | 8000 | 0.26 | 0.18 | 96 | 12 | 0.22 | 0.19 | 0.18 | 0.18 | 0.16 | 0.04 | 0.01 | 0.03 | 0.00 | 0.00 |
| | 8000 | 0.29 | 0.24 | 54 | 13 | 0.28 | 0.07 | 0.19 | 0.15 | 0.11 | 0.11 | 0.02 | 0.06 | 0.02 | 0.00 |
| | 8000 | 0.23 | 0.29 | 100 | 14 | 0.58 | 0.01 | 0.01 | 0.07 | 0.13 | 0.09 | 0.04 | 0.01 | 0.03 | 0.03 |

| | | | | | | | | | | | | | | | |
|---|---|---|---|---|---|---|---|---|---|---|---|---|---|---|---|
| RP437 Δ*fliM* Δ*fliN* Δ*cheZ* transformed with *fliM*$_N$-*cheY*-pTRC99a-FF4 and *fliM*$_{\Delta N}$ *fliN*(A93D)-pKG116 (EW714) | 500 | 0.00 | 0.00 | 40 | 1 | 1.00 | 0.00 | 0.00 | 0.00 | 0.00 | 0.00 | 0.00 | 0.00 | 0.00 | 0.00 |
| | 1000 | 0.00 | 0.01 | 24 | 1 | 1.00 | 0.00 | 0.00 | 0.00 | 0.00 | 0.00 | 0.00 | 0.00 | 0.00 | 0.00 |
| | 2000 | 0.00 | 0.00 | 18 | 1 | 1.00 | 0.00 | 0.00 | 0.00 | 0.00 | 0.00 | 0.00 | 0.00 | 0.00 | 0.00 |
| | 4000 | 0.00 | 0.00 | 38 | 1 | 1.00 | 0.00 | 0.00 | 0.00 | 0.00 | 0.00 | 0.00 | 0.00 | 0.00 | 0.00 |
| | 8000 | 0.00 | 0.01 | 50 | 1 | 1.00 | 0.00 | 0.00 | 0.00 | 0.00 | 0.00 | 0.00 | 0.00 | 0.00 | 0.00 |
| | 125 | 0.01 | 0.02 | 28 | 2 | 1.00 | 0.00 | 0.00 | 0.00 | 0.00 | 0.00 | 0.00 | 0.00 | 0.00 | 0.00 |
| | 250 | 0.02 | 0.05 | 13 | 2 | 0.92 | 0.08 | 0.00 | 0.00 | 0.00 | 0.00 | 0.00 | 0.00 | 0.00 | 0.00 |
| | 500 | 0.00 | 0.00 | 21 | 2 | 1.00 | 0.00 | 0.00 | 0.00 | 0.00 | 0.00 | 0.00 | 0.00 | 0.00 | 0.00 |
| | 1000 | 0.00 | 0.00 | 3 | 2 | 1.00 | 0.00 | 0.00 | 0.00 | 0.00 | 0.00 | 0.00 | 0.00 | 0.00 | 0.00 |
| | 2000 | 0.00 | 0.00 | 25 | 2 | 1.00 | 0.00 | 0.00 | 0.00 | 0.00 | 0.00 | 0.00 | 0.00 | 0.00 | 0.00 |
| | 4000 | 0.00 | 0.00 | 9 | 2 | 1.00 | 0.00 | 0.00 | 0.00 | 0.00 | 0.00 | 0.00 | 0.00 | 0.00 | 0.00 |
| | 8000 | 0.00 | 0.01 | 25 | 2 | 1.00 | 0.00 | 0.00 | 0.00 | 0.00 | 0.00 | 0.00 | 0.00 | 0.00 | 0.00 |
| | 8000 | 0.00 | 0.00 | 108 | 3 | 1.00 | 0.00 | 0.00 | 0.00 | 0.00 | 0.00 | 0.00 | 0.00 | 0.00 | 0.00 |
| | 8000 | 0.00 | 0.00 | 92 | 4 | 1.00 | 0.00 | 0.00 | 0.00 | 0.00 | 0.00 | 0.00 | 0.00 | 0.00 | 0.00 |
| | 8000 | 0.00 | 0.00 | 42 | 5 | 1.00 | 0.00 | 0.00 | 0.00 | 0.00 | 0.00 | 0.00 | 0.00 | 0.00 | 0.00 |
| | 8000 | 0.00 | 0.00 | 47 | 6 | 1.00 | 0.00 | 0.00 | 0.00 | 0.00 | 0.00 | 0.00 | 0.00 | 0.00 | 0.00 |
| | 8000 | 0.00 | 0.00 | 44 | 7 | 1.00 | 0.00 | 0.00 | 0.00 | 0.00 | 0.00 | 0.00 | 0.00 | 0.00 | 0.00 |
| | 8000 | 0.00 | 0.02 | 80 | 8 | 0.99 | 0.01 | 0.00 | 0.00 | 0.00 | 0.00 | 0.00 | 0.00 | 0.00 | 0.00 |
| | 8000 | 0.00 | 0.00 | 45 | 9 | 1.00 | 0.00 | 0.00 | 0.00 | 0.00 | 0.00 | 0.00 | 0.00 | 0.00 | 0.00 |

| | | | | | | | | | | | | | | | |
|---|---|---|---|---|---|---|---|---|---|---|---|---|---|---|---|
| | 8000 | 0.00 | 0.01 | 152 | 10 | 1.00 | 0.00 | 0.00 | 0.00 | 0.00 | 0.00 | 0.00 | 0.00 | 0.00 | 0.00 |
| | 8000 | 0.00 | 0.00 | 86 | 11 | 1.00 | 0.00 | 0.00 | 0.00 | 0.00 | 0.00 | 0.00 | 0.00 | 0.00 | 0.00 |
| | 8000 | 0.01 | 0.10 | 85 | 11 | 0.98 | 0.01 | 0.00 | 0.00 | 0.00 | 0.00 | 0.00 | 0.00 | 0.00 | 0.01 |

| | | | | | | | | | | | | | | | |
|---|---|---|---|---|---|---|---|---|---|---|---|---|---|---|---|
| RP437 Δ*fliM* Δ*fliN* Δ*cheZ* transformed with *fliM*$_N$-*cheY*-pTRC99a-FF4 and *fliM*$_{\Delta N}$ *fliN*(A93D)-pKG116 (EW714) + 10 mM Benzoate | 8000 | 0.01 | 0.07 | 70 | 1 | 0.96 | 0.01 | 0.01 | 0.00 | 0.00 | 0.01 | 0.00 | 0.00 | 0.00 | 0.00 |
| | 8000 | 0.00 | 0.00 | 6 | 2 | 1.00 | 0.00 | 0.00 | 0.00 | 0.00 | 0.00 | 0.00 | 0.00 | 0.00 | 0.00 |
| | 8000 | 0.01 | 0.05 | 180 | 3 | 0.98 | 0.01 | 0.00 | 0.01 | 0.00 | 0.01 | 0.00 | 0.00 | 0.00 | 0.00 |
| | 8000 | 0.03 | 0.08 | 72 | 4 | 0.88 | 0.03 | 0.07 | 0.03 | 0.00 | 0.00 | 0.00 | 0.00 | 0.00 | 0.00 |
| | 8000 | 0.00 | 0.02 | 131 | 5 | 0.99 | 0.00 | 0.01 | 0.00 | 0.00 | 0.00 | 0.00 | 0.00 | 0.00 | 0.00 |
| | 8000 | 0.00 | 0.00 | 1 | 6 | 1.00 | 0.00 | 0.00 | 0.00 | 0.00 | 0.00 | 0.00 | 0.00 | 0.00 | 0.00 |
| | 8000 | 0.00 | 0.01 | 53 | 7 | 1.00 | 0.00 | 0.00 | 0.00 | 0.00 | 0.00 | 0.00 | 0.00 | 0.00 | 0.00 |
| | 8000 | 0.01 | 0.03 | 200 | 8 | 0.98 | 0.02 | 0.01 | 0.01 | 0.00 | 0.00 | 0.00 | 0.00 | 0.00 | 0.00 |
| | 8000 | 0.00 | 0.01 | 117 | 9 | 1.00 | 0.00 | 0.00 | 0.00 | 0.00 | 0.00 | 0.00 | 0.00 | 0.00 | 0.00 |
| | 8000 | 0.02 | 0.11 | 125 | 10 | 0.94 | 0.03 | 0.01 | 0.00 | 0.00 | 0.00 | 0.00 | 0.00 | 0.02 | 0.00 |

| | | | | | | | | | | | | | | | |
|---|---|---|---|---|---|---|---|---|---|---|---|---|---|---|---|
| RP437 Δ*fliM* Δ*fliN* Δ*cheA* transformed with *fliM*$_N$-*cheY*-pTRC99a-FF4 and *fliM*$_{\Delta N}$ *fliN*(A93D)-pKG116 (EW713) | 8000 | 0.00 | 0.00 | 57 | 1 | 1.00 | 0.00 | 0.00 | 0.00 | 0.00 | 0.00 | 0.00 | 0.00 | 0.00 | 0.00 |

| | | | | | | | | | | | | | | | |
|---|---|---|---|---|---|---|---|---|---|---|---|---|---|---|---|
| RP437 Δ*fliM* Δ*fliN* Δ*cheA* transformed with *fliM*$_N$-*cheY*-pTRC99a-FF4 and *fliM*$_{\Delta N}$ *fliN*(A93D)-pKG116 (EW713) + 10 mM Acetate | 500 | 0.09 | 0.19 | 45 | 1 | 0.78 | 0.04 | 0.02 | 0.04 | 0.02 | 0.07 | 0.02 | 0.00 | 0.00 | 0.00 |
| | 4000 | 0.24 | 0.35 | 43 | 1 | 0.60 | 0.02 | 0.02 | 0.12 | 0.05 | 0.00 | 0.02 | 0.02 | 0.02 | 0.12 |
| | 500 | 0.06 | 0.13 | 44 | 2 | 0.82 | 0.02 | 0.05 | 0.11 | 0.00 | 0.00 | 0.00 | 0.00 | 0.00 | 0.00 |
| | 2000 | 0.05 | 0.13 | 57 | 2 | 0.88 | 0.02 | 0.04 | 0.02 | 0.04 | 0.02 | 0.00 | 0.00 | 0.00 | 0.00 |
| | 8000 | 0.02 | 0.08 | 30 | 2 | 0.93 | 0.00 | 0.03 | 0.03 | 0.00 | 0.00 | 0.00 | 0.00 | 0.00 | 0.00 |
| | 50 | 0.01 | 0.03 | 46 | 3 | 0.98 | 0.00 | 0.02 | 0.00 | 0.00 | 0.00 | 0.00 | 0.00 | 0.00 | 0.00 |
| | 100 | 0.04 | 0.07 | 32 | 3 | 0.88 | 0.09 | 0.00 | 0.03 | 0.00 | 0.00 | 0.00 | 0.00 | 0.00 | 0.00 |
| | 500 | 0.01 | 0.02 | 14 | 3 | 1.00 | 0.00 | 0.00 | 0.00 | 0.00 | 0.00 | 0.00 | 0.00 | 0.00 | 0.00 |
| | 2000 | 0.01 | 0.02 | 8 | 3 | 1.00 | 0.00 | 0.00 | 0.00 | 0.00 | 0.00 | 0.00 | 0.00 | 0.00 | 0.00 |
| | 8000 | 0.02 | 0.06 | 18 | 3 | 0.89 | 0.06 | 0.06 | 0.00 | 0.00 | 0.00 | 0.00 | 0.00 | 0.00 | 0.00 |
| | 50 | 0.03 | 0.10 | 14 | 4 | 0.93 | 0.00 | 0.00 | 0.07 | 0.00 | 0.00 | 0.00 | 0.00 | 0.00 | 0.00 |
| | 100 | 0.03 | 0.07 | 7 | 4 | 0.86 | 0.14 | 0.00 | 0.00 | 0.00 | 0.00 | 0.00 | 0.00 | 0.00 | 0.00 |
| | 500 | 0.00 | 0.00 | 4 | 4 | 1.00 | 0.00 | 0.00 | 0.00 | 0.00 | 0.00 | 0.00 | 0.00 | 0.00 | 0.00 |
| | 2000 | 0.12 | 0.18 | 2 | 4 | 0.50 | 0.00 | 0.50 | 0.00 | 0.00 | 0.00 | 0.00 | 0.00 | 0.00 | 0.00 |
| | 100 | 0.03 | 0.11 | 40 | 5 | 0.95 | 0.00 | 0.03 | 0.00 | 0.00 | 0.00 | 0.03 | 0.00 | 0.00 | 0.00 |
| | 2000 | 0.20 | 0.21 | 55 | 6 | 0.42 | 0.16 | 0.11 | 0.15 | 0.09 | 0.00 | 0.05 | 0.02 | 0.00 | 0.00 |
| | 4000 | 0.23 | 0.29 | 78 | 6 | 0.49 | 0.14 | 0.06 | 0.06 | 0.08 | 0.04 | 0.05 | 0.00 | 0.03 | 0.05 |
| | 8000 | 0.32 | 0.29 | 99 | 6 | 0.30 | 0.10 | 0.11 | 0.13 | 0.13 | 0.06 | 0.01 | 0.04 | 0.04 | 0.07 |

| | | | | | | | | | | | | | | | |
|---|---|---|---|---|---|---|---|---|---|---|---|---|---|---|---|
| RP437 Δ*fliM* Δ*fliN* Δ*cheZ* transformed with *fliM*$_{N}$-*cheY*-pTRC99a-FF4 and *fliM*$_{\Delta N}$(K94L)*fliN*-pKG116 (EW732) | 8000 | 0.00 | 0.00 | 27 | 1 | 1.00 | 0.00 | 0.00 | 0.00 | 0.00 | 0.00 | 0.00 | 0.00 | 0.00 | 0.00 |
| | 8000 | 0.00 | 0.00 | 81 | 2 | 1.00 | 0.00 | 0.00 | 0.00 | 0.00 | 0.00 | 0.00 | 0.00 | 0.00 | 0.00 |
| | 8000 | 0.00 | 0.00 | 296 | 3 | 1.00 | 0.00 | 0.00 | 0.00 | 0.00 | 0.00 | 0.00 | 0.00 | 0.00 | 0.00 |
| | 8000 | 0.00 | 0.00 | 57 | 4 | 1.00 | 0.00 | 0.00 | 0.00 | 0.00 | 0.00 | 0.00 | 0.00 | 0.00 | 0.00 |
| | 8000 | 0.00 | 0.00 | 56 | 5 | 1.00 | 0.00 | 0.00 | 0.00 | 0.00 | 0.00 | 0.00 | 0.00 | 0.00 | 0.00 |
| | 8000 | 0.00 | 0.01 | 36 | 6 | 1.00 | 0.00 | 0.00 | 0.00 | 0.00 | 0.00 | 0.00 | 0.00 | 0.00 | 0.00 |
| | 8000 | 0.00 | 0.00 | 144 | 7 | 1.00 | 0.00 | 0.00 | 0.00 | 0.00 | 0.00 | 0.00 | 0.00 | 0.00 | 0.00 |
| | 8000 | 0.00 | 0.00 | 37 | 8 | 1.00 | 0.00 | 0.00 | 0.00 | 0.00 | 0.00 | 0.00 | 0.00 | 0.00 | 0.00 |

| | | | | | | | | | | | | | | | |
|---|---|---|---|---|---|---|---|---|---|---|---|---|---|---|---|
| RP437 Δ*fliM* Δ*fliN* Δ*cheZ* transformed with *fliM*$_{N}$-*cheY*-pTRC99a-FF4 and *fliM*$_{\Delta N}$(K94L)*fliN*-pKG116 (EW732) + 10 mM Acetate | 8000 | 0.00 | 0.00 | 32 | 1 | 1.00 | 0.00 | 0.00 | 0.00 | 0.00 | 0.00 | 0.00 | 0.00 | 0.00 | 0.00 |
| | 8000 | 0.03 | 0.07 | 75 | 2 | 0.88 | 0.05 | 0.05 | 0.00 | 0.01 | 0.00 | 0.00 | 0.00 | 0.00 | 0.00 |
| | 8000 | 0.06 | 0.08 | 315 | 3 | 0.80 | 0.11 | 0.07 | 0.02 | 0.00 | 0.00 | 0.00 | 0.00 | 0.00 | 0.00 |
| | 8000 | 0.04 | 0.06 | 71 | 4 | 0.86 | 0.07 | 0.07 | 0.00 | 0.00 | 0.00 | 0.00 | 0.00 | 0.00 | 0.00 |
| | 8000 | 0.04 | 0.05 | 67 | 5 | 0.87 | 0.13 | 0.00 | 0.00 | 0.00 | 0.00 | 0.00 | 0.00 | 0.00 | 0.00 |
| | 8000 | 0.02 | 0.03 | 45 | 6 | 0.98 | 0.02 | 0.00 | 0.00 | 0.00 | 0.00 | 0.00 | 0.00 | 0.00 | 0.00 |
| | 8000 | 0.08 | 0.08 | 192 | 7 | 0.70 | 0.19 | 0.09 | 0.02 | 0.01 | 0.00 | 0.00 | 0.00 | 0.00 | 0.00 |
| | 8000 | 0.04 | 0.05 | 65 | 8 | 0.83 | 0.17 | 0.00 | 0.00 | 0.00 | 0.00 | 0.00 | 0.00 | 0.00 | 0.00 |
| | 8000 | 0.03 | 0.03 | 27 | 9 | 0.96 | 0.04 | 0.00 | 0.00 | 0.00 | 0.00 | 0.00 | 0.00 | 0.00 | 0.00 |

| | | | | | | | | | | | | | | | |
|---|---|---|---|---|---|---|---|---|---|---|---|---|---|---|---|
| RP437 Δ*fliM* Δ*fliN* Δ*cheZ* transformed with *fliM*$_N$-*cheY*-pTRC99a-FF4 and *fliM*$_{\Delta N}$ (K94L)*fliN*-pKG116 (EW732) + 10 mM Benzoate | 8000 | 0.00 | 0.00 | 45 | 1 | 1.00 | 0.00 | 0.00 | 0.00 | 0.00 | 0.00 | 0.00 | 0.00 | 0.00 | 0.00 |
| | 8000 | 0.00 | 0.00 | 291 | 2 | 1.00 | 0.00 | 0.00 | 0.00 | 0.00 | 0.00 | 0.00 | 0.00 | 0.00 | 0.00 |
| | 8000 | 0.00 | 0.00 | 117 | 3 | 1.00 | 0.00 | 0.00 | 0.00 | 0.00 | 0.00 | 0.00 | 0.00 | 0.00 | 0.00 |
| | 8000 | 0.00 | 0.00 | 59 | 4 | 1.00 | 0.00 | 0.00 | 0.00 | 0.00 | 0.00 | 0.00 | 0.00 | 0.00 | 0.00 |
| | 8000 | 0.00 | 0.00 | 135 | 5 | 1.00 | 0.00 | 0.00 | 0.00 | 0.00 | 0.00 | 0.00 | 0.00 | 0.00 | 0.00 |
| | 8000 | 0.00 | 0.00 | 45 | 6 | 1.00 | 0.00 | 0.00 | 0.00 | 0.00 | 0.00 | 0.00 | 0.00 | 0.00 | 0.00 |

| | | | | | | | | | | | | | | | |
|---|---|---|---|---|---|---|---|---|---|---|---|---|---|---|---|
| RP437 Δ*fliM* Δ*fliN* Δ*cheZ* transformed with *fliM*$_N$-*cheY*-pTRC99a-FF4 and *fliM*$_{\Delta N}$ (K94S)*fliN*-pKG116 (EW734) | 8000 | 0.00 | 0.00 | 103 | 1 | 1.00 | 0.00 | 0.00 | 0.00 | 0.00 | 0.00 | 0.00 | 0.00 | 0.00 | 0.00 |
| | 8000 | 0.00 | 0.00 | 38 | 2 | 1.00 | 0.00 | 0.00 | 0.00 | 0.00 | 0.00 | 0.00 | 0.00 | 0.00 | 0.00 |
| | 8000 | 0.00 | 0.00 | 83 | 3 | 1.00 | 0.00 | 0.00 | 0.00 | 0.00 | 0.00 | 0.00 | 0.00 | 0.00 | 0.00 |
| | 8000 | 0.00 | 0.00 | 158 | 4 | 1.00 | 0.00 | 0.00 | 0.00 | 0.00 | 0.00 | 0.00 | 0.00 | 0.00 | 0.00 |
| | 8000 | 0.00 | 0.00 | 61 | 5 | 1.00 | 0.00 | 0.00 | 0.00 | 0.00 | 0.00 | 0.00 | 0.00 | 0.00 | 0.00 |
| | 8000 | 0.00 | 0.00 | 51 | 6 | 1.00 | 0.00 | 0.00 | 0.00 | 0.00 | 0.00 | 0.00 | 0.00 | 0.00 | 0.00 |
| | 8000 | 0.00 | 0.00 | 50 | 7 | 1.00 | 0.00 | 0.00 | 0.00 | 0.00 | 0.00 | 0.00 | 0.00 | 0.00 | 0.00 |
| | 8000 | 0.00 | 0.00 | 135 | 8 | 1.00 | 0.00 | 0.00 | 0.00 | 0.00 | 0.00 | 0.00 | 0.00 | 0.00 | 0.00 |
| | 8000 | 0.00 | 0.00 | 47 | 9 | 1.00 | 0.00 | 0.00 | 0.00 | 0.00 | 0.00 | 0.00 | 0.00 | 0.00 | 0.00 |

| | | | | | | | | | | | | | | | |
|---|---|---|---|---|---|---|---|---|---|---|---|---|---|---|---|
| | 8000 | 0.00 | 0.00 | 35 | 10 | 1.00 | 0.00 | 0.00 | 0.00 | 0.00 | 0.00 | 0.00 | 0.00 | 0.00 | 0.00 |
| | 8000 | 0.00 | 0.00 | 41 | 11 | 1.00 | 0.00 | 0.00 | 0.00 | 0.00 | 0.00 | 0.00 | 0.00 | 0.00 | 0.00 |

| | | | | | | | | | | | | | | | |
|---|---|---|---|---|---|---|---|---|---|---|---|---|---|---|---|
| RP437 Δ*fliM* Δ*fliN* Δ*cheZ* transformed with *fliM*$_{N}$-*cheY*-pTRC99a-FF4 and *fliM*$_{\Delta N}$ (K94S)*fliN*-pKG116 (EW734) + 10 mM Acetate | 8000 | 0.03 | 0.03 | 8 | 1 | 1.00 | 0.00 | 0.00 | 0.00 | 0.00 | 0.00 | 0.00 | 0.00 | 0.00 | 0.00 |
| | 8000 | 0.01 | 0.02 | 46 | 2 | 0.98 | 0.02 | 0.00 | 0.00 | 0.00 | 0.00 | 0.00 | 0.00 | 0.00 | 0.00 |
| | 8000 | 0.08 | 0.11 | 90 | 3 | 0.68 | 0.20 | 0.04 | 0.06 | 0.02 | 0.00 | 0.00 | 0.00 | 0.00 | 0.00 |
| | 8000 | 0.14 | 0.12 | 218 | 4 | 0.44 | 0.30 | 0.13 | 0.09 | 0.04 | 0.00 | 0.00 | 0.00 | 0.00 | 0.00 |
| | 8000 | 0.12 | 0.13 | 72 | 5 | 0.63 | 0.17 | 0.06 | 0.11 | 0.04 | 0.00 | 0.00 | 0.00 | 0.00 | 0.00 |
| | 8000 | 0.07 | 0.10 | 51 | 6 | 0.73 | 0.16 | 0.08 | 0.02 | 0.02 | 0.00 | 0.00 | 0.00 | 0.00 | 0.00 |
| | 8000 | 0.07 | 0.09 | 38 | 7 | 0.82 | 0.08 | 0.05 | 0.05 | 0.00 | 0.00 | 0.00 | 0.00 | 0.00 | 0.00 |
| | 8000 | 0.16 | 0.14 | 162 | 8 | 0.43 | 0.22 | 0.14 | 0.16 | 0.04 | 0.01 | 0.00 | 0.00 | 0.00 | 0.00 |
| | 8000 | 0.06 | 0.08 | 46 | 9 | 0.76 | 0.15 | 0.07 | 0.02 | 0.00 | 0.00 | 0.00 | 0.00 | 0.00 | 0.00 |
| | 8000 | 0.05 | 0.08 | 31 | 10 | 0.84 | 0.06 | 0.10 | 0.00 | 0.00 | 0.00 | 0.00 | 0.00 | 0.00 | 0.00 |

| | | | | | | | | | | | | | | | |
|---|---|---|---|---|---|---|---|---|---|---|---|---|---|---|---|
| RP437 Δ*fliM* Δ*fliN* Δ*cheZ* transformed with *fliM*$_{N}$-*cheY*-pTRC99a-FF4 and *fliM*$_{\Delta N}$ (K94S)*fliN*-pKG116 (EW734) + | 8000 | 0.00 | 0.00 | 65 | 1 | 1.00 | 0.00 | 0.00 | 0.00 | 0.00 | 0.00 | 0.00 | 0.00 | 0.00 | 0.00 |
| | 8000 | 0.00 | 0.00 | 145 | 2 | 1.00 | 0.00 | 0.00 | 0.00 | 0.00 | 0.00 | 0.00 | 0.00 | 0.00 | 0.00 |
| | 8000 | 0.00 | 0.00 | 137 | 3 | 1.00 | 0.00 | 0.00 | 0.00 | 0.00 | 0.00 | 0.00 | 0.00 | 0.00 | 0.00 |
| | 8000 | 0.00 | 0.00 | 55 | 4 | 1.00 | 0.00 | 0.00 | 0.00 | 0.00 | 0.00 | 0.00 | 0.00 | 0.00 | 0.00 |
| | 8000 | 0.01 | 0.03 | 80 | 5 | 0.99 | 0.00 | 0.01 | 0.00 | 0.00 | 0.00 | 0.00 | 0.00 | 0.00 | 0.00 |
| | 8000 | 0.00 | 0.00 | 56 | 6 | 1.00 | 0.00 | 0.00 | 0.00 | 0.00 | 0.00 | 0.00 | 0.00 | 0.00 | 0.00 |

| |
|---|
| 10 mM Benzoate |

| | | | | | | | | | | | | | | | |
|---|---|---|---|---|---|---|---|---|---|---|---|---|---|---|---|
| RP437 Δ*fliM* Δ*fliN* Δ*cheZ* transformed with *fliM*$_{N}$-*cheY*-pTRC99a-FF4 and *fliM*$_{\Delta N}$ (E214W)*fliN*-pKG116 (EW718) | 0 | 0.01 | 0.04 | 41 | 1 | 0.98 | 0.00 | 0.02 | 0.00 | 0.00 | 0.00 | 0.00 | 0.00 | 0.00 | 0.00 |
| | 1 | 0.06 | 0.15 | 31 | 1 | 0.87 | 0.00 | 0.03 | 0.00 | 0.06 | 0.03 | 0.00 | 0.00 | 0.00 | 0.00 |
| | 5 | 0.21 | 0.29 | 48 | 1 | 0.56 | 0.10 | 0.04 | 0.13 | 0.02 | 0.04 | 0.00 | 0.02 | 0.00 | 0.08 |
| | 10 | 0.24 | 0.35 | 53 | 1 | 0.66 | 0.00 | 0.00 | 0.00 | 0.09 | 0.04 | 0.00 | 0.04 | 0.13 | 0.04 |
| | 20 | 0.33 | 0.39 | 21 | 1 | 0.57 | 0.00 | 0.00 | 0.00 | 0.05 | 0.00 | 0.05 | 0.14 | 0.14 | 0.05 |
| | 40 | 0.26 | 0.38 | 18 | 1 | 0.67 | 0.00 | 0.00 | 0.00 | 0.00 | 0.00 | 0.17 | 0.06 | 0.06 | 0.06 |
| | 5 | 0.15 | 0.24 | 109 | 2 | 0.67 | 0.06 | 0.04 | 0.07 | 0.07 | 0.03 | 0.01 | 0.02 | 0.01 | 0.03 |
| | 10 | 0.16 | 0.31 | 70 | 2 | 0.77 | 0.00 | 0.00 | 0.00 | 0.00 | 0.11 | 0.01 | 0.01 | 0.04 | 0.04 |
| | 20 | 0.36 | 0.43 | 82 | 2 | 0.57 | 0.01 | 0.00 | 0.00 | 0.01 | 0.00 | 0.04 | 0.06 | 0.10 | 0.21 |
| | 40 | 0.27 | 0.42 | 86 | 2 | 0.71 | 0.00 | 0.00 | 0.00 | 0.00 | 0.00 | 0.01 | 0.03 | 0.03 | 0.21 |
| | 800 | 0.20 | 0.38 | 92 | 2 | 0.77 | 0.00 | 0.00 | 0.01 | 0.00 | 0.00 | 0.01 | 0.01 | 0.08 | 0.12 |
| | 0 | 0.01 | 0.06 | 84 | 3 | 0.95 | 0.01 | 0.01 | 0.02 | 0.00 | 0.00 | 0.00 | 0.00 | 0.00 | 0.00 |
| | 1 | 0.03 | 0.14 | 50 | 3 | 0.96 | 0.00 | 0.00 | 0.00 | 0.02 | 0.00 | 0.00 | 0.00 | 0.00 | 0.02 |
| | 5 | 0.02 | 0.08 | 58 | 3 | 0.95 | 0.02 | 0.02 | 0.00 | 0.02 | 0.00 | 0.00 | 0.00 | 0.00 | 0.00 |
| | 10 | 0.11 | 0.18 | 40 | 3 | 0.73 | 0.05 | 0.08 | 0.03 | 0.05 | 0.08 | 0.00 | 0.00 | 0.00 | 0.00 |
| | 20 | 0.19 | 0.26 | 39 | 3 | 0.59 | 0.03 | 0.05 | 0.13 | 0.08 | 0.05 | 0.00 | 0.03 | 0.03 | 0.03 |

| | | | | | | | | | | | | | | | |
|---|---|---|---|---|---|---|---|---|---|---|---|---|---|---|---|
| (cont.) | 40 | 0.23 | 0.27 | 46 | 3 | 0.54 | 0.02 | 0.04 | 0.09 | 0.09 | 0.11 | 0.04 | 0.02 | 0.04 | 0.00 |
| | 60 | 0.24 | 0.34 | 23 | 3 | 0.65 | 0.00 | 0.00 | 0.00 | 0.00 | 0.13 | 0.04 | 0.09 | 0.09 | 0.00 |
| | 80 | 0.38 | 0.33 | 25 | 3 | 0.32 | 0.04 | 0.08 | 0.08 | 0.08 | 0.12 | 0.08 | 0.12 | 0.00 | 0.08 |
| | 5 | 0.02 | 0.02 | 2 | 4 | 1.00 | 0.00 | 0.00 | 0.00 | 0.00 | 0.00 | 0.00 | 0.00 | 0.00 | 0.00 |
| | 20 | 0.35 | 0.28 | 208 | 4 | 0.33 | 0.04 | 0.07 | 0.08 | 0.14 | 0.13 | 0.11 | 0.06 | 0.02 | 0.02 |
| | 40 | 0.52 | 0.30 | 92 | 4 | 0.20 | 0.01 | 0.01 | 0.02 | 0.11 | 0.18 | 0.14 | 0.13 | 0.15 | 0.04 |
| | 60 | 0.44 | 0.29 | 103 | 4 | 0.23 | 0.00 | 0.06 | 0.09 | 0.16 | 0.09 | 0.19 | 0.11 | 0.05 | 0.03 |
| | 80 | 0.42 | 0.32 | 87 | 4 | 0.32 | 0.00 | 0.01 | 0.03 | 0.13 | 0.17 | 0.11 | 0.10 | 0.07 | 0.05 |
| | 100 | 0.34 | 0.30 | 78 | 4 | 0.37 | 0.01 | 0.04 | 0.10 | 0.03 | 0.24 | 0.12 | 0.04 | 0.04 | 0.01 |
| | 800 | 0.16 | 0.26 | 56 | 4 | 0.73 | 0.00 | 0.00 | 0.00 | 0.09 | 0.09 | 0.07 | 0.02 | 0.00 | 0.00 |
| | 0 | 0.01 | 0.04 | 350 | 5 | 0.96 | 0.03 | 0.00 | 0.00 | 0.00 | 0.00 | 0.00 | 0.00 | 0.00 | 0.00 |
| | 50 | 0.41 | 0.26 | 341 | 5 | 0.13 | 0.11 | 0.12 | 0.16 | 0.16 | 0.10 | 0.07 | 0.05 | 0.04 | 0.05 |
| | 100 | 0.51 | 0.22 | 193 | 5 | 0.06 | 0.04 | 0.06 | 0.11 | 0.18 | 0.22 | 0.17 | 0.05 | 0.08 | 0.03 |
| | 200 | 0.52 | 0.23 | 260 | 5 | 0.04 | 0.03 | 0.11 | 0.12 | 0.21 | 0.17 | 0.12 | 0.07 | 0.08 | 0.07 |
| | 400 | 0.53 | 0.25 | 234 | 5 | 0.08 | 0.02 | 0.05 | 0.11 | 0.18 | 0.19 | 0.14 | 0.09 | 0.03 | 0.10 |
| | 800 | 0.56 | 0.26 | 225 | 5 | 0.09 | 0.01 | 0.05 | 0.07 | 0.13 | 0.17 | 0.19 | 0.11 | 0.07 | 0.11 |
| | 800 | 0.44 | 0.22 | 158 | 5 | 0.09 | 0.03 | 0.10 | 0.17 | 0.21 | 0.18 | 0.09 | 0.07 | 0.03 | 0.03 |

***Table S2. Strains used in this study***

| Strain | Description | Source |
|---|---|---|
| B275 | $P^-$, *lacY*1, $TL^-$, *metF*159(Am), *rpsL*136(strR) | [11] |
| JW1929 | $F^-$, Δ(*araD-araB*)567, Δ*lacZ*4787(::*rrnB*-3), $\lambda^-$, *ΔfliM*720::kan, rph-1, Δ(*rhaD-rhaB*)568, *hsdR*514 | [12] |
| JW1870 | $F^-$, Δ(*araD-araB*)567, Δ*lacZ*4787(::*rrnB*-3), $\lambda^-$, *ΔcheZ734*::kan, rph-1, Δ(*rhaD-rhaB*)568, *hsdR*514 | [12] |
| JW1877 | $F^-$, Δ(*araD-araB*)567, Δ*lacZ*4787(::*rrnB*-3), $\lambda^-$, Δ*cheA*741::kan, rph-1, Δ(*rhaD-rhaB*)568, *hsdR*514 | [12] |
| RP437 | $F^-$, thr-1, *araC*14, *leuB*6(Am), *fhuA*31, *lacY*1, *tsx*-78, $\lambda^-$, *eda*-50, *hisG*4(Oc), *rfbC*1, *rpsL*136(strR), *xylA*5, mtl-1, *metF*159(Am), *thiE*1 | [13] |
| UU1631 | RP437 bearing in frame Δ*cheY* deletion mutation | S. Parkinson |
| JPA945 | RP437 with genomic fliM-Ypet produced by allelic exchange | [14] |
| sPW416 | RP437Δ*fliM* Δ*fliN* transformed with *cheY*-pTRC99a-FF4 and *fliM*Δ(1-16)*fliN*-pKG116 | D. Blair |
| sPW417 | RP437 Δ*fliM* Δ*fliN* transformed with *fliM*$_{N-34}$-*cheY*-pTRC99a-FF4 and *fliM*Δ(1-16)*fliN*-pKG116 | D. Blair |
| sPW455 | RP437 Δ*fliM* Δ*fliN* transformed with *fliM*$_{N-34}$-*cheY*-pTRC99a-FF4 and *fliM*Δ(1-16)(N201W)*fliN*-pKG116 | D. Blair |
| sPW456 | RP437 Δ*fliM* Δ*fliN* transformed with *fliM*$_{N-34}$-*cheY*-pTRC99a-FF4 and *fliM*Δ(1-16)*fliN*(A93D)-pKG116 | D. Blair |
| sPW457 | RP437 Δ*fliM* Δ*fliN* transformed with *fliM*$_{N-34}$-*cheY*-pTRC99a-FF4 and *fliM*Δ(1-16)*fliN*(V113D)-pKG116 | D. Blair |

| | | |
|---|---|---|
| sPW462 | RP437 Δ*fliM* Δ*fliN* transformed with *fliM*$_{N\text{-}34}$-*cheY*-pTRC99a-FF4 and *fliM*Δ(1-16)(E214W)*fliN*-pKG116 | D. Blair |
| sPW585 | RP437 Δ*fliM* Δ*fliN* transformed with *fliM*$_{N\text{-}34}$-*cheY*-pTRC99a-FF4 and *fliM*Δ(1-16)(R94L)*fliN*-pKG116 | D. Blair |
| sPW586 | RP437 Δ*fliM* Δ*fliN* transformed with *fliM*$_{N\text{-}34}$-*cheY*-pTRC99a-FF4 and *fliM*Δ(1-16) (R94S)*fliN*-pKG116 | D. Blair |
| EW535 | B275 P1 transduced with Δ*fliM* from JW1929 | This study |
| EW496 | EW535 transformed with *fliM*-pCA24N | This study |
| EW497 | EW535 transformed with *fliM*Δ(1-30)-pCA24N | This study |
| EW542 | EW497 transformed with pRL22Δ*cheZ* | This study |
| EW566 | JPA945 with genomic *fliM*Δ(1-16)-*YPet* produced by allelic exchange as described by Philippe et al. (2004) | This study |
| EW568 | EW566 transformed with *cheY*-pCA24N | This study |
| EW575 | EW566 transformed with *cheY-mCherry*-pCA24N | This study |
| EW634 | EW568 P1 transduced with Δ*cheA* from JW1870 | This study |
| EW635 | EW568 P1 transduced with Δ*cheZ* from JW1877 | This study |
| EW636 | EW575 P1 transduced with Δ*cheA* from JW1870 | This study |
| EW637 | EW575 P1 transduced with Δ*cheZ* from JW1877 | This study |
| EW659 | EW566 transformed with *mCherry*-pCA24N | This study |
| EW668 | JPA945 P1 transduced with Δ*cheA* from JW1877 | This study |
| EW669 | JPA945 P1 transduced with Δ*cheZ* from JW1870 | This study |
| EW670 | RP437 transformed with *cheY*(I95)-pCA24N | This study |
| EW677 | JPA945 transformed with *cheY-mCherry*-pCA24N | This study |
| EW690 | EW566 P1 transduced with Δ*cheA* from JW1877 | This study |
| EW691 | EW566 P1 transduced with Δ*cheZ* from JW1870 | This study |
| EW693 | EW690 transformed with *cheY*-pTRC99a-FF4 | This study |

| | | |
|---|---|---|
| EW694 | EW691 transformed with *cheY*-pTRC99a-FF4 | This study |
| EW696 | EW690 transformed with *fliM*$_{N-34}$-*cheY*-pTRC99a-FF4 | This study |
| EW697 | EW691 transformed with *fliM*$_{N-34}$-*cheY*-pTRC99a-FF4 | This study |
| EW707 | sPW416 P1 transduced with Δ*cheA* from JW1877 | This study |
| EW708 | sPW416 P1 transduced with Δ*cheZ* from JW1870 | This study |
| EW709 | sPW417 P1 transduced with Δ*cheA* from JW1877 | This study |
| EW710 | sPW417 P1 transduced with Δ*cheZ* from JW1870 | This study |
| EW711 | sPW455 P1 transduced with Δ*cheA* from JW1877 | This study |
| EW712 | sPW455 P1 transduced with Δ*cheZ* from JW1870 | This study |
| EW713 | sPW456 P1 transduced with Δ*cheA* from JW1877 | This study |
| EW714 | sPW456 P1 transduced with Δ*cheZ* from JW1870 | This study |
| EW715 | sPW457 P1 transduced with Δ*cheA* from JW1877 | This study |
| EW716 | sPW457 P1 transduced with Δ*cheZ* from JW1870 | This study |
| EW717 | sPW462 P1 transduced with Δ*cheA* from JW1877 | This study |
| EW718 | sPW462 P1 transduced with Δ*cheZ* from JW1870 | This study |
| EW732 | sPW585 P1 transduced with Δ*cheZ* from JW1870 | This study |
| EW734 | sPW586 P1 transduced with Δ*cheZ* from JW1870 | This study |
| EW677 | JPA945 Δ*cheY* transformed with *cheY-mCherry*-pCA24N | This study |
| EW675 | EW669 transformed with *cheY-mCherry*-pCA24N | This study |
| EW737 | EW690 transformed with *cheY(D13K)-pTRC99a-FF4* | This study |
| EW739 | EW690 transformed with *fliM*$_{N-34}$-*cheY*(D13K)-pTRC99a-FF4 | This study |

***Table S3 Plasmids used in this study***

| Plasmid | Source |
| --- | --- |
| *6xHis-cheY*-pCA24N | [15] |
| *6xHis-cheY*(I95V)-pCA24N | This study |
| *6xHis-fliM*-pCA24N | [15] |
| *6xHis-fliM*Δ(1-30)-pCA24N | This study |
| *6xHis-cheY-mCherry*-pCA24N | This study |
| *6xHis-mCherry*-pCA24N | This study |
| *cheY*-pTRC99a-FF4 | D. Blair |
| *fliM*$_{N\text{-}34}$-*cheY*-pTRC99a-FF4 | D. Blair |
| *cheY*(D13K)-pTRC99a-FF4 | This study |
| *fliM*$_{N\text{-}34}$-*cheY*(D13K)-pTRC99a-FF4 | This study |
| *fliM*Δ(1-16) *fliN*-pKG116 | D. Blair |
| *fliM*Δ(1-16)*fliN*(A93D)-pKG116 | D. Blair |
| *fliM*Δ(1-16)(E214W)*fliN*-pKG116 | D. Blair |
| *fliM*Δ(1-16) (R94S)*fliN*-pKG116 | D. Blair |
| *fliM*Δ(1-16) (R94L)*fliN*-pKG116 | D. Blair |
| pRL22Δ*cheZ* | [16] |

#### Supplementary Figures

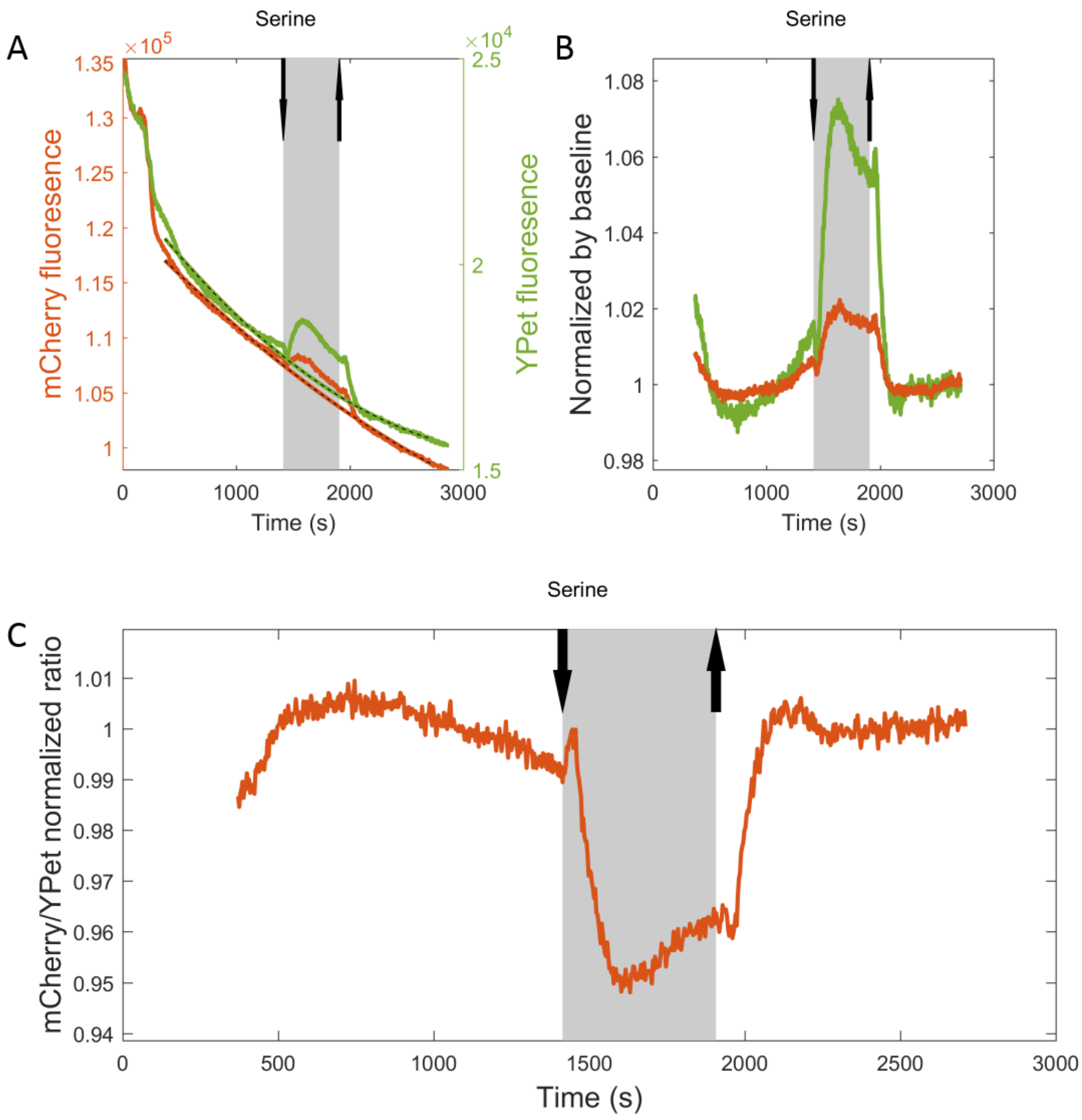

**Figure S1**. A detailed analysis of the functional interaction of CheY-mCherry with YPet-labeled motors (Figure 1D, blue). The experiment was done in a Δ*cheZ* genetic background using $FliM_{\Delta N}$-YPet and CheY-mCherry as a FRET pair (strain EW637). The expression of CheY-mCherry was induced with 400 µM IPTG for 4 h. Down and up arrows indicate addition and removal of serine (0.1 mM). Shaded regions show the time at which serine was present in the flow chamber. **A**, Raw readings of the fluorescence intensity of CheY-mCherry and $FliM_{\Delta N}$-YPet. Dotted curves show the estimated baselines of the measurements. **B**, The readings from A, normalized according to the estimated baseline. **C**, Calculated mCherry/YPet ratio from B.

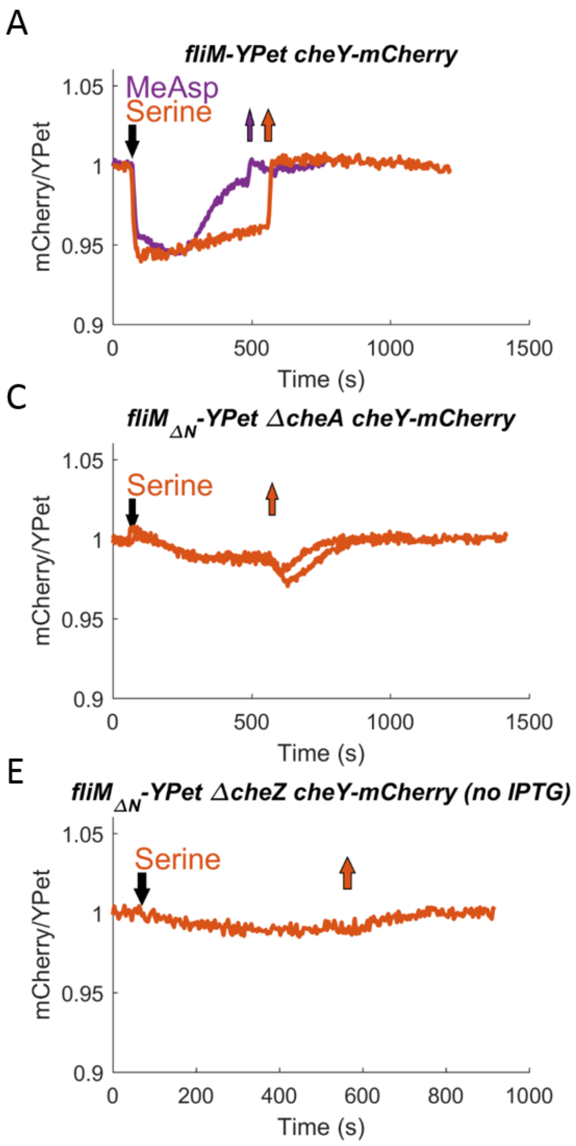

**Figure S2**. Changes in (CheY-mCherry)-(FliM$_{\Delta N}$-YPet) FRET reflect changes in the occupancy of CheY at the motor. **A**-**E**, Measured FRET responses in different backgrounds and CheY-mCherry expression levels. Each curve is an independent measurement of the response to attractant addition or removal (down or up arrows, respectively). Red and green curves are the responses to serine (at 0.1 and 1 mM, respectively) and purple curve is the response to 1 mM MeAsp. Strains used: EW677 (A), EW659 (B), EW636 (C), EW637 (D, E).

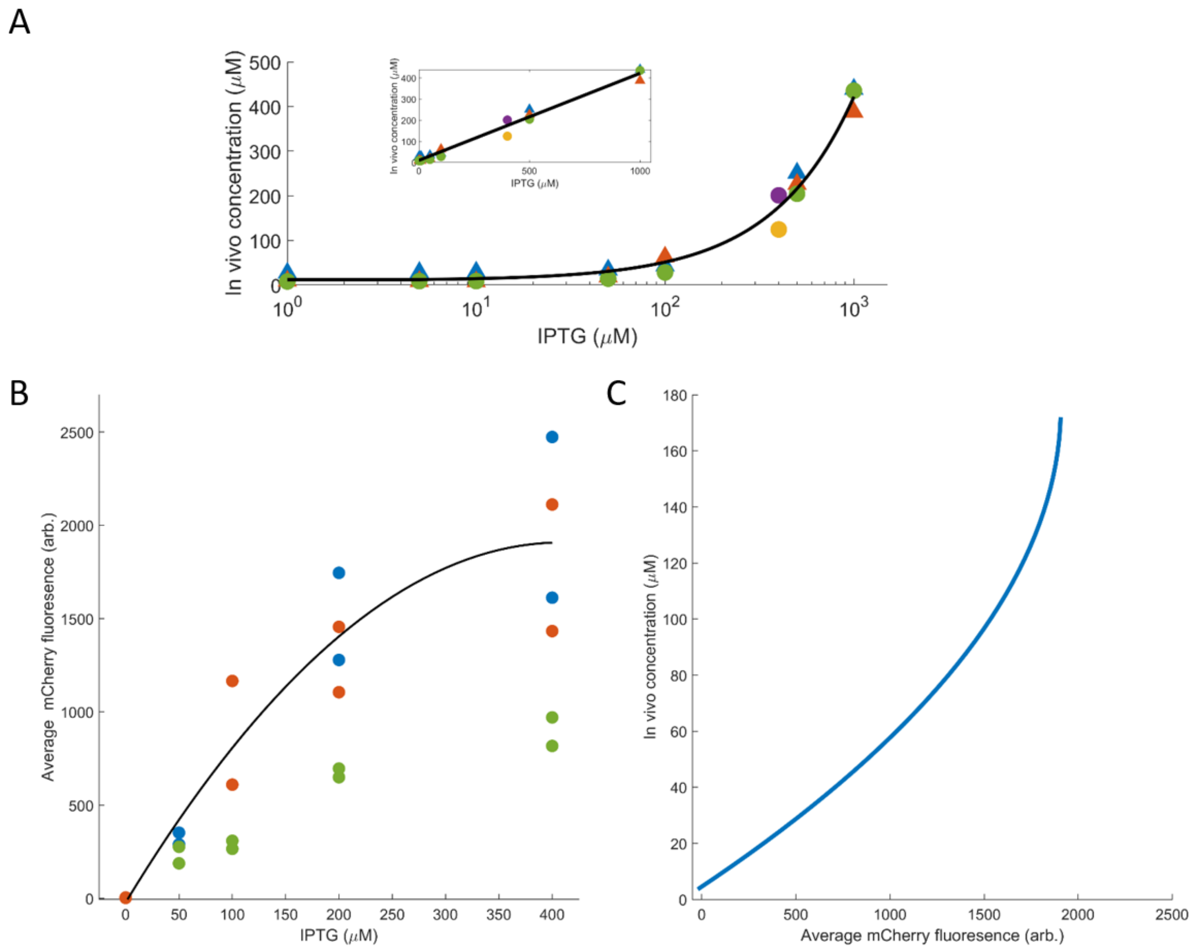

**Figure S3**. Quantification of the *in vivo* concentration of CheY. **A**, Calibration of CheY concentration at different IPTG levels. Cells expressing either CheY-mCherry or mCherry (strains EW575 and EW659, respectively) were incubated for 4-6 h with different concentrations of IPTG, and the *in vivo* concentration of the expressed protein was measured as described in Supplementary Methods above. Each color represents an independent repetition of the experiment. Note that the leaky expression of the *lac* promoter in this plasmid is equivalent to 10 μM protein concentration, which is roughly the concentration of chromosomally-expressed CheY in the cell. ***Insert***, as in main panel, for IPTG concentrations >50 μM, but on a linear scale. **B**, The average protein fluorescence in cells (both CheY-mCherry and mCherry expressing cells) of three different experiments (red, blue and green), as a function of IPTG induction levels. Datapoints were fitted with a second-degree polynomial expression (black). In one experiment (green) fluorescence values were markedly different from the two other experiments, and therefore it was excluded from the fit. **C**, The correlation between mCherry fluoresce and the concentration of the protein *in vivo*. This correlation was calculated by matching the curve in panel B with the linear part of the curve in panel A.

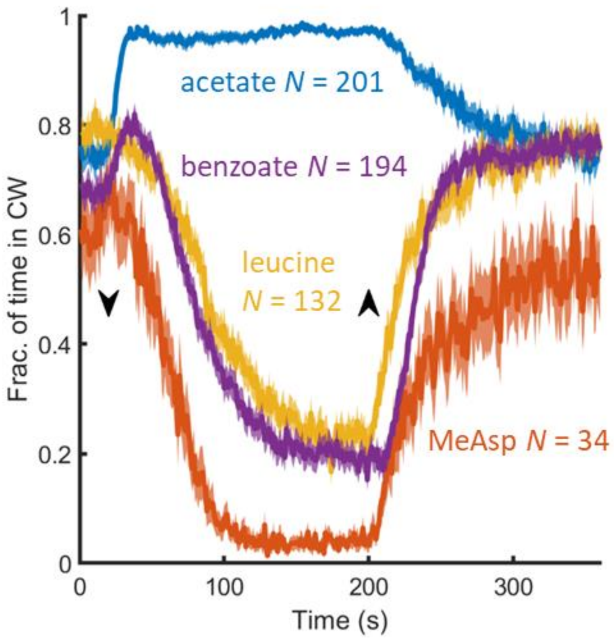

**Figure S4**. Mean response of $fliM_{\Delta N}$ cells expressing $FliM_N$-CheY (sPW417) to the introduction or removal of acetate, benzoate (each at 50 mM, pH 7.0), leucine and MeAsp (each at 1 mM). $FliM_N$-CheY expression was induced with 800 µM IPTG. Lines and shaded regions are mean ±SEM. *N* is the number of measured cells.

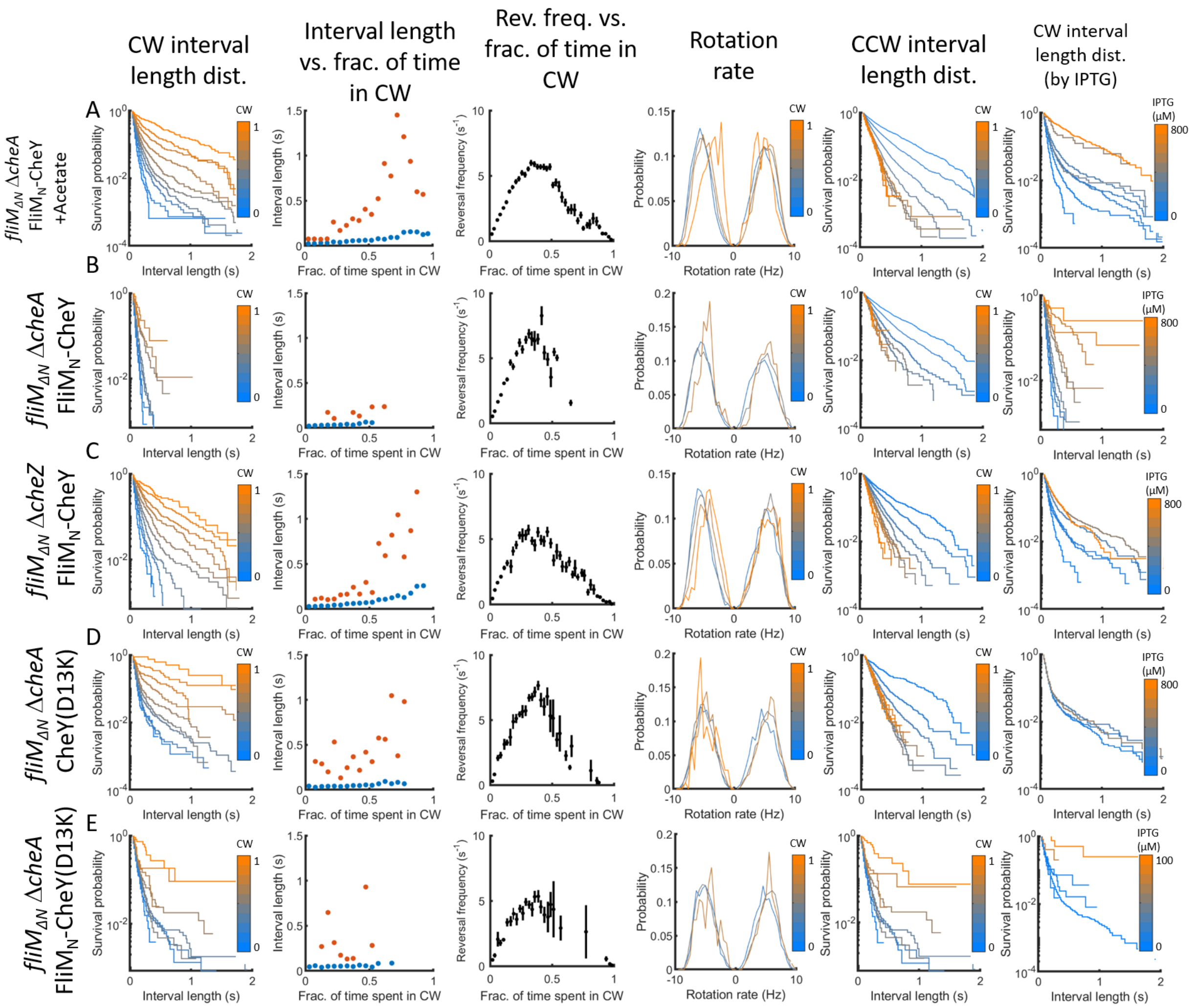

CW interval length dist.
Interval length vs. frac. of time in CW
Rev. freq. vs. frac. of time in CW
Rotation rate
CCW interval length dist.
CW interval length dist. (by IPTG)
A
fliMΔN ΔcheA FliMN-CheY +Acetate
B
fliMΔN ΔcheA FliMN-CheY
C
fliMΔN ΔcheZ FliMN-CheY
D
fliMΔN ΔcheA CheY(D13K)
E
fliMΔN ΔcheA FliMN-CheY(D13K)
Survival probability
Interval length (s)
Interval length (s)
Frac. of time spent in CW
Reversal frequency (s-1)
Probability
Rotation rate (Hz)
CW
IPTG (μM)
800
100

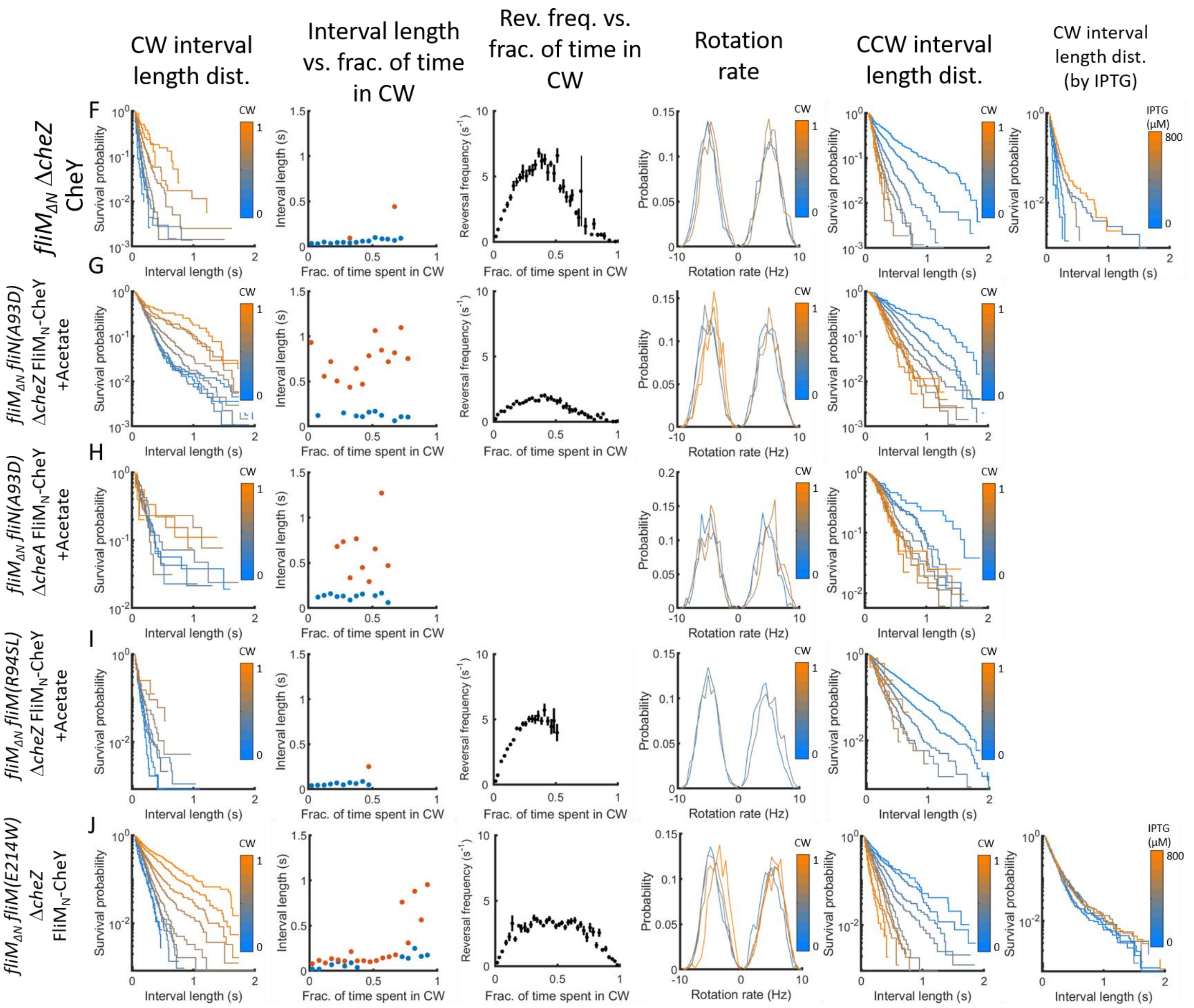

**Figure S5**. Rotation kinetics and velocity in most of the studied strains of this work. The results were pooled from all tethering experiments with the specified strains. **First column**, Distribution of clockwise interval lengths at 10 different clockwise levels (light blue is 0, orange is 1). **Second column**, Average duration of clockwise intervals contributed by the different switching phases as a function of the clockwise level. **Third column**, Average reversal frequency as a function of clockwise levels. **Fourth column,** Rotation rate, the number of a cycles covered by one second of clockwise or counterclockwise intervals (positive or negative rate, respectively). **Fifth column**, Distribution of counterclockwise interval lengths at 10 different clockwise levels (light blue is 0, orange is 1). **Sixth column,** Distribution of clockwise interval lengths from cells induced by different IPTG concentrations (light blue is 0, orange is 800 μM for all experiments except for J, where it is 100 μM). Note that the first and last columns differ from each other in the way that the experimental results are clustered: In the first one according to the clockwise level, in the last one according to the IPTG concentration employed for the induction of CheY expression. **A**, *fliM*$_{\Delta N}$ $\Delta$*cheA* cells expressing FliM$_N$-CheY in the presence of acetate (strain EW696). **B**, *fliM*$_{\Delta N}$ $\Delta$*cheA* cells expressing FliM$_N$-CheY (strain EW696). **C**, *fliM*$_{\Delta N}$ $\Delta$*cheZ* cells expressing FliM$_N$-CheY (strain EW697). **D**, *fliM*$_{\Delta N}$ $\Delta$*cheA* cells expressing CheY(D13K) (strain EW737). **E**, *fliM*$_{\Delta N}$ $\Delta$*cheA* cells expressing FliM$_N$-CheY(D13K) (strain EW738). **F**, *fliM*$_{\Delta N}$ $\Delta$*cheZ* cells expressing CheY (strain EW694). **G**, *fliM*$_{\Delta N}$ *fliN(A93D)* $\Delta$*cheZ* cells expressing FliM$_N$-CheY in the presence of acetate (strain EW714). **H**, *fliM*$_{\Delta N}$ *fliN(A93D)* $\Delta$*cheA*

cells expressing FliM$_{N}$-CheY in the presence of acetate (strain EW713). **I**, *fliM$_{\Delta N}$ fliM(R94SL)* Δ*cheZ* cells expressing FliM$_{N}$-CheY in the presence of acetate (strains EW731 and EW733). **J**, *fliM$_{\Delta N}$ fliM(E214W)* Δ*cheZ* cells expressing FliM$_{N}$-CheY (strain EW718).

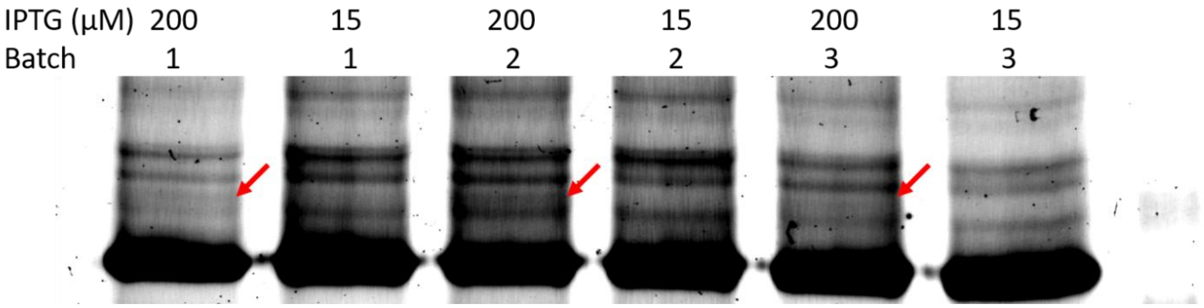

**Figure S6**. In-gel fluorescence of FliM$_{\Delta N}$-YPet after *in vivo* crosslinking by glutaraldehyde. *fliM$_{\Delta N}$-YPet* Δ*cheZ* cells overexpressing CheY (strain EW694) grown to O.D.$_{600nm}$=1 were washed 3 times in NaPi (10 mM, pH 7.6) and crosslinked with 0.001% glutaraldehyde for 1 h at room temperature. The reaction was quenched with 0.8 M Tris-HCl (pH 8.0). Following sonication, the cell lysate was resolved on 12% or 15% SDS-PAGE. The gel was visualized for YPet fluorescence. The image shows three independent repetitions of the same experiment, resolved on the same gel. The whole image was adjusted for contrast. Red arrows mark the crosslinked product.

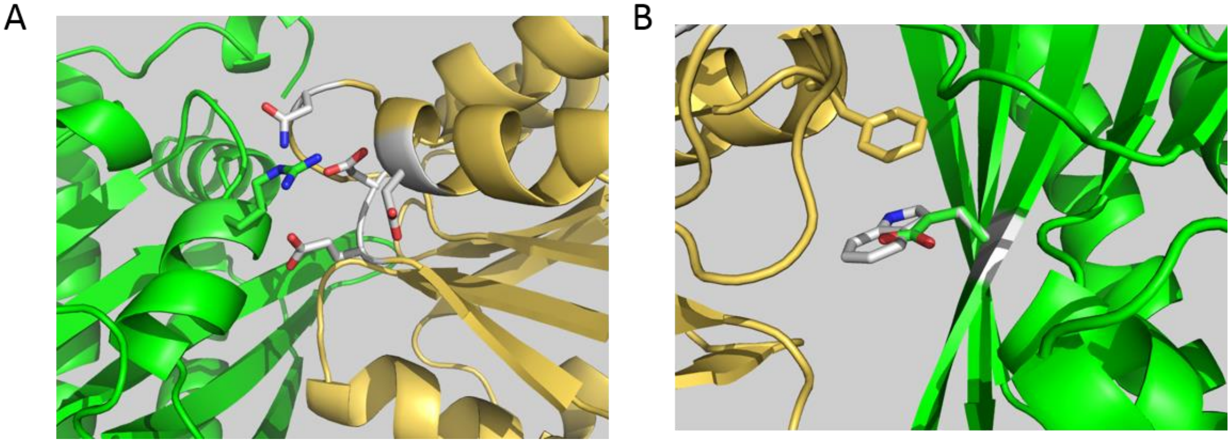

**Figure S7**. Predicted interfaces of CheY with FliM$_{M}$. **A**, An illustration of the predicted interaction of FliM(R94) with the acidic pocket in CheY. FliM is in green. K94 of FliM is shown as sticks. CheY backbone is in beige. Acidic residues in CheY are shown as gray sticks. Red and blue denote oxygen and nitrogen atoms, respectively. **B**, Molecular representation of the FliM(E214W) mutation site. FliM$_{M}$ is in green. The mutated Asp residue in FliM is shown in green sticks, the mutation in Trp is shown in white sticks. CheY is drawn in beige. Other colors are as in A. Note that the Asp to Trp mutation is expected to increase the hydrophobic interaction of FliM with F14 of CheY (drawn as sticks).

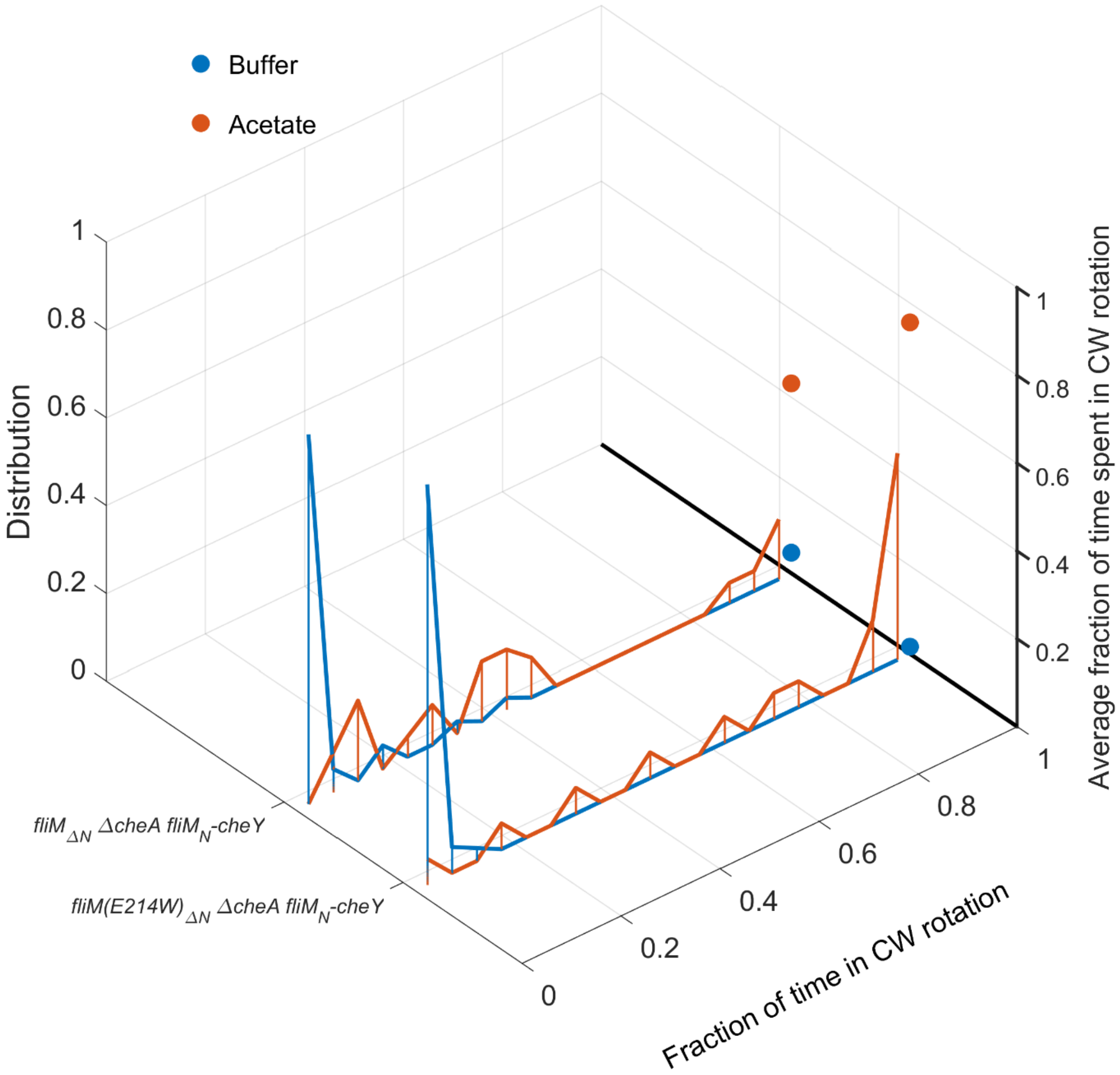

**Figure S8**. E214W mutation in *fliM*$_{\Delta N}$ does not affect the intrinsic motor bias. Cells deleted from the genome for *fliM*, *fliN* and *cheA* and expressing from a plasmid either FliM$_{\Delta N}$ FliN or FliM(E214W)$_{\Delta N}$ FliN (strains EW709 and EW717, respectively), were induced with 0.8 mM IPTG for FliM$_N$-CheY expression, and their direction of rotation was measured. The figure shows a normalized histogram of the measured cells. Dots are the mean of the distribution.

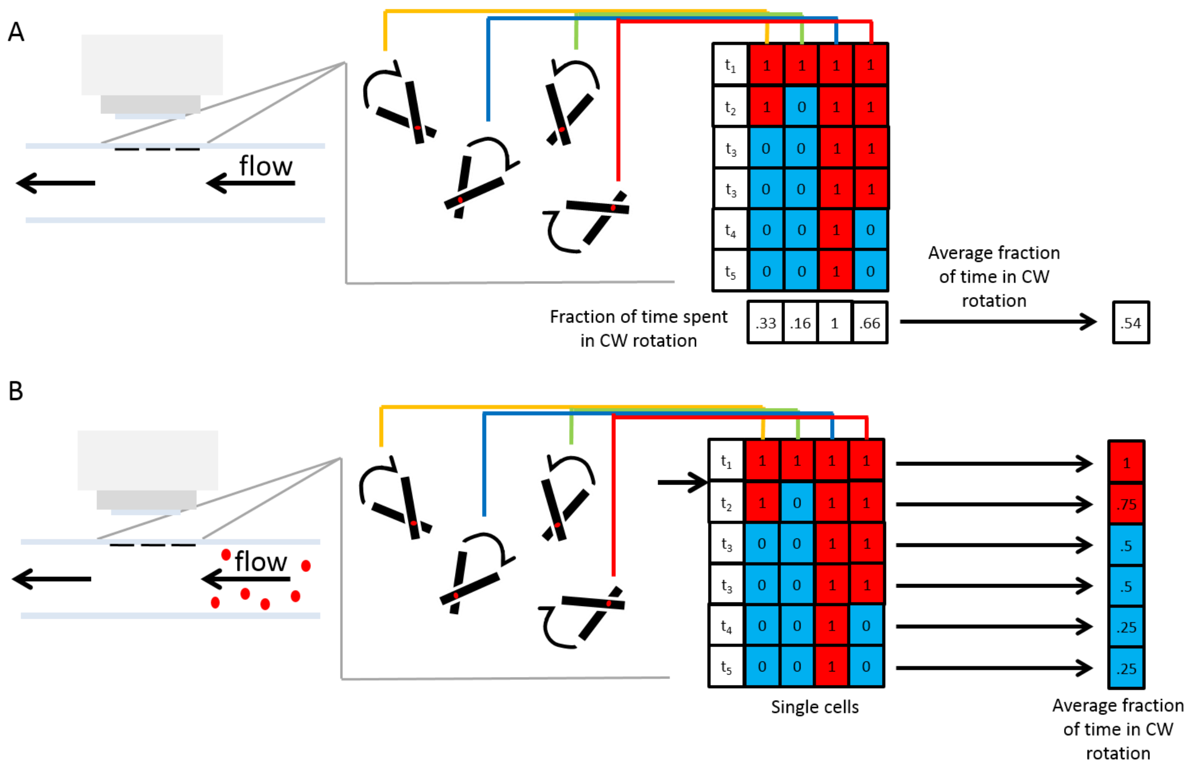

**Figure S9.** Calculation of the fraction of time spent in clockwise rotation. **A**, Measurement of the average clockwise (CW) rotation in non-stimulated cells. On the left, a schematic presentation of a flow chamber. The rotation of multiple cells is measured over time (as illustrated in the middle). The level of clockwise rotation is measured for each cell individually and a grand average is calculated from all observed cells. **B**, Measurement of the average clockwise rotation in response to attractant stimulation. On the left, a schematic presentation of a flow chamber, red circles mark an incoming attractant. The rotation of multiple cells is measured over time (as illustrated in the middle). On the right, a table shows the direction of angular displacement of the measured cells. The arrow marks the time at which the attractant arrived to the cells. The probability of clockwise rotation is averaged over small-time intervals for all cells (usually over 1 s; far right).

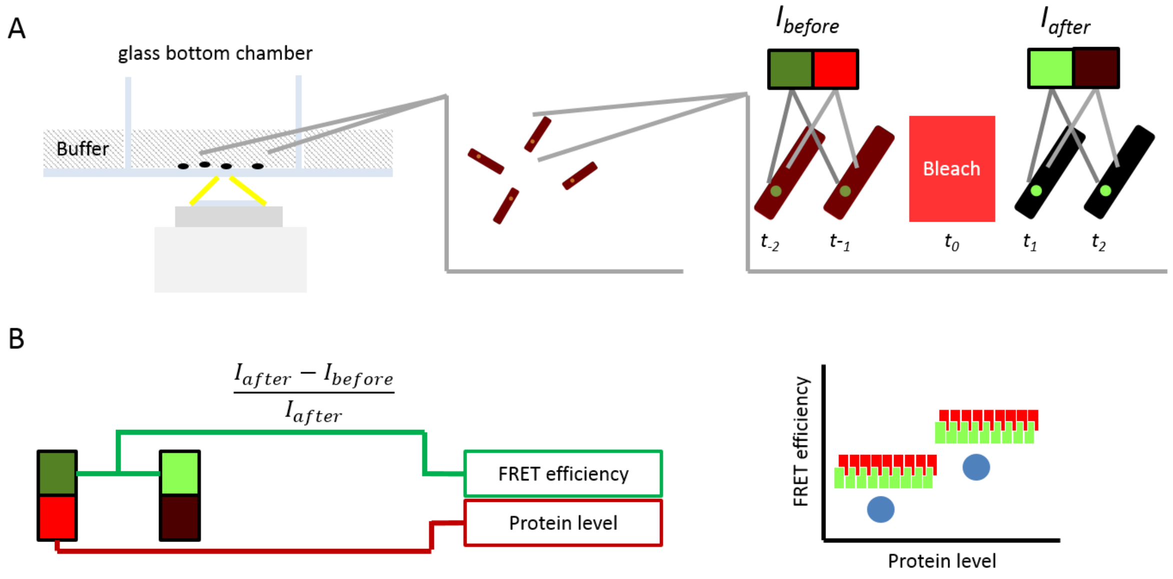

**Figure S10.** FRET, experimental setup and analysis*.* **A**, Experimental setup. On the left, cells, illustrated as black dots, were attached to a glass bottom chamber by incubation with poly-L-Lysine, and observed by an inverted fluorescent confocal microscope. In the middle are the same cells from the objective point of view. On the right is an illustration of a FRET measurement in a single cell. Green dot marks the motor fluorescence. The reddish color of the cell marks CheY fluorescence. Colored pair of rectangular shapes show the average fluorescence of the motor (green) or CheY (red), before (<$t_0$) or after (>$t_0$) bleaching. **B**, FRET analysis. For each measured cell, the recorded CheY fluorescence prior to bleaching was paired with the FRET efficiency (calculated from the motor fluorescence before and after bleaching). The pair of values were sorted by the mCherry fluorescence and divided to groups of 100 sorted data points.

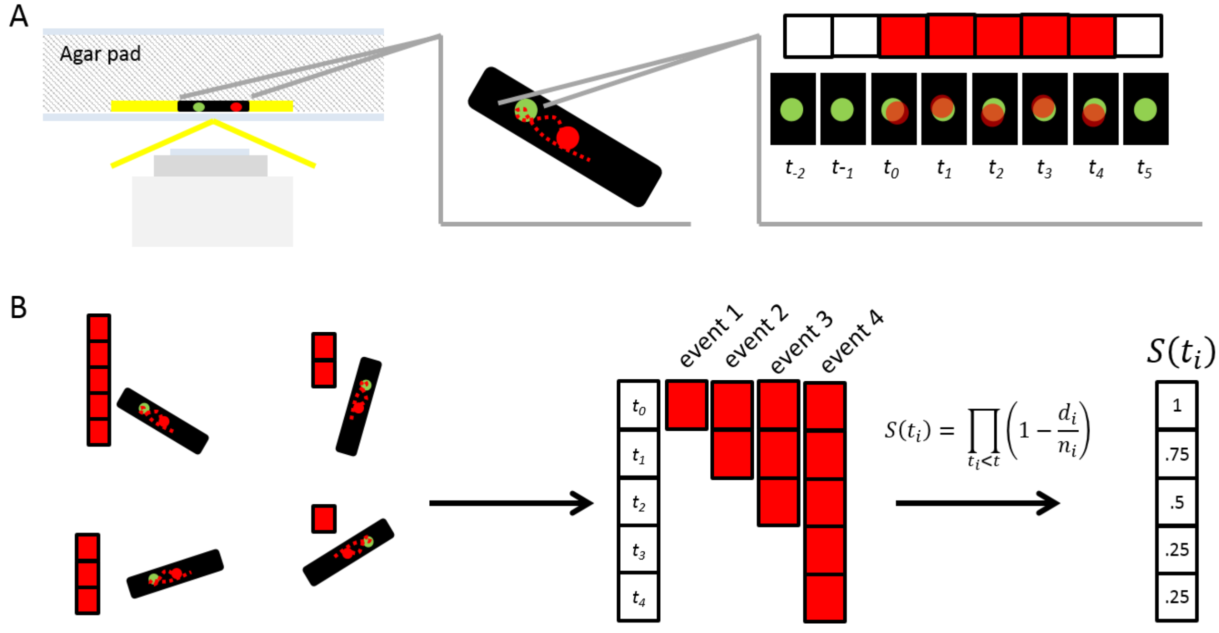

**Figure S11.** Single molecule observation, experimental setup and analysis. **A**, Experimental setup. On the left, a cell, illustrated as a black rectangle, was deposited onto an agar pad and visualized by an inverted microscope using HILO-TIRF illumination. In the middle, the same cell is seen from the objective point of view. A green dot marks the fluorescence of the motor. A red dot marks CheY. Dotted trajectory illustrates the diffusion of the CheY molecule in time. On the right is an illustration of a measurement of the dwell of CheY with the motor. Each black rectangular shape with a green dot in its middle illustrates the motor and CheY fluorescence in time. The row above it illustrates for each point of time whether the motor was vacant (white fill) or CheY bound (red fill). **B**, Survival analysis. On the left, different cells were measured for different dwell times of CheY near the motor. In the middle, these dwell time values were collected from all cells, sorted and subjected to survival analysis. The column on the right shows the calculated survival values.

#### SUPPLEMENTARY MOVIES

**Movie S1**. Demonstration of small clockwise intervals in the rotation of a tethered $FliM_{\Delta N}$ containing motor (strain EW697). Cell rotation was documented at ~140 frames/s. Two clockwise intervals are observable in the movie, one spans for ~28 ms (starting at 7.34 s) and the other spans for ~64 ms (starting at 7.88 s). The image shows the position of the tethered cell. Up-right panel shows the angle of the cell as a function of time and the down-right panel shows the rotation rate as a function of time. The blue marker shows the time point which corresponds to the current image.

**Movie S2**. Single-molecule tracking of CheY in a single cell. The movie shows a single *fliM*-yPET Δ*cheZ* cell (strain EW669), electroporated with CheY(I95V)-Atto647 molecules and documented after exposure to 50 mM acetate. White circles stand for 75 nm radius around the estimated FliM-YPet locations. White dots show the calculated location of CheY(I95V)-Atto647 molecules. Each frame in the movie shows the location data overlaid on top of both the average of the first ten images observed via the green channel (showing the fluorescence of FliM-YPet; left panel) and the time-lapse images taken in the red channel (fluorescence of internalized CheY(I95V)-Atto647; right panel). CheY(I95V)-Atto647 and FliM-YPet binding intervals are considered as events in which CheY(I95V)-Atto647 dwelled within the 75 nm radius of FliM. For clarity, the movie is slowed down from 100 to 20 frames/s.
